# Selective phase separation of transcription factors is driven by orthogonal molecular grammar

**DOI:** 10.1101/2024.04.12.589262

**Authors:** Mark D. Driver, Patrick R. Onck

**Author notes:** Corresponding author.; Contributing author.

## Abstract

Protein production is critically dependent on gene transcription rates, which are regulated by RNA polymerase and a large collection of transcription factors (TFs). Previous studies identified the formation of super enhancer regions where increased transcriptional activity is observed. This has been linked to phase separation, in which the differential condensation behaviour of separate TF families has been hypothesised to cause the selectivity in gene expression. The underlying molecular forces that are responsible for this selectivity, however, are unknown. Here, we conduct phase separation studies on six TFs (FUS, EWS, TAF15, SP1, SP2, and HNF1A) from three different TF families by carrying out residue-scale coarse-grained molecular dynamics simulations. Our exploration of ternary TF phase diagrams revealed four dominant sticker motifs and two orthogonal driving forces, consisting of hydrophobic (aromatic, aliphatic) interactions and electrostatic/cation-***π*** interactions. The contribution of these driving forces to the homotypic and heterotypic intermolecular strengths dictate the resultant condensate morphology. These results point to sequence-dependent orthogonal grammar as a generic mechanism responsible for selective transcriptional condensation in gene expression. Interestingly, our results also show how RNA polymerase is able to overcome this orthogonality to coalesce with TF condensate providing a framework in which co-condensation is the primary nuclear organisational principle that controls selective partitioning of RNAP to targeted genes for transcription.

## 1 Introduction

Transcription is the first step in extracting the information stored in DNA for RNA production [1, 2]. RNA polymerase (RNAP), the enzyme responsible for RNA production, is a multi-subunit protein that must be directed to the correct gene for RNA production to start [1–3]. This function is performed by an auxiliary network of transcription factors (TFs), which localises the required RNAP at the correct gene on the DNA to initiate transcription [2–9], copying the genetic information to RNA. The transcription rates of different genes are controlled through transcriptional (or gene) regulatory networks (TRNs) [10] by the recognition of specific promoter regions on DNA by the TFs, which are located near the targeted gene in space [4, 11, 12]. Independent gene regulation requires multiple differential transcription activation pathways within a TRN to enable selective gene expression to ensure that the correct genes are transcribed by the associated TF [13–16].

In recent years it has been discovered that gene transcription is not only a biochemical process; spatial localization also plays a key role. Super-enhancer regions spatially contribute to increased expression of specific genes, with down regulation of other genes [17–20]. The formation of such super enhancer regions and the associated transcriptional regulation has been increasingly linked to liquid-liquid phase separation (PS) [13, 17, 19–28]. PS has been identified in the recent decade to drive the formation of membraneless organelles in several areas of cell biology [29–31]. Phase separation is driven by multivalent interactions of low complexity domains (LCDs) [19, 29, 32–36] which are common features of transcription factors, while the RNAP C terminal domain region also contains a LCD [37, 38]. The interaction of these LCDs is important for RNAP binding, with the size of the RNAP C terminal domain modulating the uptake into condensates mediating the transcription rates [38–40]. Disruption and misregulation of TF condensates are oncogenic drivers, leading to promotion of aberrant gene transcription behaviour [41, 42].

Understanding the driving forces for TF condensate formation is therefore essential to understand how gene expression can be regulated, and how it can be disrupted in the case of disease. In this work we aim to identify the molecular interactions that drive selective condensate formation, by focusing on a range of transcription factors from three different transcription families (FET, SP/KLF and HNF). It has been hypothesized that the selectivity of TFs is caused by differential phase separation propensities [22, 24, 43, 44]. Here we test this hypothesis by analyzing ternary phase diagrams of six TFs that all undergo PS to form condensates on their own under biological conditions [22, 24, 45].

We use coarse-grained molecular dynamics simulations at amino-acid resolution to determine the molecular interactions that drive TF selectivity. Results are presented in terms of ternary phase diagrams and residue-scale contact maps. These revealed two distinct driving forces for condensate formation, consisting of hydrophobic interactions (from aromatic and aliphatic residues) and electrostatic/cation-*π* interactions (from cationic residues interacting with anionic or aromatic residues). We identified four dominant amino acid types, i.e. aromatic, aliphatic, cationic and anionic residues, responsible for driving collective phase separation forming four orthogonal sticker motifs in the conceptual ‘sticker-spacer’ model of polymers [36]. The sequence composition of the TFs enables control over the homotypic and heterotypic intermolecular interactions, and therefore condensate selectivity. The relative homotypic and heterotypic strengths result in four distinct droplet morphologies: marbled, coated, bimodal, or separated droplets (heterotypic interactions are significantly weaker than homotypic interactions). The ability of the LCD of RNAP (POL II) to interact with condensates of all TFs highlights its unique central position and points to a new universal principle in which selective transcription is regulated by sequence-based homotypic and heterotypic phase separation.

## 2 Results

### 2.1 Selection of transcription factors

Droplet simulations were undertaken of six TFs from three different TF families: the FET family (FUS, EWS, TAF15), the SP/KLF family (SP1, and SP2), and the HNF family (HNF1A) to explore the orthogonal driving forces for phase separation (see the amino acid sequences in Fig. 1). We examine a large range of relative compositions of the ternary compositional space to fully describe the key motifs responsible for selective partitioning. To understand how the balance of homotypic and heterotypic interaction strengths varies for different members within the same family or with a different family, simulations of the seven key points on new phase diagrams for different ternary combinations were conducted as shown in Table 1. Finally we conducted a study of TF-POLII condensate interactions. Simulations were done at a monovalent ion concentration of 150 mM, with the maximum number of molecules set to 120 for FUS, EWS, and TAF15, 60 for SP1, SP2, and POLII, and 90 for HNF1A, so that the maximum number of amino acids per molecule type is approximately the same. PS is observed across all compositions of the TFs and POLII studied.

**Fig. 1.**
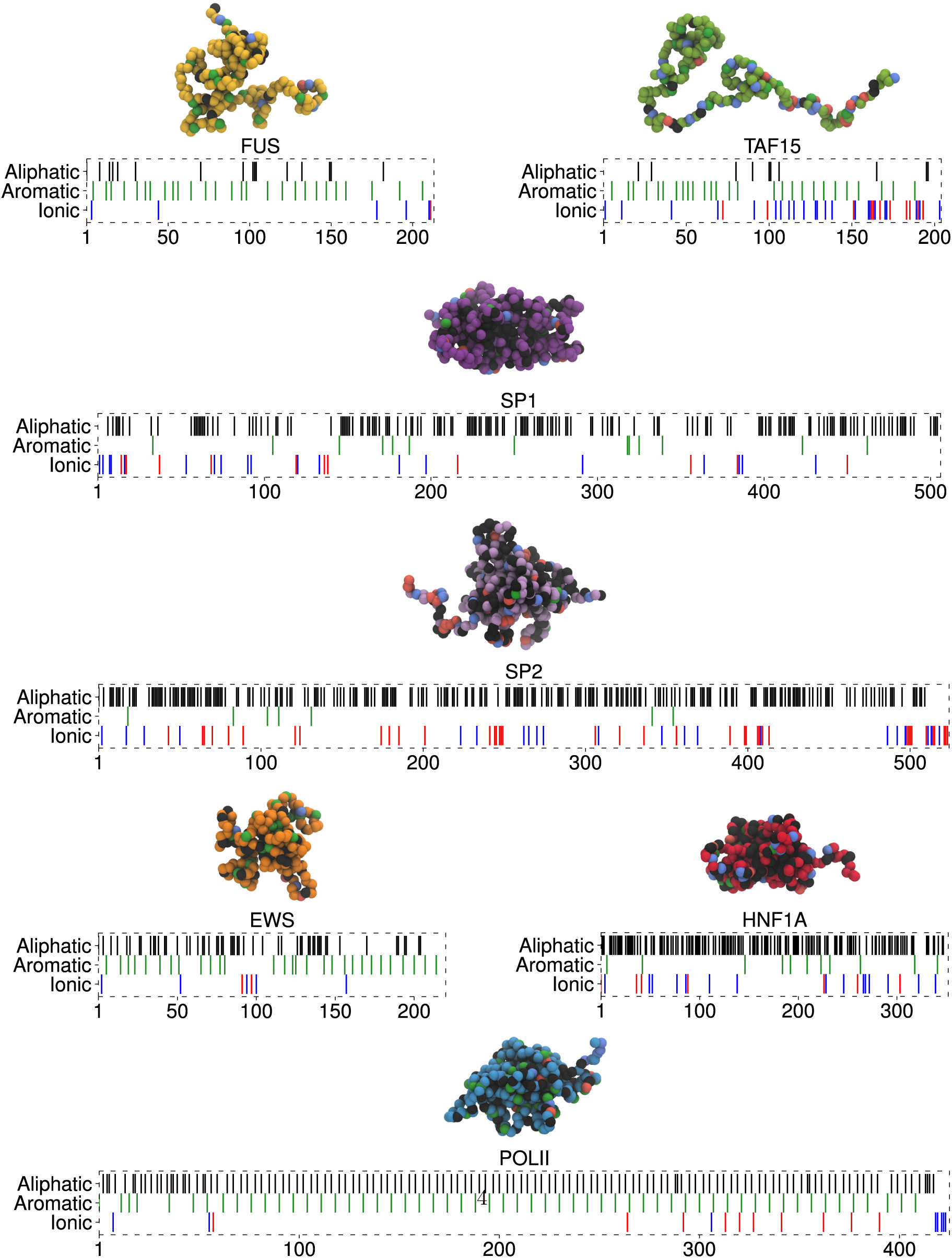
Coarse-grained representation and residue composition of the TFs studied and POL II. A) FUS, B) TAF15, C) SP1, D) SP2, E) EWS, F) HNF1A, G) POL II. Residues are categorised into 5 groups: cations (R, K)-red, anions (D, E)-blue, aromatic (F, Y, W)-green, aliphatic (A, C, I, L, M, P, V)-black, hydropillic (G, N, S, H, Q, T)-white. The hydrophillic residues in the 1 bead-per-residue molecular representation are coloured differently to distinguish the proteins studied in this work: FUS (yellow), TAF15 (green), SP1 (purple), SP2 (light purple), EWS (orange), HNF1A (red), POL II (blue).

**Table 1.**
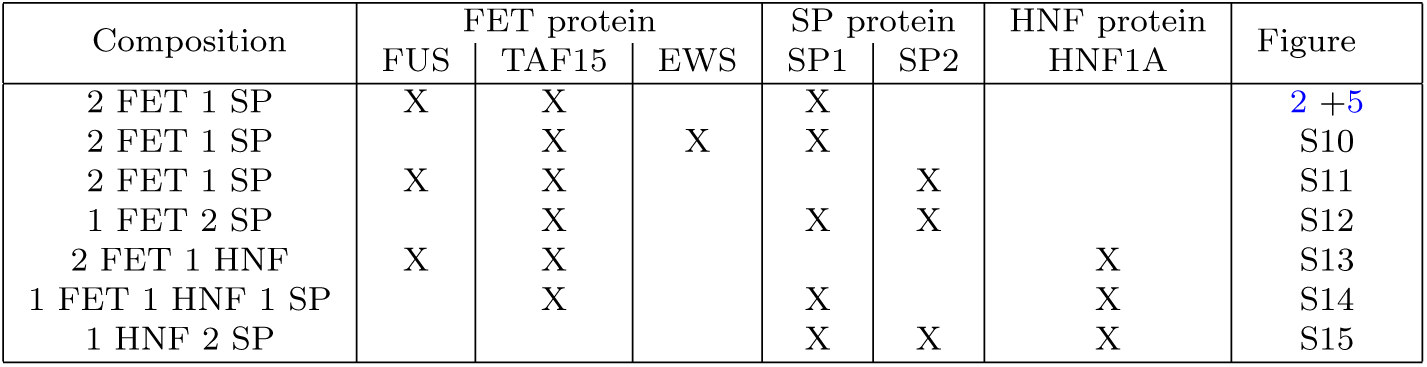
Summary of transcription condensate systems studied by molecular composition and the corresponding figures with simulation images. Sequence information can be found in Fig. 1.

| Composition | FET protein |  |  | SP protein |  | HNF protein<br>HNF1A | Figure |
| --- | --- | --- | --- | --- | --- | --- | --- |
|  | FUS | TAF15 | EWS | SP1 | SP2 |  |  |
| 2 FET 1 SP | X | X |  | X |  |  | 2 + 5 |
| 2 FET 1 SP |  | X | X | X |  |  | S10 |
| 2 FET 1 SP | X | X |  |  | X |  | S11 |
| 1 FET 2 SP |  | X |  | X | X |  | S12 |
| 2 FET 1 HNF | X | X |  |  |  | X | S13 |
| 1 FET 1 HNF 1 SP |  | X |  | X |  | X | S14 |
| 1 HNF 2 SP |  |  |  | X | X | X | S15 |

In the next section we start out by analyzing the FUS-SP1-TAF15 in detail as a reference system that has been studied experimentally [24].

### 2.2 Aromatic, aliphatic and cation-*π* interactions drive simple coacervation of TFs in the FUS-SP1-TAF15 system

Pure FUS, SP1 and TAF15 all exhibited strong homotypic phase separation forming stable droplets, as shown at the corners of the ternary phase diagram of Fig. 2. The molecular interactions driving PS can be determined by examining the molecular contacts between the amino-acids for the single component systems, shown in Fig. 3. Fig. 3A and D show the contact maps for FUS, which indicate that the hydrophobic *π − π* interactions between tyrosine residues are the main driving force for PS. For FUS the 1D summations of interactions (Fig. 3A) show increased contacts around the locations of aromatic residues, highlighted with black dashed lines. This is clearly indicative of the ‘sticker-spacer’ model [36] with the tyrosine residues acting as stickers that drive phase separation. Interactions with and between the glycine, serine, and glutamine are also present in FUS, with a significant decrease in interactions around anionic residues, due to the repulsive electrostatic nature, leading to white streaks on the contact map. The contact maps for the SP1 simulation (Fig. 3B and E), show a completely different interaction profile: here aliphatic hydrophobic interactions are key contributors to PS due to the high abundance of these residues in SP1, that is also reflected in the 1D summations of the interactions. The SP1 N-terminus has much lower contact frequencies due to a greater density of ionic residues, which have repulsive interactions with the aliphatic residues that constitute the majority of the molecules. This arises from the hydrophilic nature of the ionic residues, making the interactions with neutral hydrophobic aliphatic residues repulsive. In SP1 alanine (A), isoleucine (I), leucine (L), proline (P) and valine (V) are the aliphatic residues promoting PS.

**Fig. 2.**
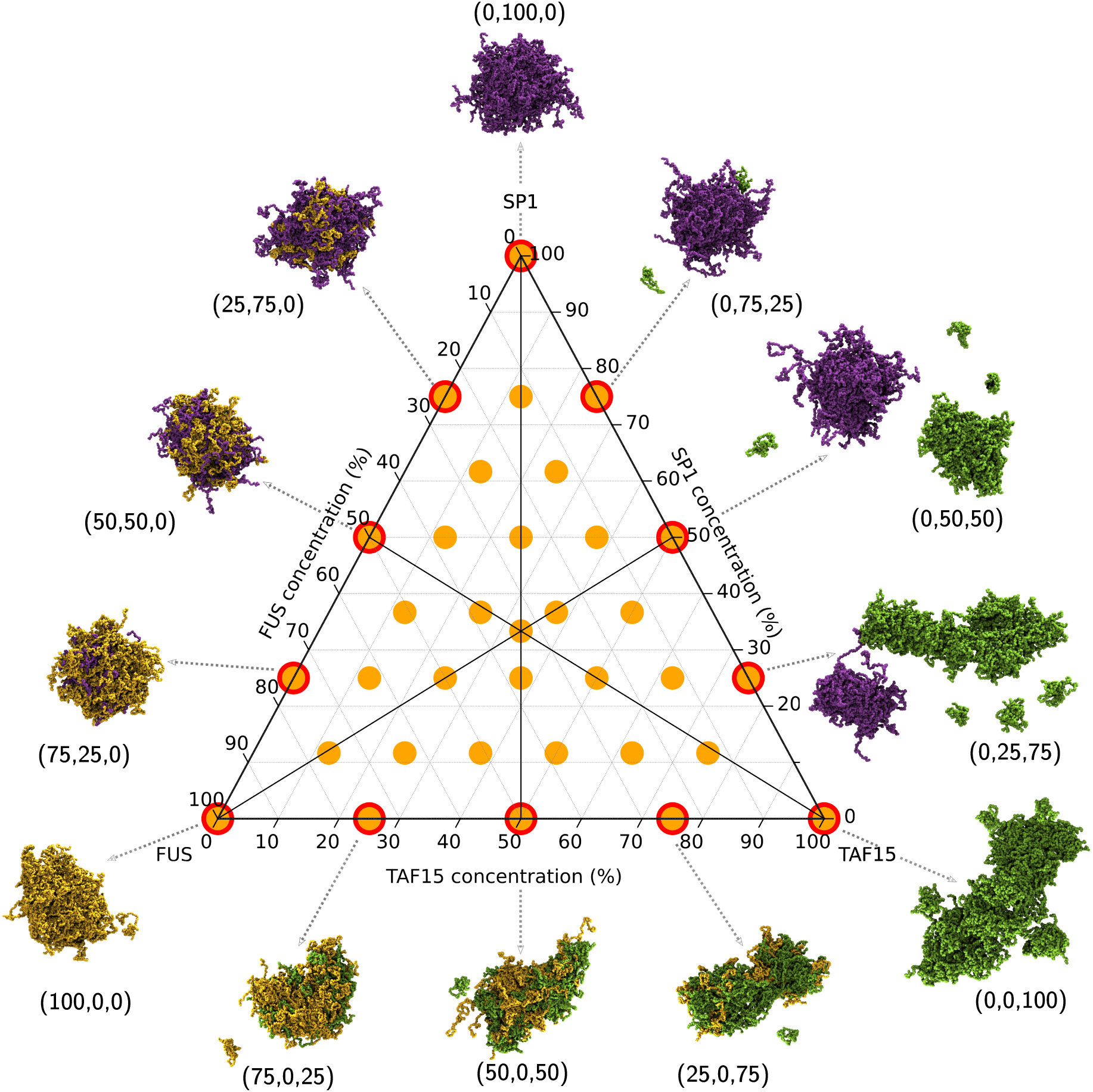
Ternary phase diagram of the three TFs with snapshots of the two-component droplets along the edge of the diagram. Ternary phase diagram showing the simulations undertaken in this work (orange dots) run with a total amino acid concentration of 80000 *µM*. The total composition is defined relative to 120 FUS molecules, 120 TAF15 molecules, and 60 SP1 molecules, to give the percentage compositions (% FUS, % SP1, % TAF15). The end frame of 3 *µ*s of simulation is displayed as a representative state. FUS molecules are coloured in yellow, SP1 molecules are coloured in purple, and TAF15 molecules are coloured in green. Droplets are formed in all simulations of these molecules under these concentration conditions, irrespective of composition.

**Fig. 3.**
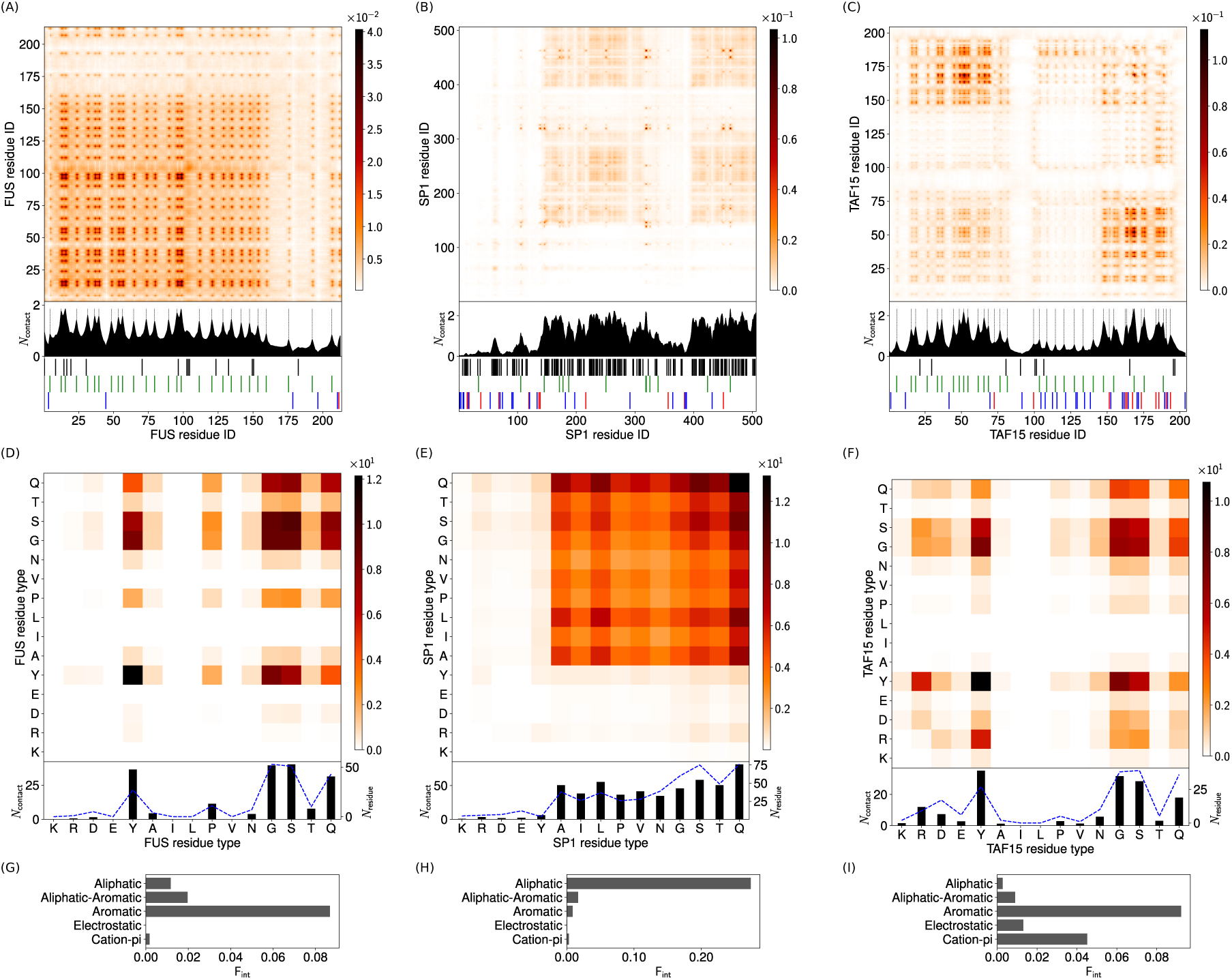
Intermolecular contact maps for homotypic interactions in single-component droplets. (A)-(C) Intermolecular contacts by residue index for (A) 100% FUS, (B) 100% SP1, and (C) 100% TAF15, at 150 mM ion concentration and 300 K. The contacts are averaged in time and normalised by the number of molecules in the simulation (see section 3.2 in the SI for more details). A 1D contact profile (summation of the 2D map) is included below the contact map to show the total interactions per residue index (*N*_contact_). The black dashed lines in (A) and (C) highlight the residues with the most contacts in FUS and TAF15 (which correspond to peaks in the 1D profiles). These are the aromatic residues in FUS driving *π*-*π* aromatic contacts, and the aromatic and cationic residues in TAF15 driving aromatic and cation-*π* interactions. Broad peaks are seen in (B) corresponding to the aliphatic residue stretches in SP1. (D)-(F) Intermolecular contact map by residue type for (D) 100% FUS, (E) 100% SP1, and (F) 100% TAF15. The contact maps in (D), (E) and (F) are similar to the contact maps by residue index in (A), (B), and (C), respectively, but aggregated by residue type. A 1D contact profile (summation of the 2D map), are also included below (*N*_contact_) together with the abundance for the residues (*N*_residue_) shown by blue dashed lines. (G)-(I) Intermolecular interaction summary for (G) FUS in 100% FUS (H) SP1 in 100% SP1, and (I) TAF15 in 100% TAF15 at 150 mM and 300 K. The fraction of interactions, F_int_, are aggregated by type and normalised by the total number of the intermolecular interactions in (A)-(C) respectively. Aromatic and aliphatic interactions denote aromatic-aromatic and aliphatic-aliphatic interactions respectively. This convention is used throughout this work. Details of the contact definitions can be found in section 3.2 of the SI.

In contrast to the purely hydrophobic driving forces for FUS and SP1, TAF15 PS has significant contributions from cation-*π* interactions in addition to the dominant aromatic contacts as can be seen in the contact maps in Fig. 3C and F. For TAF15 the 1D summations of interactions show increased contacts around the locations of cationic and aromatic residues indicative of cation-*π* interactions (black dashed lines in Fig. 3C). The N-terminus has a large number of tyrosine residues, with a relatively low number of anionic residues. The C-terminus, on the other hand, contains the majority of the cationic residues, leading to the high number of cation-*π* (arginine-tyrosine) contacts between the opposite ends of two TAF15 proteins. The central region contains a greater concentration of anionic residues, leading to a decrease in contacts, because it is self-repulsive and also repels the N-terminus, leading to the broad white bands. Interactions between glycine, serine and glutamine residues within TAF15 also contribute to PS due to the high abundance of these residues, but they are significantly weaker than the arginine-tyrosine interactions (see Fig. S25C for the contact map normalised by residue abundance).

Fig. 3G-I shows that the most important interactions are aromatic interactions for FUS and TAF15, and aliphatic interactions for SP1. Cation-*π* interactions also play a substantial role for TAF15. It is striking to see that for FUS and TAF15, despite the large difference in spatial contact maps (Fig. 3A and C based on residue index), the nature of the molecular interactions (Fig. 3G and I), and the residue-based contact maps (Fig. 3D and F) are remarkably similar, with the appearance of additional arginine-tyrosine (cation-*π*) contacts for TAF15.

The locations of the sticker residues within the molecules have a distinct effect on the topology of the single component droplets. FUS has a relatively even distribution of stickers throughout the protein allowing for the formation of relatively spherical droplets. TAF15 has a bimodal distribution of stickers located at the two termini, and a repulsive spacer region in the centre that repels other chains. This leads TAF15 to form a more open and porous structure where the droplet is formed of a collection of associating smaller TAF15 clusters (Fig. 2). Increasing the system size (increasing the number of copies of molecules) does not change this behaviour (Fig. S8), indicating that finite size effects do not play a role here. In all cases highly dynamic contacts are seen with contact lifetimes in the range 4-6 ns (Fig. S9), showing that highly dynamical cross links are present [46, 47].

### 2.3 Heterotypic interactions compete with homotypic interactions

As previously mentioned, PS is observed for all compositions, but the droplet morphologies and the molecular interactions change in nature as a second component was introduced, shown in Fig. 2. FUS-TAF15 mixtures form droplets with both proteins mixed and uniformly distributed throughout the condensate, with the droplet becoming increasingly less spherical with increasing TAF15 fraction. Whereas for FUS-SP1 mixtures the condensate is more spherical, with FUS-SP1 being well mixed at high SP1 fractions, before FUS starts to form a coating on the surface of an SP1 droplet at high FUS fractions. Interestingly, for TAF15-SP1 mixtures the condensate morphology is completely different. A singular mixed droplet is not favoured, instead the molecules are partitioned into two condensates: a TAF15 condensate and a SP1 condensate which do not interact.

Before examining the contact maps to clarify this, it is important to note that the intermolecular contact maps for the homotypic interactions of FUS, TAF15 and SP1 shown in Fig. 3 for the single component droplets are identical to the homotypic intermolecular contact maps observed in mixed systems (see Figs. S23-S30 in the SI). In this section we therefore exclusively focus on the heterotypic interactions between the two components in a system.

FUS-SP1 intermolecular interactions, shown in Fig. 4A and D, are primarily hydrophobic interactions driven by the large number of aliphatic residues in SP1 and tyrosine residues in FUS. The cation-*π* interactions appearing sharply in Fig. 4A are only a small part of the overall FUS-SP1 interactions with aliphatic-aromatic interactions the main FUS-SP1 intermolecular interactions (Fig. 4D and G). FUS-TAF15 intermolecular interactions, shown in Fig. 4B are driven by aromatic interactions between tyrosine residues in FUS and TAF15. Additional cation-pi contacts, from cations in TAF15 interacting with the tyrosines in FUS are also observed (Fig. 4E and H). SP1-TAF15 intermolecular interactions are minimal, shown in Fig. 4C by the 1D contact summaries being two orders of magnitude smaller than for FUS-SP1 or FUS-TAF15 contacts in Fig. 4A and B. The large number of anionic residues in the centre and C-terminal of TAF15 (TAF15 has a relatively high net negative charge per residue of −12/204) only offer repulsive interactions with the large number of aliphatic and anionic (net charge per residue of −8/506) residues in SP1. This restricts favourable SP1-TAF15 interactions to much smaller chain segments, around cations in SP1, and the formation of very few contacts. The SP1-TAF15 interactions are much more localised than between SP1 and FUS, or TAF15 and FUS, or the homotypic interactions, making any mixing of the two molecules in one condensate unfavourable. By looking at the residue-based contact maps in Fig. 4D-F, the stickers and spacers of FUS and TAF15 are strikingly similar to each other and rather different from that of SP1, clearly highlighting that they originate from two different families of TFs.

**Fig. 4.**
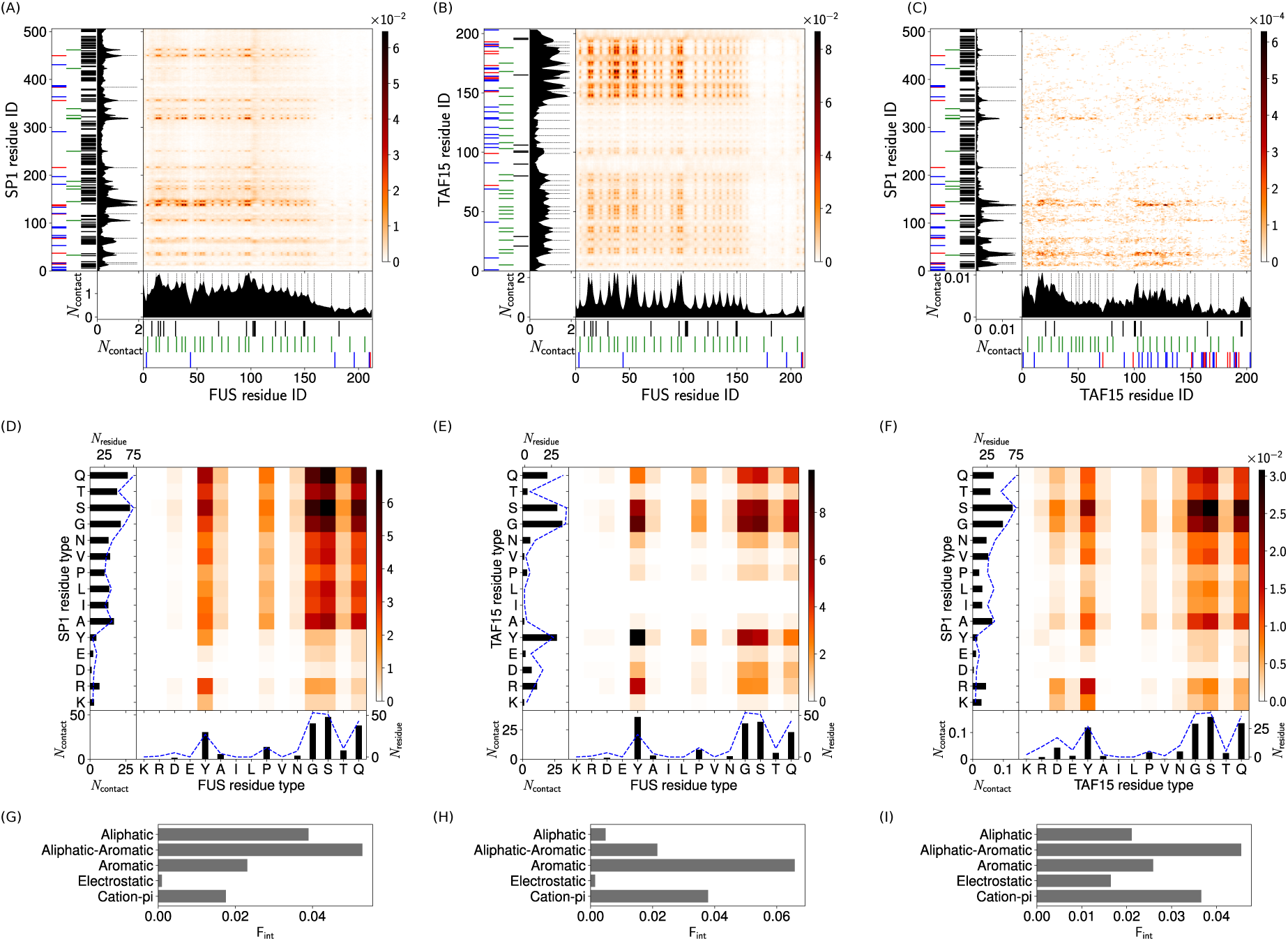
Intermolecular contact maps for heterotypic interactions in two-component droplets. (A)-(C) Intermolecular contact map by residue index for (A) FUS with SP1 (50%, 50%), (B) FUS with TAF15 (50%, 50%), and (C) SP1 with TAF15 (50%, 50%) at 150 mM and 300 K. For the definitions of the different contact types see the caption of Fig. 3 and section 3.2 of the SI. The 1D contact profiles denote a summation of the 2D map of the corresponding molecules. The black dashed lines highlight the key residues: (A) the aromatic residues in FUS with cationic and aromatic residues in SP1, (B) aromatic residues in FUS with the aromatic and cationic residues in TAF15, and (C) the cationic residues in SP1 with the aromatic residues in TAF15. (D)-(F) Intermolecular contact map by residue type for (D) FUS with SP1 (50%, 50%), (E) FUS with TAF15 (50%, 50%), and (F) SP1 with TAF15 (50%, 50%) at 150 mM and 300 K. (G)-(I) Intermolecular interaction summary for (G) FUS-SP1 interactions in (50%, 50%, 0%), (H) FUS-TAF15 interactions in (50%, 0%, 50%), and (I) SP1-TAF15 interactions in (0%, 50%, 50%) at 150 mM and 300 K. The fraction of interactions, F_int_, are aggregated by type and normalised by the total number of the intermolecular interactions in (A)-(C) respectively.

The normalised number of interactions between different species provides a method for comparison (full data for all simulations can be found in Tables S9-S13 and Figs. S3-6 in the SI). The single component mixtures all have comparable numbers of homotypic interactions for FUS (0.015), SP1 (0.016), and TAF15 (0.014) with TAF15 forming slightly more open condensates. In the two component mixtures we see the emergence of a difference in the relative strengths of the homotypic and heterotypic interactions. Three possible cases exist for binary mixtures. In the first case the heterotypic interactions are stronger than the homotypic interactions (FUS-TAF15), then mixing of the two species in a marbled condensate is preferred (see (75,0,25) in Fig. 2 and Fig. S5B). In the second case when the homotypic interactions are both stronger than the heterotypic interactions (SP1-TAF15), then a bimodal/separated condensate system is preferred (see (0, 50, 50) in Fig. 2 and Fig. S5C). For the third case the homotypic and heterotypic interactions are of a comparable strength (FUS-SP1), leading to a coated (or core-shell) condensate structure (see (75, 25,0) in Fig.2 and Fig. S5A).

### 2.4 Ternary systems display complex condensate morphology driven by interaction orthogonality

Moving into the centre of the FUS-SP1-TAF15 phase diagram we now see the competition between the attractive groups in the different molecules for interactions in the resultant ternary condensates. A combination of the previously observed morphologies is now seen, as shown in Fig. 5. Attractive interactions of FUS with both SP1 and TAF15 are opposed by the repulsive interactions between SP1 and TAF15 in the formation of a single heterogeneous condensate. Competition between SP1 and TAF15 for the interactions with FUS leads to a complex, composition-dependent behaviour of the droplets. In high FUS fraction compositions a single droplet is observed with SP1 and TAF15 localised at different ends of the condensate such that SP1-TAF15 contacts are minimised. At low FUS fractions a singular large condensate is lost, and instead separate droplets are formed containing only TAF15 with FUS inclusions or SP1 with FUS inclusions, see e.g., the (25, 50, 25) and (12.5, 12.5, 75) compositions. This transition to separate droplets occurs at 50% SP1 for SP1 rich systems, whereas TAF15 rich systems require 75% TAF15 for analogous behaviour.

**Fig. 5.**
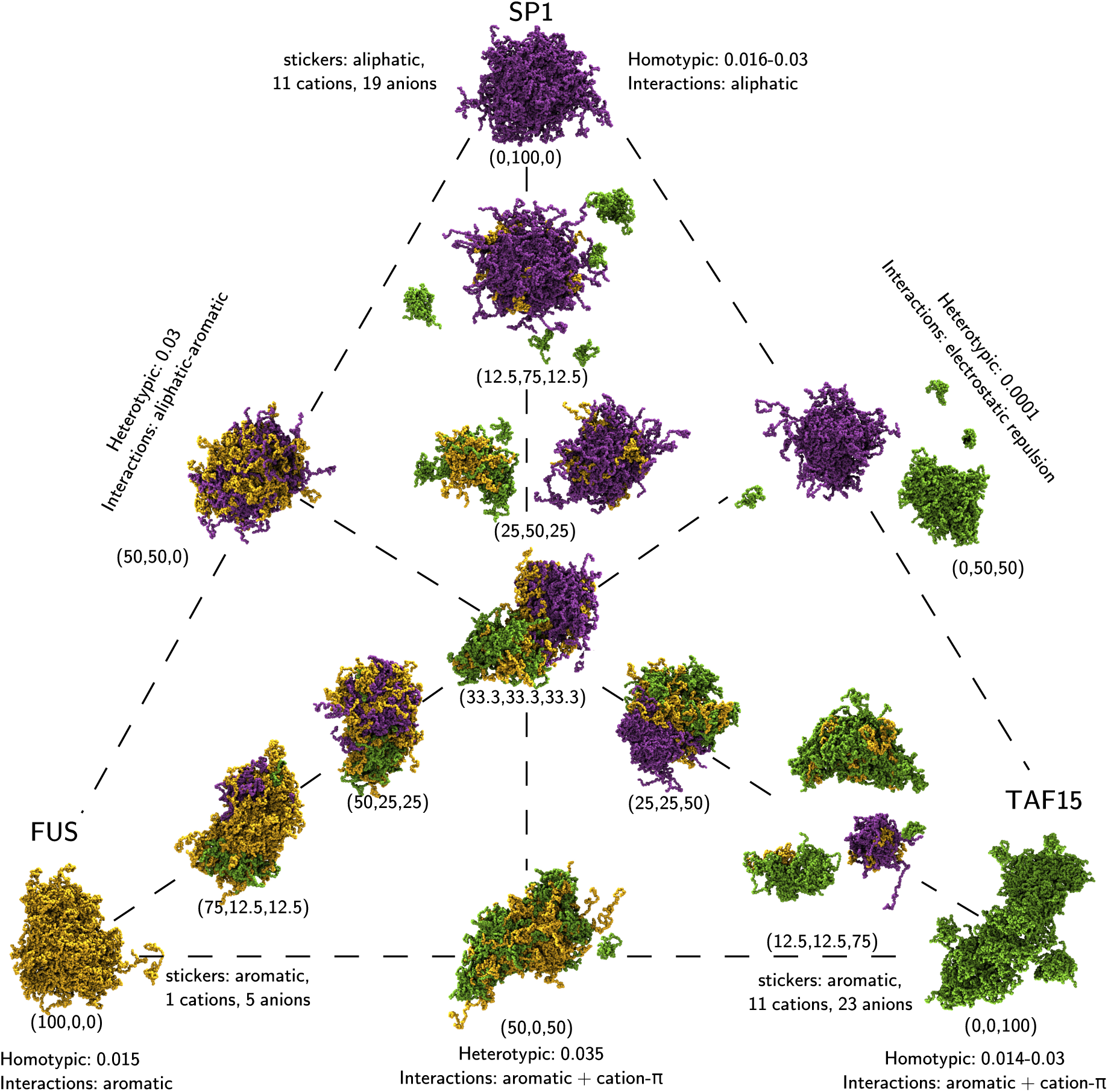
Snapshots of a selection of the ternary droplets. The percentage compositions are described in the image. The end frame of 3*µ*s of simulation is displayed as a representative state. Composition as a percentage is shown next to the droplets in brackets: (% FUS, % SP1, % TAF15). Droplets are formed in all simulations of these molecules under these concentration conditions, irrespective of composition.

The intermolecular contact maps for the three-component mixtures do not yield any extra information compared to the contact maps from the two-component mixtures. The intermolecular contact maps for all residue pairings exhibit the same interaction patterns, with only the absolute magnitude of each contact map changing (see Figs. S23-S30 in the SI).

In the ternary mixtures (Fig. 5) the competition for interactions is dominated by the competition for the strongest heterotypic interactions. The strongest heterotypic interaction is a function of composition, with FUS-TAF15 the strongest for the majority of compositions, and FUS-SP1 interactions the strongest at 75% SP1 content, and SP1-TAF15 the weakest for all compositions (Fig. S5). The dominance of the FUS-TAF15 interactions drive the formation of FUS-TAF15 marbled condensates, since it out-competes the other heterotypic interactions. Since the FUS-SP1 interactions are also strong, but weaker than SP1-SP1 interactions this promotes the FUS to coat the SP1 droplet. This results in a complex bimodal structure where a FUS-TAF15 marbled droplet is partially coating a SP1 droplet. The intriguing behaviour shown in the simulations is that if sufficient FUS is present, both SP1 and TAF15 are incorporated into the same bimodal droplet even though there interactions are extremely unfavourable (*<*0.001). At lower FUS compositions the ability of FUS to provide sufficient interactions to shield TAF15 from SP1 in the same droplet is reduced and eventually leads to the fragmentation of the droplets as in the binary SP1-TAF15 systems. A characteristic feature of the ternary interactions in Fig. S5 is that the homotypic interactions of SP1 and TAF15 can increase with droplet composition. This is particularly strong for SP1 in high TAF15 fraction mixtures and TAF15 in high SP1 fraction mixtures (Fig. S5(D)-S5(F)), indicating that the presence of the other proteins has a crowding effect through repulsive protein-protein interactions.

We identify four different residue types that can behave as stickers: aromatic, aliphatic, anionic, cationic. The table in Fig. 6A shows the mean interaction propensity calculated for all 34 FUS-SP1-TAF15 systems studied (shown by the orange dots in Fig. 2, data in Figs. S3 and S4 in the SI). This shows that cation-*π* interactions are the strongest interaction type, with 20% of all theoretically possible interactions being formed. The second most important interaction type are the aliphatic interactions which have a greater uncertainty than the cation-*π* interactions. Aromatic interactions are the third most important interaction, with a similar uncertainty to cation-*π*. Electrostatic and aliphatic-aromatic interactions provide the smallest relative contribution to the total. From these mean interaction propensities in the table in Fig. 6A, we compute the possible fraction of interactions we expect to see in any FUS-SP1-TAF15 stochiometry as shown in Fig. 6B-F. In Fig. 6B, aromatic-aliphatic interactions are seen across all compositions, with fewer present in higher FUS/TAF15 compositions. Aromatic interactions occur most frequently in FUS and TAF15 rich mixtures. Cation-*π* and electrostatic interactions occur most frequently at large TAF15 fractions, due to the lower fraction of cationic residues in SP1 and FUS sequences. Aliphatic interactions decrease with decreasing SP1 fractions, since SP1 contains the majority of the aliphatic residues in the sequences investigated. This allows the interactions to be grouped into three main classes: SP1 preferred interactions (aliphatic), FUS/TAF15 preferred interactions (aromatic) and TAF15 preferred interaction (cation-*π* and electrostatic).

**Fig. 6.**
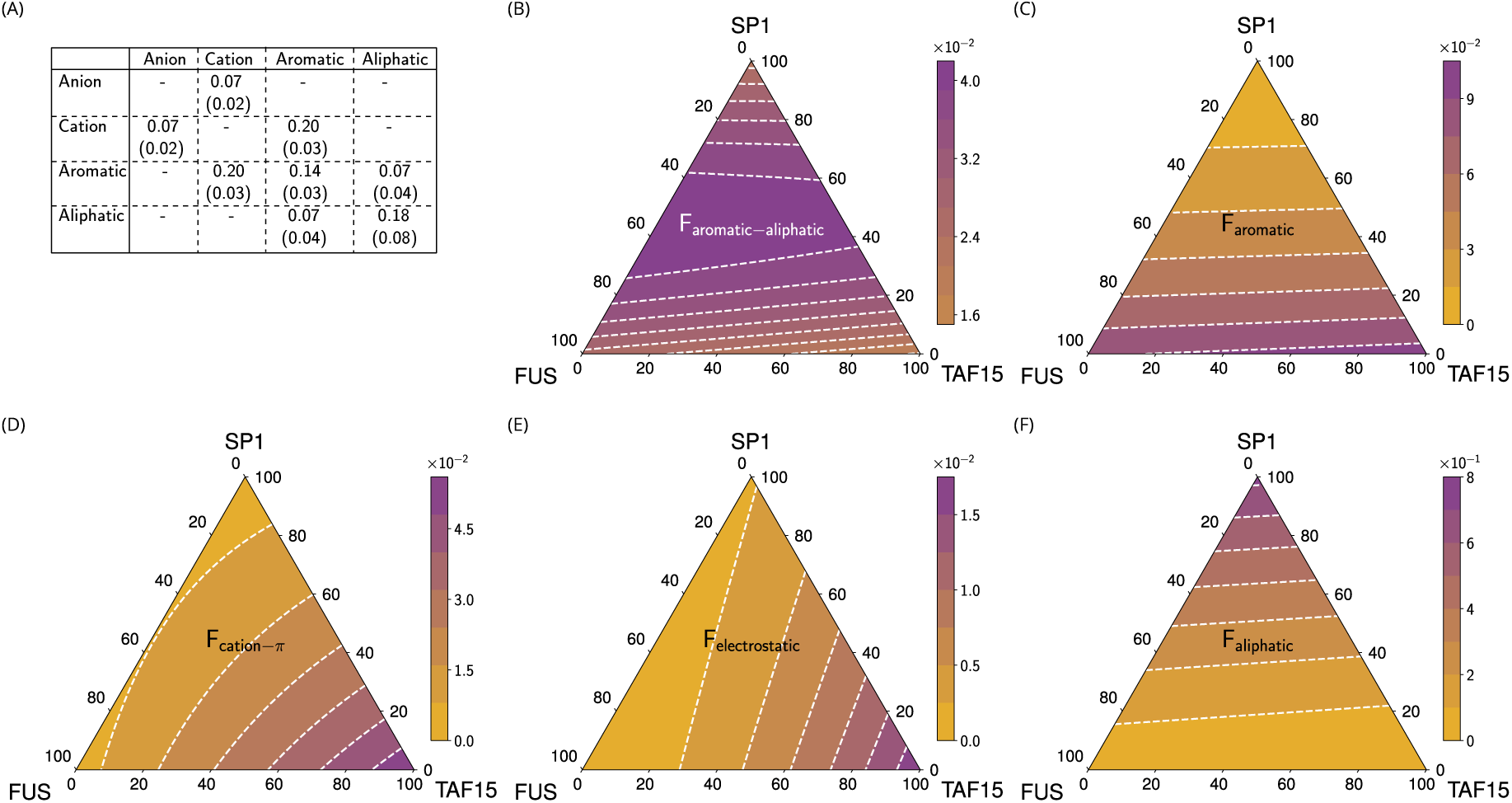
Computed contour plots of the five different interactions for ternary FUS-SP1-TAF15 mixtures. (A) Mean propensity for different attractive interaction types. Standard deviation is displayed in parentheses. The standard deviations indicate an uncertainty of around 20-30% in the relative values of these interactions. The contour maps (B)-(F) were generated using the mean values in table (A). Fraction of (B) Aromatic-aliphatic interactions (C) Aromatic interactions (D) Cation-*π* interactions (E) electrostatic interactions (F) aliphatic interactions as a function of FUS-SP1-TAF15 stoichiometry. Additional details of the calculation are the SI section 3.2.

### 2.5 Universality of TF orthogonality

In this section we extend the analysis to also include the TFs EWS, SP2 and HNF1A. These TFs also undergo PS on their own (Figs. S10-S15) [22, 45]. The IDR of EWS contains a high aromatic content similar to FUS and TAF15, but with more aliphatic residues (Fig. 1 and Fig. S16A). This leads to EWS interactions to be driven by aliphatic contacts from the higher alanine and proline content, and aromatic (tyrosine-tyrosine) contacts (Fig. S16G). A similarly high aliphatic residue content in SP2 as SP1 leads to the same dominance of aliphatic interactions in driving SP2 condensate formation (Fig. S16H). These aliphatic interactions in SP2 are similarly distributed along the entire length of the SP2 IDR in the same manner as SP1 (Fig. 1 and Fig. S16B). HNF1A contains a high aliphatic residue content (Fig. 1 and Fig. S16C), which leads to the dominance of leucine, valine, alanine and proline interactions (Fig. S16F). It is interesting to see that the sticker-spacer paradigm is valid for FUS, TAF15 and EWS, in contrast to SP1, SP2 and HNF1A where it is the more uniform attraction between the aliphatic residues that drive PS.

Next we extend the analysis to binary and ternary systems, following the combinations shown in Table 1. Analysis of the different interaction types of the FUS-TAF15-SP1 system (Fig. 6), the contact maps (Figs. 4, S18-S20) and the ternary phase diagrams (Figs. 2, 5 and S10-15) of all systems studied, it can be concluded that the selectivity of phase separation is related to two major groupings of interactions (shown in Fig. 7A). The first grouping of hydrophobicity-controlled interactions encompasses aliphatic, aromatic and aliphatic-aromatic interactions that utilise stickers that can interact with other residues of the same type. The second grouping consists of cation-controlled interactions which encompasses cation-*π* and electrostatic interactions. Here sticker residues belong to different residue types, so the stickers in the interactions must be heterogeneous, i.e. cationic, anionic and aromatic residues. Interestingly, aromatic and anionic residues both have attractive interactions with cationic residues, but anionic residues have repulsive interactions with aromatic or other anionic residues. This categorisation of interactions highlights the molecular mechanism to achieve orthogonality of phase separation. By exploiting the sticker orthogonality the relative homotypic and heterotypic interaction strengths can be tuned through sequence composition. These orthogonal interactions can be summarised in four characteristic binary droplet morphologies: marbled droplets (such as the SP1-SP2 system) with heterogeneous interactions larger than homotypic interactions (Fig. 7B), coated (core-shell) droplets (such as the EWS-TAF15 system) with heterogeneous interactions of comparable strength to the homotypic interactions (Fig. 7C), bimodal droplets (such as the HNF1A-TAF15 system) with heterogeneous interactions smaller than homotypic interactions (Fig. 7D), separated droplets (such as the SP1-TAF15 system) with heterogeneous interactions significantly smaller (almost negligible) than homotypic interactions (Fig. 7E).

**Fig. 7.**
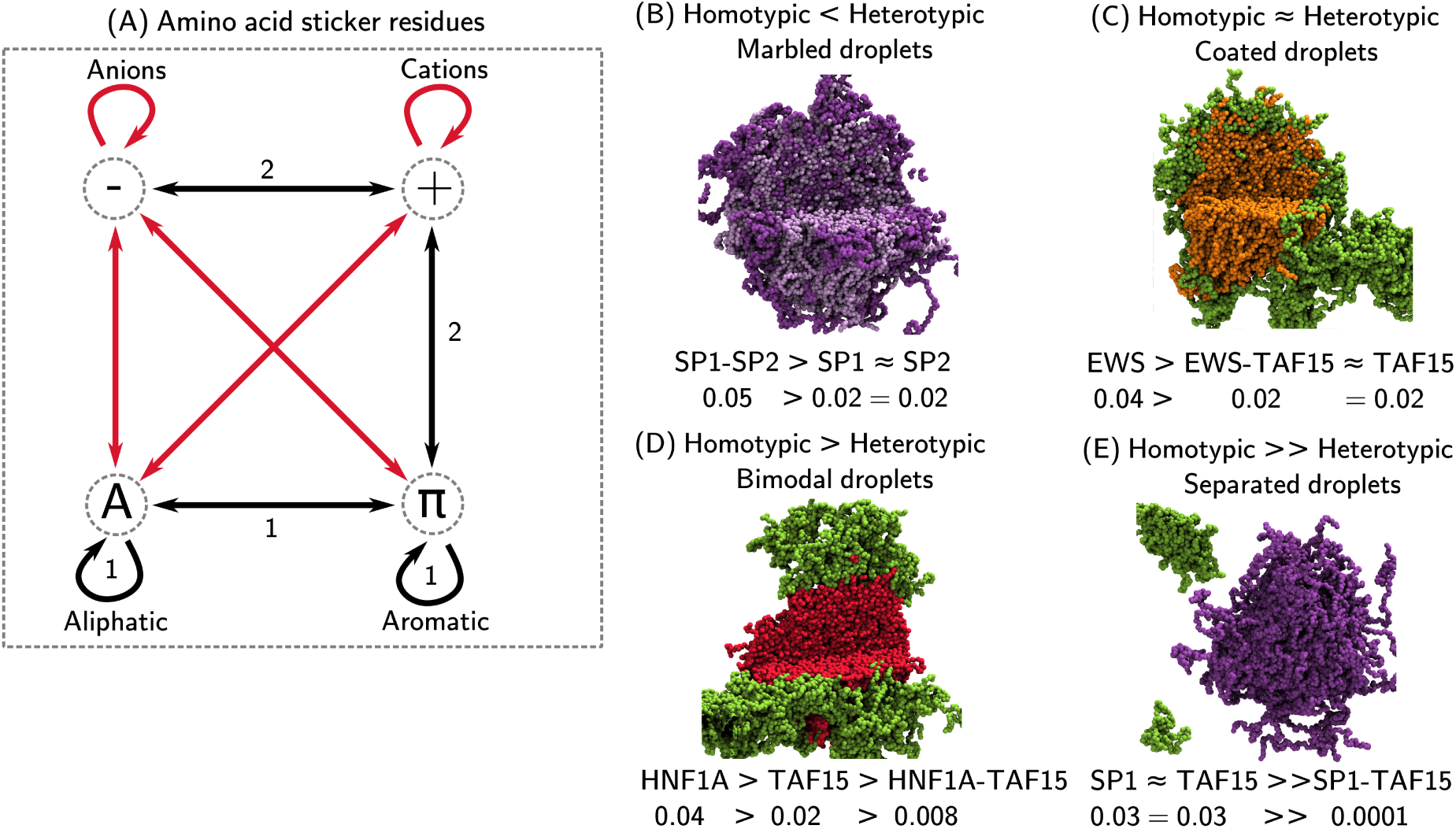
Summary of condensate interactions. (A) Amino acid sticker types and interactions. Favourable interactions (black) promote condensate formation, unfavourable interactions (red) oppose condensate formation. Sticker interactions fall into two orthogonal groupings: 1) Hydrophobic interactions (aliphatic, aromatic and alipatic-aromatic contacts), and 2) Cationic interactions (Cation-*π* and electrostatic). (B)-(E) The four cases of relative homotypic and heterotypic interaction strengths in (50%-50%) mixtures that result in different condensate morphologies, illustrated with examples from simulations undertaken in this work.

The universality of condensate selectivity can be nicely depicted by discussing the new phase diagrams (of Table 1) relative to the reference system of FUS-SP1-TAF15. The first contains the substitution of FUS by EWS. The additional aliphatic residues in EWS lead to some important changes (Fig. S10) when compared to the FUS-SP1-TAF15 triangle (Figs. 2 and 5). Of EWS, SP1, and TAF15 the homotypic interactions of EWS are the strongest (0.025), also stronger than FUS (0.015). Moreover, the additional heterotypic aliphatic interactions of EWS with SP1 (Fig. S19) results in heterotypic EWS-SP1 interactions that are stronger than SP1 homotypic interactions, but weaker than EWS homotypic interactions and causes the formation of a binary SP1-coated EWS droplet (Fig. 7C). The additional aliphatic residue content of EWS compared to FUS leads to more repulsion with the ionic residues in TAF15, resulting in decreased heterotypic interactions of 0.01-0.02 (Fig. S19B), leading to a more patchy binary interaction with TAF15 compared to FUS and a distinct separation from TAF15 in the 3-component system (Fig. 8A and Fig. S10).

**Fig. 8.**
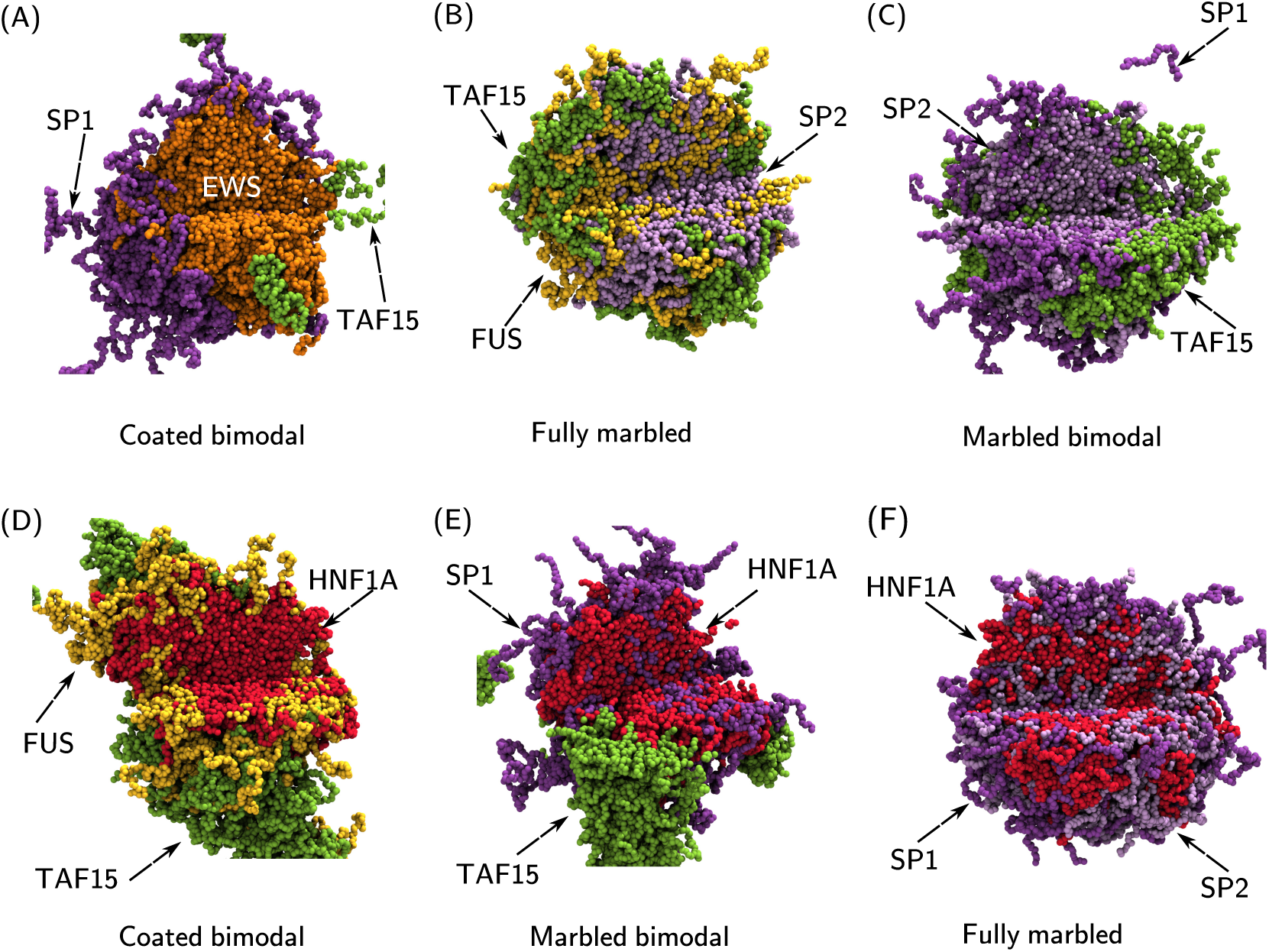
Simulation images for equal-component ternary TF mixtures. (A) EWS-SP1-TAF15 (B) FUS-SP2-TAF15 (C) SP1-SP2-TAF15 (D) FUS-HNF1A-TAF15 (E) HNF1A-SP1-TAF15 (F) HNF1A-SP1-SP2. All simulations were run with a total amino acid concentration of 80000 *µM*. The end frame of 3 *µ*s of simulation is displayed as a representative state. A segment is not displayed to reveal the internal droplet structure. FUS molecules are coloured in yellow, EWS molecules are coloured in orange, SP1 molecules are coloured in purple, SP2 molecules are coloured in light purple, HNF1A molecules are coloured in red, and TAF15 molecules are coloured in green.

The replacement of SP1 by SP2 leads to the FUS-SP2-TAF15 triangle in Fig. 8B and Fig. S11. SP2 has an increased cationic residue content compared to SP1. These cationic residues in SP2 are clustered in three main locations around residue 250, 400 and in the C terminal end (above residue 500), see Fig. 1. These clusters are sufficient to enable the formation of strong electrostatic and cation-*π* interactions with TAF15 (Fig. S18C), and the formation of marbled SP2-TAF15 condensates (Fig. S11). All the heterotypic interactions in this triangle are stronger than homotypic interactions (Fig. S7), resulting in the formation of a three-component marbled droplet (Fig. 8B and Fig. S11). If we then replace FUS by SP2 the phase diagram of SP1-SP2-TAF15 in Fig. S12 is created. The large number of aliphatic residues in both SP1 and SP2 dominate the heterotypic interactions leading to the formation of marbled condensates (see contact maps in Fig. S18B). The heterotypic SP1-SP2 interactions are much stronger than the homotypic interactions (Fig. S7B). Intriguingly, the strong heterotypic interactions of SP2 with both SP1 and TAF15 are able to overcome the repulsive SP1-TAF15 heterotypic interactions (like FUS in the FUS-SP1-TAF15 droplets). In Fig. 8C the equal contribution ternary SP1-SP2-TAF15 mixture results in marbled SP1-SP2 condensates forming around a disk of TAF15, with SP2 acting as a glue at the interface to reduce contact between SP1 and TAF15.

Three distinct cases of ternary phase diagrams are considered with HNF1A. The first case (Fig. S13) is HNF1A with two FET proteins (FUS and TAF15). The heterotypic HNF1A-TAF15 interactions are unfavourable, producing a bimodal condensate with a minimal interfacial region between the HNF1A and TAF15 condensates. This is in contrast to the marbled FUS-HNF1A and FUS-TAF15 condensates where favourable heterotypic interactions are present in the binary mixtures. In the ternary mixture (Fig. 8D), FUS-HNF1A heterotypic contacts dominate over those of FUS-TAF15, leading to formation of a HNF1A condensate coated in FUS with a TAF15 condensate interacting with FUS on a shared interface. The second case (Fig. S14) is HNF1A with one FET (TAF15) and one SP/KLF (SP1) protein. Here SP1-HNF1A heterotypic interactions are the strongest giving rise to the marbled HNF1A-SP1 binary condensates. In the ternary mixture the weak HNF1A-TAF15 heterotypic interactions lead to the association of a TAF15 homotypic domain to the marbled HNF1A-SP1 droplet (Fig. 8E). The third case (Fig. S15) of HNF1A with two SP/KLF family TFs (SP1 and SP2) contains three heterotypic interactions that are more favourable than homotypic interactions (Fig. S7). Therefore a marbled three component droplet is observed for this system driven by the large number of aliphatic residues in all three components.

The HNF1A IDR from the HNF family contains a high number of aliphatic residues, with a small number of aromatic and ionic residues, meaning it more closely resembles the IDRs of the SP/KLF family proteins than the FET proteins. Aliphatic-aromatic interactions dominate the heterotypic interactions of HNF1A with FUS and TAF15 (Figs. S19C and S20C). The heterotypic interactions of HNF1A with SP1 and SP2 are dominated by aliphatic residue contacts (see contact maps in Figs. S20A and B).

### 2.6 Pol II interaction promiscuity overcomes interaction orthogonality

The function of transcription factors is to direct RNAP to the required gene for transcription initiation. The C-terminal domain of RNAP II (referred to as POL II) is known to be able to undergo PS [38–40, 48]. A key question exists: how can POL II circumvent the orthogonality of the interactions displayed by the transcription factors, such that it can reach all required genes to initiate and undertake transcription? The answer lies in the POL II amino acid sequence, which is based on a heptad repeat unit containing the consensus sequence YSPTSPS [37]. POL II homotypic interactions are dominated by aromatic tyrosine contacts (see contact maps in Fig. S17). The use of aromatic interactions in a low charge IDR provides the opportunity to maximise the propensity for co-condensation with any TF, as shown in Fig. 9, where POL II was found to undergo condensation with all TFs in this study (FUS, EWS, HNF1A, SP1, SP2, and TAF15), overcoming the potential barrier posed by the orthogonal interactions used by different TF families. This is achieved by heterotypic hydrophobic interactions of POL II with all TFs (Fig. S21 and S22). The ability for POL II to enter any TF condensate means that the local orthogonal TF condensate environment formed in the nucleus plays a crucial role in orchestrating POL II localisation to the correct genes for transcription.

**Fig. 9.**
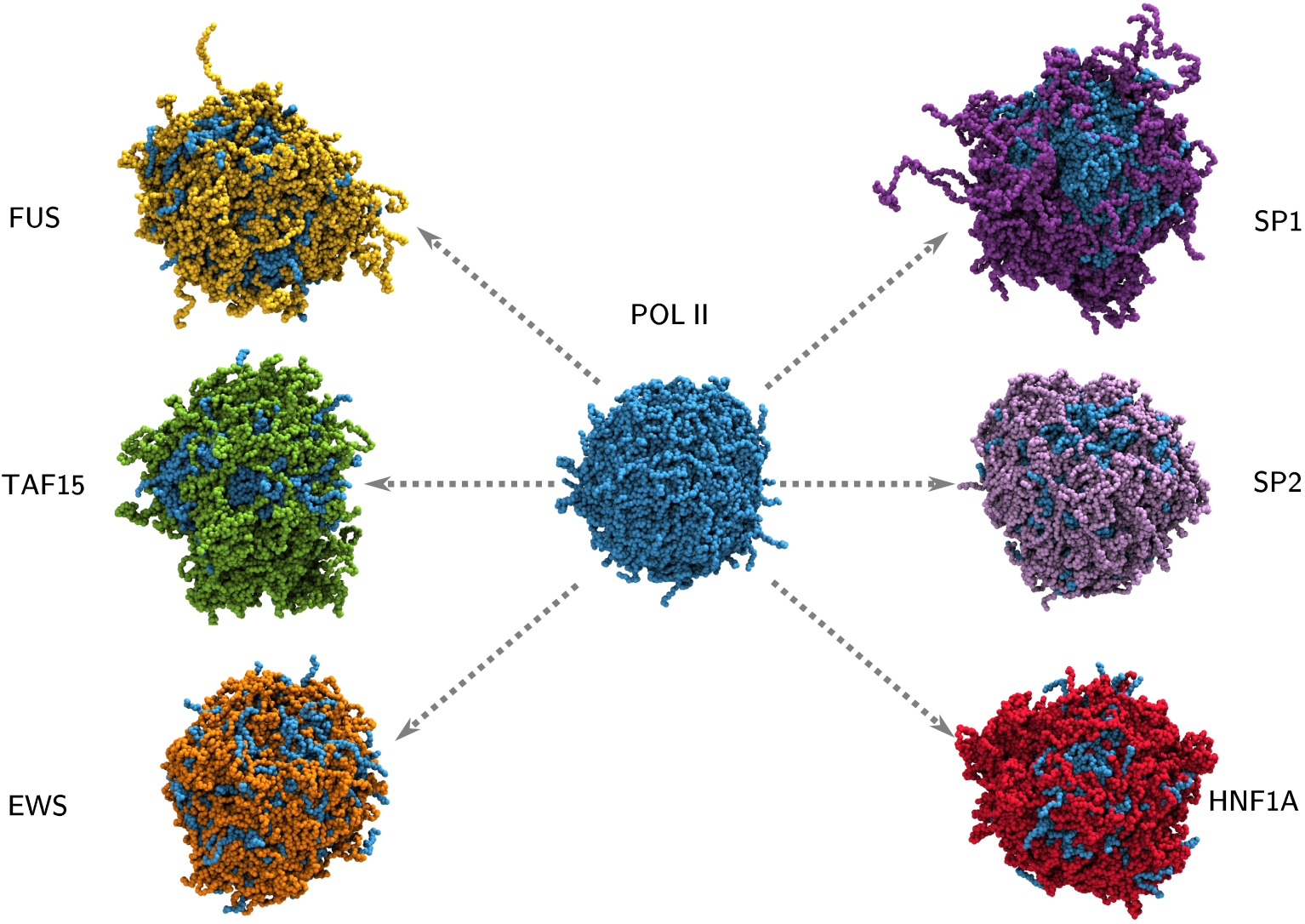
RNA polymerase II (POL II) C terminal domain PS with TFs. Simulations of 60 POL II molecules (centre) and 30 POL II with: 60 FUS (top left), 60 TAF15 (centre left), 60 EWS (bottom left), 30 SP1 (top right), 30 SP2 (centre right), and 45 HNF1A (bottom right). All simulations were run with a total amino acid concentration of 80000 *µM*. The end frame of 3 *µ*s of simulation is displayed as a representative state. POL II molecules are coloured in blue, FUS molecules are coloured in yellow, EWS molecules are coloured in orange, SP1 molecules are coloured in purple, SP2 molecules are coloured in light purple, HNF1A molecules are coloured in red, and TAF15 molecules are coloured in green. Droplets are formed in all simulations of these molecules under these concentration conditions, irrespective of composition.

## 3 Discussion

The nature of the droplets we have observed in our simulations are representative of droplet-spanning percolation networks through phase separation coupled to percolation [36, 46, 49]. From all intermolecular contact maps it can be deduced that at all times percolations exist within both homotypic as well as heterotypic droplets. The intermolecular interactions in the TF condensates, however, are not at all permanent; they are highly dynamic with contact lifetimes on the order of 5 ns (see Fig. S9) such that the cross-links between TFs are transient in nature, similar to FG-nucleoporin condensates [47]. The nature of this phase state can best be categorized as viscoelastic: on one hand the continuous percolations suggest elastic properties, while their transient nature is indicative of a fluid, with mostly spherical morphologies and the ability of two droplets to merge into one [50].

We explored transcriptional condensates consisting of six TFs from three different families. We focused on different spatial levels of molecular interactions by analysing (i) residue scale contact maps (ii) the relative strengths of aliphatic, aromatic, aliphatic-aromatic, electrostatic, and cation-*π* interactions and (iii) the aggregated overall homotypic and heterotypic interaction strength between the molecules. We have found that sticker motifs [36] exist that feature orthogonal driving forces causing PS in ternary TF systems, shown in Fig. 7A. The competitive interactions driving PS within the ternary TF systems can be ranked by their relative strength (the table in Fig. 6A): cation-*π* are the strongest, followed by aliphatic, then aromatic, and finally aliphatic-aromatic and electrostatic are the weakest. The orthogonal sticker motifs use either hydrophobic interactions (aromatic, aliphatic-aromatic, and aliphatic interactions) between Y, A, L, V, P residues, or cation-*π* and electrostatic interactions of R and K with Y, D, or E, to drive PS (Fig. 7A).

PS of the single component droplets is driven by homotypic intermolecular interactions (Fig. 3 and Fig. S16): PS of FUS and EWS is driven by aromatic interactions, TAF15 by electrostatic, cation-*π* and (mostly) aromatic interactions, and SP1, SP2, and HNF1A by aliphatic interactions. The two component mixtures feature the competition of these homotypic intermolecular interactions with heterotypic intermolecular interactions (Fig. 4 and Figs. S18-S20) leading to the identification of four distinct droplet morphologies in binary systems where both species can undergo phase separation: 1) heterotypic interactions being stronger than homotypic interactions leads to a single marbled condensate (Fig. 7B), 2) heterotypic interactions being comparable to homotypic interactions leads to a single coated condensate (Fig. 7C), 3) heterotypic interactions being weaker than homotypic interactions leads to bimodal condensates (Fig. 7D) that have a shared interface. 4) heterotypic interactions being substanially weaker than homotypic interactions leads to two separate condensates (Fig. 7E). These morphologies are also represented in the ternary systems studied, featuring orthogonal phase separation that is at the root of selective gene localisation (as summarised in Fig. 8).

Since one of the orthogonal driving forces for phase separation involves ionic residues, the localized partitioning of transcriptional condensates at the DNA is sentsitive to local electrostatic environment. To explore this we have studied the effect of ion concentration in one specific ternary system that has also been investigated experimentally: FUS-TAF15-SP1 [24]. The results of our exploration of ion concentration on condensation showed that with increasing ion concentration the strength of FUS-TAF15 interactions grows with a corresponding weakening of FUS-SP1 interactions in ternary mixtures, making SP1 inclusion in the condensates less favourable (see SI section 4 and Fig. S6). In living cells, this orthogonality is expected to be even larger, due to crowding or modified local electrostatic environments (for example the presence of large amounts of negatively charged DNA), enhancing the heterotypic FUS-TAF15 interactions at the expense of heterotypic interactions with SP1. This was shown in the previous *in vivo* work of Chong and coworkers [22, 24], where FUS-TAF15 condensates were observed, with SP1 forming separate condensates.

The presence of aromatic residues in both classes of orthogonal sticker motifs provides the mechanism enabling POL II IDR to interact with any TF IDR to enable co-condensation. The ability for POL II IDR to enter all TF IDR condensates provides a simple mechanism that aids in enzyme partitioning to active transcription sites where increased TF concentrations are present. The condensates would enable the RNAP to dock to the correct chromatin location through interaction of its IDR with TFs while at the same time facilitating the active enzymatic site to attach to the DNA and initiate transcription. Orthogonal molecular grammar thus suggests that molecular interactions are evolutionary converged to a state in which subtle differences in amino acid sequence might steer different genetic programs.

Beyond gene expression, orthogonal phase separation might also provide a universal mechanistic basis for the cell to organise targeted (enzymatic) reactions in its highly crowded environment through transient compartmentalization. From an application point of view, such a sequence-based orthogonality can also be exploited for the design of new bio-inspired orthogonal nanoscale soft-matter systems for catalysis [51] and storage [52] while retaining compatibly with biological systems.

## 4 Methods

### 4.1 Protein sequences

The exact sequences for the LCDs used for the work, as shown in 1, are contained in Table S1 of the supporting information.

### 4.2 The 1 bead per amino acid (1BPA) molecular dynamics model

In this work we use the 1BPA model that was previously developed for the study of intrinsically disordered proteins in the nuclear pore complex (NPC) [53, 54]. It has been used extensively to model the behaviour of the disordered nucleoporin regions which fill the center of the yeast NPC and provide a selective barrier to the transport of cargo between the cytoplasm and nucleoplasm [47, 55, 56]. In this work an updated 1BPA model is developed (1BPA-2.1) that extends the applicability domain beyond the intrinsically disordered domains of yeast nucleoporins, by incorporating a greater training set for parameterisation (collating experimental data from [35, 57–60]), and further improves the performance of the previously used 1BPA-2.0 model [45, 61]. The full details of the force field are given in the ESI.

### 4.3 Droplet simulation protocol

To be able to explore the behaviour of the transcription condensates we used droplet formation simulations to be able to look at the internal structure. Self-assembly and clustering of individual monomers into phase separated condensates can be a slow process to observe. To speed up this process we start by forming a condensed phase droplet at the start of the simulation, which is then inserted into an empty dilute phase. If PS is favoured, this droplet structure should remain stable throughout the subsequent simulation; if unstable this droplet would breakup into a dilute phase of monomers.

The initial cubic simulation box is populated with molecules (using a random initial conformation) with their centre of mass placed upon a regular grid, with a small buffer region to avoid overlap between molecules. All simulations are carried out at a temperature of 300 K and use a timestep of 20 fs. For equilibration of the droplet, energy minimisation on the initial configuration is used (energy tolerance of 1 kJ mol^−1^ nm^−1^), then 50 ns NVT Langevin dynamics simulations (Nosé-hoover thermostat with *τ_t_* = 100 ps), followed by 500 ns NPT Langevin dynamics (Nosé-hoover thermostat with *τ_t_* = 100 ps and a Berendsen barostat with *τ_p_* = 10 ps, 1 bar reference pressure and a compressiblity of 4.5 × 10^−5^ bar^−1^). The end state of the NPT equilibration step is inserted into a new p eriodic box with a volume chosen to give a total residue density of 80,000 *µ*M, after recentering on the center of mass and after the molecules have been unwrapped across the previous periodic boundary conditions. A second energy minimisation step is applied in the new simulation box to relax the molecules after the box expansion (energy tolerance of 1 kJ mol^−1^ nm^−1^). A final 3 *µ*s NVT production run (Nosé-hoover thermostat with *τ_t_* = 100 ps) is used for data collection. The trajectory is sampled every 5 ns to determine whether convergence was reached (see SI for details on molecular connectivity graph creation for convergence testing).

## Supplementary information

Supplementary information contains the protein sequences, 1BPA-v2.1 model parameters and comparison to 1BPA-v1.0 and 1BPA-v2.0, data extraction protocol for simulation analysis, simulation images from large scale simulations, molecular interaction data, contact lifetimes, ternary phase diagrams for the additional ternary systems, intermolecular contact maps for systems discussed in the main text.

## Acknowledgements

We thank H.G.S. Arends and J. Postema for discussions. We thank the oLife COFUND project for funding MDD. The COFUND project oLife has received funding from the European Union’s Horizon 2020 research and innovation programme under grant agreement No 847675. We thank the Center for Information Technology of the University of Groningen for their support and for providing access to the Peregrine and Habrok high performance computing clusters. This work made use of the Dutch national e-infrastructure with the support of the SURF Cooperative using grant no. EINF-3233 and EINF-5917.

## Declarations

### 4.5 Author contribution

MDD and PRO designed the research. MDD carried out all simulations, analysed the data, and wrote the first draft. PRO reviewed and edited the article and supervised the research.

