## Supplemental information for "Selective phase separation of transcription factors is driven by orthogonal molecular grammar"

M.D. Driver, P. R. Onck

April 12, 2024

**4 Summary figures** **12**

**5 Simulation interaction data** **14**

**6 Simulation Images** **21**

**7 Contact Lifetimes** **21**

**8 Additional ternary systems** **22**

**9 Contact data** **28**

**10 Simulation data on all TFs** **35**

### 1 Sequence information

| Molecule name | Sequence |
|---|---|
| **FUS** | ASNDYTQQATQSYGAYPTQPGQGYSQQSSQPYGQQSYSGYSQSTDTSGYGQSSYS |
| | SYGQSQNTGYGTQSTPQGYGSTGGYGSSQSSQSSYGQQSSYPGYGQQPAPSSTSG |
| | SYGSSSQSSSYGQPQSGSYSQQPSYGGQQQSYGQQQSYNPPQGYGQQNQYNSSSG |
| | GGGGGGGGGNYGQDQSSMSSGGGSGGGYGNQDQSGGGGSGGYGQQDRG |
| **TAF15** | SDSGSYGQSGGEQQSYSTYGNPGSQGYGQASQSYSGYGQTTDSSYGQNYSGYSSY |
| | GQSQSGYSQSYGGYENQKQSSYSQQPYNNQGQQQNMESSGSQGGRAPSYDQPDYG |
| | QQDSYDQQSGYDQHQGSYDEQSNYDQQHDSYSQNQQSYHSQRENYSHHTQDDRRD |
| | VSRYGEDNRGYGGSQGGGRGRGGYDKDGRGPMTGSSGGD |
| **SP1** | SDQDHSMDEMTAVVKIEKGVGGNNGGNGNGGGAFSQARSSSTGSSSSTGGGGQES |
| | QPSPLALLAATCSRIESPNENSNNSQGPSQSGGTGELDLTATQLSQGANGWQIIS |
| | SSSGATPTSKEQSGSSTNGSNGSESSKNRTVSGGQYVVAAAPNLQNQQVLTGLPG |
| | VMPNIQYQVIPQFQTVDGQQLQFAATGAQVQQDGSGQIQIIPGANQQIITNRGSG |
| | GNIIAAMPNLLQQAVPLQGLANNVLSGQTQYVTNVPVALNGNITLLPVNSVSAAT |
| | LTPSSQAVTISSSGSQESGSQPVTSGTTISSASLVSSQASSSSFFTNANSYSTTT |
| | TTSNMGIMNFTTSGSSGTNSQGQTPQRVSGLQGSDALNIQQNQTSGGSLQAGQQK |
| | EGEQNQQTQQQQILIQPQLVQGGQALQALQAAPLSGQTFTTQAISQETLQNLQLQ |
| | AVPNSGPIIIRTPTVGPNGQVSWQTLQLQNLQVQNPQAQTITLAPMQGVSLGQTS |
| | SSNTTLTPIAS |
| **EWS** | PTDVSYTQAQTTATYGQTAYATSYGQPPTGYTTPTAPQAYSQPVQGYGTGAYDTT |
| | TATVTTTQASYAAQSAYGTQPAYPAYGQQPAATAPTRPQDGNKPTETSQPQSSTG |
| | GYNQPSLGYGQSNYSYPQVPGSYPMQPVTAPPSYPPTSYSSTQPTSYDQSSYSQQ |
| | NTYGQPSSYGQQSSYGQQSSYGQQPPTSYPPQTGSYSQAPSQYSQQSSSYGQQSS |
| **SP2** | MSDPQTSMAATAAVSPSDYLQPAASTTQDSQPSPLALLAATCSKIGPPAVEAAVT |
| | PPAPPQPTPRKLVPIKPAPLPLSPGKNSFGILSSKGNILQIQGSQLSASYPGGQL |
| | VFAIQNPTMINKGTRSNANIQYQAVPQIQASNSQTIQVQPNLTNQIQIIPGTNQA |
| | IITPSPSSHKPVPIKPAPIQKSSTTTTPVQSGANVVKLTGGGGNVTLTLPVNNLV |
| | NASDTGAPTQLLTESPPTPLSKTNKKARKKSLPASQPPVAVAEQVETVLIETTAD |
| | NIIQAGNNLLIVQSPGGGQPAVVQQVQVVPPKAEQQQVVQIPQQALRVVQAASAT |
| | LPTVPQKPSQNFQIQAAEPTPTQVYIRTPSGEVQTVLVQDSPPATAAATSNTTCS |
| | SPASRAPHLSGTSKKHSAAILRKERPLPKIAPAGSIISLNAAQLAAAAQAMQTIN |
| | INGVQVQGVPVTITNTGGQQQLTVQNVSGNNLTISGLSPTQIQLQMEQALAGETQ |
| | PGEKRRRMACTCPNCKDGEKRSGEQGKKK |
| **HNF1A** | KLAMDTYSGPPPGPGPGPALPAHSSPGLPPPALSPSKVHGVRYGQPATSETAEVP |
| | SSSGGPLVTVSTPLHQVSPTGLEPSHSLLSTEAKLVSAAGGPLPPVSTLTALHSL |
| | EQTSPGLNQQPQNLIMASLPGVMTIGPGEPASLGPTFTNTGASTLVIGLASTQAQ |
| | SVPVINSMGSSLTTLQPVQFSQPLHPSYQQPLMPPVQSHVTQSPFMATMAQLQSP |
| | HALYSHKPEVAQYTHTGLLPQTMLITDTTNLSALASLTPTKQVFTSDTEASSESG |
| | LHTPASQATTLHVPSQDPAGIQHLQPAHRLSASPTVSSSSLVLYQSSDSSNGQSH |
| | LLPSNHSVIETFISTQMASSSQ |
| **POL II** | FSPSAASDASGFSPGYSPAWSPTPGSPGSPGPSSPYIPSPGGAMSPSYSPTSPAY |
| | EPRSPGGYTPQSPSYSPTSPSYSPTSPSYSPTSPNYSPTSPSYSPTSPSYSPTSP |
| | SYSPTSPSYSPTSPSYSPTSPSYSPTSPSYSPTSPSYSPTSPSYSPTSPSYSPTS |
| | PSYSPTSPSYSPTSPSYSPTSPSYSPTSPSYSPTSPNYSPTSPNYTPTSPSYSPT |
| | SPSYSPTSPNYTPTSPNYSPTSPSYSPTSPSYSPTSPSYSPSSPRYTPQSPTYTP |
| | SSPSYSPSSPSYSPASPKYTPTSPSYSPSSPEYTPTSPKYSPTSPKYSPTSPKYS |
| | PTSPTYSPTTPKYSPTSPTYSPTSPVYTPTSPKYSPTSPTYSPTSPKYSPTSPTY |
| | SPTSPKGSTYSPTSPGYSPTSPTYSLTSPAISPDDSDEEN |

Table S1: Amino acid residue sequences used in all simulations.

#### 2 1BPA model

The 1BPA model was previously developed for the study of the nuclear pore complex (NPC) [1, 2]. It has been extensively used to model the behaviour of the intrinsically disordered nucleoporins (called FG-Nups) which fill the center of the NPC and provide a selective barrier to the transport of cargo between the cytoplasm and nucleoplasm [3–12].

##### 2.1 Original 1BPA potential

As a starting point for the parameterisation in this work the 1BPA-cp version is used [9, 11] (referred to as 1BPA-1.0 in this work). The bonded potential, $\phi_{\mathrm{b}}$ (equation (1)), consists of three components: bonding ($\phi_{\mathrm{bond}}$, equation (2)), bending ($\phi_{\mathrm{bend}}$) and torsion ($\phi_{\mathrm{torsion}}$) components. The bonding potential, $\phi_{\mathrm{bond}}$ is a simple harmonic potential where $k = 8038$ kJ mol$^{-1}$nm$^{-2}$ and $b = 0.38$ nm. An iterative Boltzmann inversion was used to fit Ramachandran data to generate the bending and torsion potentials, as described in [1]. During this analysis, the presence of glycine or proline residues was found to create distinctive Ramanchandran plots, which resulted in the definition of 6 different angular potentials and nine torsional potentials, described in table S2.

$$\phi_{\mathrm{b}} = \phi_{\mathrm{bond}} + \phi_{\mathrm{bend}} + \phi_{\mathrm{torsion}}, \tag{1}$$

$$\phi_{\mathrm{bond}} = k(r - b)^2, \tag{2}$$

| Bending type | Torsion type |
|---|---|
| ZGX | ZGGX |
| ZPX | ZGPX |
| ZXX | ZGXX |
| ZGP | ZPGX |
| ZPP | ZPPX |
| ZXP | ZPXX |
| | ZXGX |
| | ZXPX |
| | ZXXX |

Table S2: Unique combinations for bending and torsion potentials, using single letter amino acid codes. G is glycine, P is proline, X represents any of the other 18 amino acids, and Z represents any of the 20 amino acid residues. Order is defined from the N-terminus (left) to the C-terminus (right). [1].

There are three components that constitute the non-bonded potential, $\phi_{\mathrm{nb}}$ (equation (3)), of the 1BPA model: hydrophobic interactions ($\phi_{\mathrm{hp}}$, equation (4)), cation-$\pi$ interactions ($\phi_{\mathrm{cp}}$, equation (5)) and electrostatic interactions ($\phi_{\mathrm{el}}$, equation (6)).

$$\phi_{\mathrm{nb}} = \phi_{\mathrm{hp}} + \phi_{\mathrm{cp}} + \phi_{\mathrm{el}}, \tag{3}$$

$$\phi_{\mathrm{hp}} = \begin{cases} \epsilon_{\mathrm{rep}} \left(\frac{\sigma}{r}\right)^8 - \epsilon_{ij} \left[\frac{4}{3}\left(\frac{\sigma}{r}\right)^6 - \frac{1}{3}\right], & r \leq \sigma, \\ (\epsilon_{\mathrm{rep}} - \epsilon_{ij}) \left(\frac{\sigma}{r}\right)^8, & \sigma \leq r, \end{cases} \tag{4}$$

where $\epsilon_{ij} = \epsilon_{\mathrm{hp}} \sqrt{(\epsilon_i \epsilon_j)^{\alpha}}$, $\sigma = 0.6$ nm, $\epsilon_{\mathrm{rep}} = 10$ kJ mol$^{-1}$, $\alpha = 0.27$, $\epsilon_{\mathrm{hp}} = 13.0$ kJ mol$^{-1}$ and $\epsilon_i$, $\epsilon_j$ are residue specific hydrophobicities. The cation-$\pi$ interactions are represented by:

$$\phi_{\mathrm{cp}}(r) = \epsilon_{cp,ij}[3\left(\frac{\sigma_{cp}}{r}\right)^8 - 4\left(\frac{\sigma_{cp}}{r}\right)^6], \tag{5}$$

where $\epsilon_{cp,ij}$ is the energy of the cation-$\pi$ interaction, and $\sigma_{cp} = 0.45$ nm is the radius used for cation-$\pi$ interactions. The parameterisation of $\epsilon_{cp,ij}$ was undertaken by Jafarinia *et al.* [11]. The electrostatic interactions are represented by:

$$\phi_{el} = \frac{q_i q_j}{4\pi\epsilon_0 \epsilon_r(r) r} e^{(-\kappa r)}, \tag{6}$$

$$\epsilon_r(r) = S_s \left[1 - \frac{r^2}{z^2} \frac{e^{r/z}}{\left(e^{r/z} - 1\right)^2}\right] \tag{7}$$

where $S_s = 80$ and $z = 0.25$ nm. The Debye screening coefficient, $\kappa = \left(\epsilon_0 \epsilon_r k_b T)/2N_A e^2 I\right)^{-0.5}$, where $\epsilon_0$ is the permittivity of free space, $\epsilon_r$ is the permittivity of water, $k_b$ is Boltzmann's constant, $N_A$ isAvogadro's number, $e$ is the unit charge of an electron, $T = 300$ K and $I$ equal to the experimental ion concentration (150 mM) unless specified otherwise.

#### 2.2 Parameterisation data

Data originally used in the 1BPA parameterisation for FG-Nups is from Yamada *et al.* [13], which contains the hydrodynamic radii ($R_h$) data for the FG-Nups of yeast. Additional data from the work of Dignon and coworkers for the HPS model [14] and the Mpipi model from Joseph *et al.* [15] contains experimental $R_g$ information on a broader set of IDPs. Additional data on hnRNPA variants from Bremer *et al.* [16] and human Nups from Kapinos *et al.* [17] were incorporated in the reparameterisation. This information is included in Table S8. The total number of IDPs with single molecule data now totals 70 molecules, with a wider variance in charged residue content.

#### 2.3 Updated model parameters in the 1BPA-2.0 and 1BPA-2.1 models

The updates to the basic 1BPA-1.0 model [9, 11] are included in the following tables detailing the changes made: Table S3 for changes to global parameters, Table S4 for hydrophobicity changes, Table S5 for aromatic residue hydrophobicity changes, Table S6 for cation-pi interaction changes, and Table S7 for the neutral hydrophobic potential pairs. These parameters were found to reduce the mean error in $\delta(R_{g/h})$, while maintaining performance on the Yamada FG-Nups data [13] used in the original parameterisation.

| Parameter | 1BPA-1.0 | 1BPA-2.0/2.1 |
|---|---|---|
| $\epsilon_{rep}$ | 10 | 5 |
| $\epsilon_{hp}$ | 13 | 6.5 |
| $\alpha$ | 0.27 | 0.15 |
| $\sigma_{catpi}$ | 0.45 | 0.6 |

Table S3: Parameter changes in the current 1BPA-2.0 and 1BPA-2.1 models, relative to the 1BPA-1.0 model [9, 11], with $\epsilon$ in kJ mol$^{-1}$ and $r$ in nm. The cation-pi scale is systematically offset by 2 kJ mol$^{-1}$, to increase the cation-$\pi$ interaction strengths, shown in Table S6.

| Residue Name | 1BPA-1.0 $\epsilon_i$ | 1BPA-2.0 $\epsilon_i$ | 1BPA-2.1 $\epsilon_i$ |
|---|---|---|---|
| A | 0.70 | 0.70 | 0.70 |
| C | 0.68 | 0.68 | 0.68 |
| D | 0.01 | 0.01 | 0.01 |
| E | 0.01 | 0.01 | 0.01 |
| F | 1.00 | 1.00 | 0.80 |
| G | 0.41 | 0.45 | 0.45 |
| H | 0.53 | 0.56 | 0.56 |
| I | 0.98 | 0.98 | 0.98 |
| K | 0.01 | 0.01 | 0.01 |
| L | 1.00 | 1.00 | 1.00 |
| M | 0.78 | 0.78 | 0.78 |
| N | 0.33 | 0.28 | 0.28 |
| P | 0.65 | 0.67 | 0.67 |
| Q | 0.64 | 0.55 | 0.40 |
| R | 0.01 | 0.01 | 0.01 |
| S | 0.45 | 0.44 | 0.42 |
| T | 0.51 | 0.43 | 0.43 |
| V | 0.94 | 0.94 | 0.94 |
| W | 0.96 | 0.95 | 0.80 |
| Y | 0.82 | 0.95 | 0.55 |

Table S4: Change in hydrophobicity parameters for different amino acid residues in the current 1BPA-2.1 model, relative to the 1BPA-2.0 model and 1BPA-1.0 model [11].

| Aromatic residue | 1BPA-2.1 aromatic $\epsilon_i$ |
|---|---|
| F | 2.60 |
| Y | 3.99 |
| W | 5.98 |
| H | 1.0 |

Table S5: Change in hydrophobicity parameters for aromatic residues in 1BPA-2.1 for interactions with other aromatic residues. This replaces the $\epsilon_i$ in Table S4 for aromatic interactions.

| Cationic residue | Aromatic residue | 1BPA-1.0 $\epsilon_{cp,ij}$ / kJ mol$^{-1}$ | 1BPA-2.X $\epsilon_{cp,ij}$ / kJ mol$^{-1}$ |
|---|---|---|---|
| R | F | 4.3 | 6.3 |
| R | Y | 5.0 | 7.0 |
| R | W | 6.7 | 8.7 |
| K | F | 1.79 | 3.79 |
| K | Y | 3.13 | 5.13 |
| K | W | 4.36 | 6.36 |

Table S6: Change in $\epsilon_{cp,ij}$ interaction energies between different cation-aromatic residue pairs in the current 1BPA 2.X (X is 0 or 1) model, relative to the 1BPA-1.0 model [9, 11].

| Residue 1 | Residue 2 | Reason |
|---|---|---|
| R/K/D/E | R/K/D/E | 1 |
| R/K/D/E | G/S/Q | 2 |

Table S7: Residue pairs set to have steric (volume exclusion) hydrophobic potentials, i.e. $\epsilon_{rep} = \epsilon_{ij}$. Reasons for the changes are as follows: 1) To better balance the hydrophobic and electrostatic interactions, the hydrophobic interactions of the charged amino acids are adjusted to be neutral. 2) To better balance the hydrophobic and electrostatic/cation-$\pi$ interactions, especially in FUS and TAF15, we adjusted the hydrophobic interactions of the charged amino acids with G, S, Q to be steric only.

#### 2.4 Model comparison

Single molecule simulations were run on the molecules previously described to evaluate the performance of the updated parameters. Simulations were run for 10 $\mu$s with the first 0.5 $\mu$s discarded as equilibration, with GROMACS 2019.6 [18]. The change in relative errors are plotted in Figure S1. A decrease in average error $\delta R_{g/h}$ for the complete dataset collated in this work shows the 1BPA-2.1 model is an improvement over the 1BPA-1.0 and 1BPA-2.0 models. The Yamada data [13] shows no change in $\delta R_{g/h}$. Significant reductions in $\delta R_{g/h}$ are seen for the Joseph [15] and Bremer [16] data. Minor improvements in $\delta R_{g/h}$ for the molecules in the Dignon [14] and Kapinos [17] sets are not statistically significant due to the overlapping uncertainty.

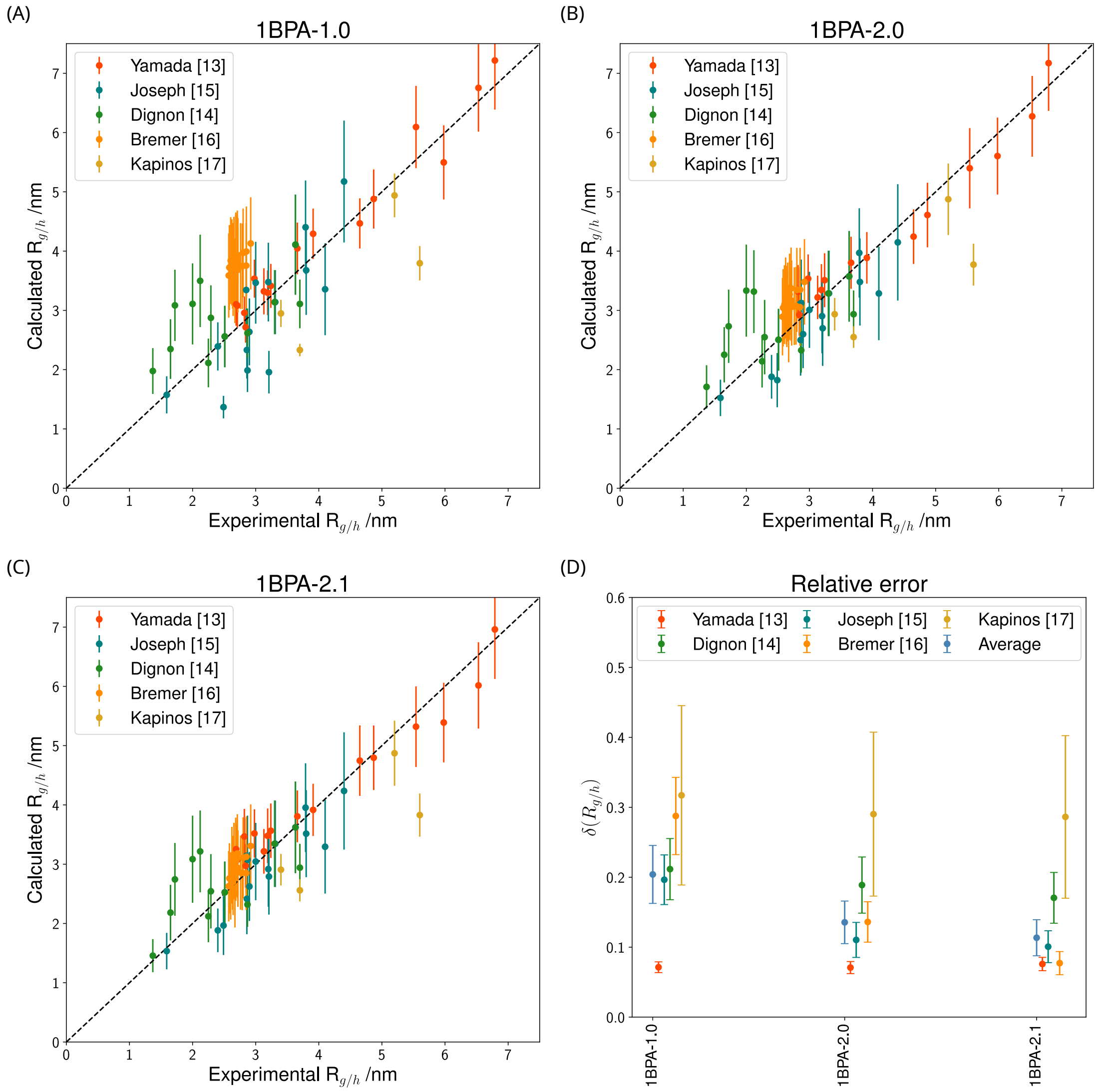

Figure S1: Comparison of computed and experimental radius of gyration, $R_g$ or hydrodynamic (Stokes) radius, $R_h$ for (A) version 1BPA-1.0 (from [11]), (B) version 1BPA-2.0 (used in [19, 20]), (C) version 1BPA-2.1 (developed in this work). (D) $\delta R_{g/h}$, the average relative error between calculated and experimental radius of gyration, $R_g$ or hydrodynamic (Stokes) radius, $R_h$ for the datasets shown in (A)-(C). Each dataset is plotted with a small offset for increased clarity. The average dataset is the average of $\delta R_{g/h}$ for the combination of all molecules.

| Name | $R_{g,exp}$/ nm | $R_{h,exp}$/ nm | Source | V1 $R_h$/ nm | V1 $R_g$/ nm | V2 $R_h$/ nm | V2 $R_g$/ nm | V2.1 $R_h$/ nm | V2.1 $R_g$/ nm |
|---|---|---|---|---|---|---|---|---|---|
| A1-LCD+12D | 2.80 (0.01) | - | [16] | 3.35 (0.30) | 3.97 (0.75) | 2.99 (0.31) | 3.35 (0.71) | 2.86 (0.31) | 3.11 (0.68) |
| A1-LCD+12E | 2.85 (0.01) | - | [16] | 3.36 (0.30) | 3.99 (0.76) | 2.99 (0.31) | 3.35 (0.71) | 2.87 (0.31) | 3.13 (0.68) |
| A1-LCD+2R | 2.62 (0.02) | - | [16] | 3.24 (0.29) | 3.81 (0.73) | 2.89 (0.30) | 3.21 (0.68) | 2.77 (0.30) | 2.98 (0.65) |
| A1-LCD+4D | 2.72 (0.03) | - | [16] | 3.27 (0.30) | 3.83 (0.73) | 2.88 (0.31) | 3.16 (0.68) | 2.75 (0.30) | 2.91 (0.63) |
| A1-LCD+7F-7Y | 2.72 (0.01) | - | [16] | 3.18 (0.30) | 3.68 (0.72) | 2.82 (0.30) | 3.06 (0.65) | 2.76 (0.30) | 2.94 (0.64) |
| A1-LCD+7K+12D | 2.92 (0.01) | - | [16] | 3.43 (0.31) | 4.13 (0.78) | 3.07 (0.31) | 3.48 (0.72) | 2.97 (0.31) | 3.31 (0.70) |
| A1-LCD+7R | 2.71 (0.01) | - | [16] | 3.32 (0.30) | 3.94 (0.75) | 3.00 (0.30) | 3.37 (0.69) | 2.89 (0.30) | 3.18 (0.66) |
| A1-LCD+8D | 2.69 (0.01) | - | [16] | 3.31 (0.30) | 3.90 (0.75) | 2.94 (0.31) | 3.27 (0.69) | 2.82 (0.30) | 3.04 (0.65) |
| A1-LCD+NLS | 2.58 (0.01) | - | [16] | 3.20 (0.30) | 3.72 (0.72) | 2.82 (0.30) | 3.04 (0.65) | 2.66 (0.28) | 2.76 (0.59) |
| A1-LCD-10R | 2.67 (0.01) | - | [16] | 3.04 (0.30) | 3.50 (0.72) | 2.64 (0.30) | 2.78 (0.65) | 2.48 (0.28) | 2.52 (0.58) |
| A1-LCD-10R+10K | 2.85 (0.01) | - | [16] | 3.22 (0.30) | 3.76 (0.73) | 2.82 (0.30) | 3.05 (0.65) | 2.71 (0.30) | 2.85 (0.62) |
| A1-LCD-12F+12Y | 2.60 (0.02) | - | [16] | 3.27 (0.30) | 3.84 (0.73) | 2.84 (0.30) | 3.10 (0.66) | 2.59 (0.28) | 2.63 (0.57) |
| A1-LCD-3R+3K | 2.63 (0.01) | - | [16] | 3.22 (0.30) | 3.75 (0.72) | 2.84 (0.30) | 3.09 (0.66) | 2.71 (0.30) | 2.85 (0.62) |
| A1-LCD-4D | 2.64 (0.01) | - | [16] | 3.14 (0.29) | 3.60 (0.69) | 2.80 (0.29) | 3.02 (0.63) | 2.63 (0.27) | 2.70 (0.56) |
| A1-LCD-6R | 2.57 (0.01) | - | [16] | 3.10 (0.29) | 3.59 (0.71) | 2.71 (0.30) | 2.89 (0.65) | 2.56 (0.28) | 2.63 (0.59) |
| A1-LCD-6R+6K | 2.79 (0.01) | - | [16] | 3.22 (0.30) | 3.75 (0.72) | 2.83 (0.30) | 3.08 (0.66) | 2.71 (0.30) | 2.85 (0.63) |
| A1-LCD-8F+4Y | 2.71 (0.01) | - | [16] | 3.27 (0.30) | 3.86 (0.74) | 2.87 (0.30) | 3.15 (0.66) | 2.84 (0.30) | 3.09 (0.65) |
| A1-LCD-9F+3Y | 2.68 (0.01) | - | [16] | 3.29 (0.29) | 3.88 (0.73) | 2.88 (0.30) | 3.16 (0.66) | 2.87 (0.30) | 3.13 (0.66) |
| A1-LCD-9F+6Y | 2.65 (0.01) | - | [16] | 3.27 (0.30) | 3.85 (0.73) | 2.86 (0.30) | 3.14 (0.67) | 2.82 (0.30) | 3.06 (0.64) |
| A1-LCD-NLS | 2.76 (0.02) | - | [16] | 3.22 (0.30) | 3.75 (0.72) | 2.83 (0.30) | 3.07 (0.65) | 2.72 (0.30) | 2.86 (0.63) |
| ACTR | 2.51 | - | [15] | 2.19 (0.21) | 2.56 (0.52) | 2.16 (0.21) | 2.50 (0.52) | 2.17 (0.21) | 2.53 (0.52) |
| ACTR | 2.51 (0.13) | - | [14] | 2.19 (0.21) | 2.56 (0.52) | 2.16 (0.21) | 2.50 (0.52) | 2.17 (0.21) | 2.53 (0.52) |
| Ash1 | 2.85 | - | [15] | 2.66 (0.23) | 3.34 (0.63) | 2.55 (0.25) | 3.11 (0.62) | 2.56 (0.25) | 3.14 (0.63) |
| CspTm (4-57) | 1.37 (0.07) | - | [14] | 1.78 (0.16) | 1.98 (0.39) | 1.61 (0.16) | 1.71 (0.37) | 1.47 (0.12) | 1.46 (0.28) |
| IBB | 3.20 | - | [15] | 2.82 (0.25) | 3.48 (0.66) | 2.55 (0.27) | 2.90 (0.63) | 2.56 (0.27) | 2.92 (0.63) |
| IN (4-60) | 2.25 (0.11) | - | [14] | 1.85 (0.17) | 2.11 (0.41) | 1.85 (0.18) | 2.14 (0.44) | 1.84 (0.18) | 2.12 (0.44) |
| K18 | 3.80 | - | [15] | 3.16 (0.33) | 3.68 (0.75) | 3.04 (0.32) | 3.48 (0.74) | 3.06 (0.32) | 3.51 (0.73) |
| K25 | 4.40 | - | [15] | 4.30 (0.42) | 5.17 (1.03) | 3.73 (0.46) | 4.15 (0.98) | 3.78 (0.46) | 4.24 (0.99) |
| N49 | 1.59 | - | [15] | 1.38 (0.12) | 1.57 (0.31) | 1.36 (0.12) | 1.52 (0.31) | 1.37 (0.12) | 1.53 (0.31) |
| N98 | 2.86 | - | [15] | 2.44 (0.25) | 2.33 (0.48) | 2.56 (0.31) | 2.50 (0.60) | 2.50 (0.30) | 2.42 (0.60) |
| NLS | 2.40 | - | [15] | 1.90 (0.14) | 2.39 (0.41) | 1.68 (0.15) | 1.88 (0.37) | 1.68 (0.15) | 1.88 (0.37) |
| NSP | 4.10 | - | [15] | 3.17 (0.37) | 3.36 (0.78) | 3.13 (0.38) | 3.29 (0.79) | 3.13 (0.38) | 3.29 (0.79) |
| NUL | 3.00 | - | [15] | 2.92 (0.28) | 3.47 (0.69) | 2.68 (0.28) | 3.01 (0.64) | 2.70 (0.29) | 3.04 (0.65) |
| NUS | 2.49 | - | [15] | 1.61 (0.11) | 1.37 (0.19) | 1.86 (0.23) | 1.82 (0.46) | 1.94 (0.24) | 1.97 (0.50) |
| Nsp1m | - | 6.53 (0.01) | [13] | 6.75 (0.74) | 8.05 (1.76) | 6.27 (0.68) | 7.24 (1.55) | 6.02 (0.73) | 6.84 (1.64) |

Continued on next page

| Name | $R_{g,exp}$/ nm | $R_{h,exp}$/ nm | Source | V1 $R_h$/ nm | V1 $R_g$/ nm | V2 $R_h$/ nm | V2 $R_g$/ nm | V2.1 $R_h$/ nm | V2.1 $R_g$/ nm |
|---|---|---|---|---|---|---|---|---|---|
| Nsp1n | - | 2.71 (0.00) | [13] | 3.08 (0.37) | 3.24 (0.76) | 3.08 (0.38) | 3.23 (0.78) | 3.08 (0.39) | 3.22 (0.79) |
| Nup100n | - | 4.87 (0.04) | [13] | 4.88 (0.50) | 4.50 (0.96) | 4.61 (0.54) | 4.02 (0.94) | 4.80 (0.54) | 4.27 (0.94) |
| Nup100s | - | 3.66 (0.03) | [13] | 4.05 (0.44) | 4.71 (0.99) | 3.80 (0.44) | 4.29 (0.96) | 3.81 (0.44) | 4.28 (0.95) |
| Nup116m 1c | - | 4.65 (0.00) | [13] | 4.47 (0.42) | 3.97 (0.68) | 4.24 (0.46) | 3.62 (0.77) | 4.75 (0.59) | 4.40 (1.09) |
| Nup116s | - | 3.91 (0.02) | [13] | 4.29 (0.43) | 5.08 (1.01) | 3.89 (0.44) | 4.37 (0.94) | 3.92 (0.44) | 4.43 (0.97) |
| Nup145Ns | - | 2.98 (0.00) | [13] | 3.53 (0.32) | 3.70 (0.65) | 3.54 (0.41) | 3.81 (0.86) | 3.52 (0.41) | 3.77 (0.85) |
| Nup145n | - | 2.82 (0.02) | [13] | 2.96 (0.28) | 2.70 (0.55) | 3.31 (0.45) | 3.24 (0.87) | 3.47 (0.46) | 3.50 (0.91) |
| Nup159 | - | 5.54 (0.02) | [13] | 6.09 (0.69) | 7.10 (1.59) | 5.40 (0.68) | 5.87 (1.44) | 5.32 (0.68) | 5.75 (1.43) |
| Nup1c | - | 3.24 (0.04) | [13] | 3.41 (0.37) | 3.16 (0.69) | 3.51 (0.45) | 3.33 (0.86) | 3.56 (0.46) | 3.43 (0.88) |
| Nup1m | - | 6.79 (0.02) | [13] | 7.22 (0.83) | 8.11 (1.79) | 7.17 (0.81) | 8.13 (1.76) | 6.96 (0.83) | 7.78 (1.82) |
| Nup2 | - | 5.98 (0.03) | [13] | 5.50 (0.63) | 6.07 (1.35) | 5.60 (0.65) | 6.38 (1.44) | 5.39 (0.67) | 6.01 (1.45) |
| Nup42 | - | 2.84 (0.05) | [13] | 2.72 (0.26) | 2.37 (0.43) | 2.93 (0.36) | 2.72 (0.64) | 2.97 (0.37) | 2.78 (0.67) |
| Nup49 | - | 2.69 (0.00) | [13] | 3.10 (0.36) | 3.18 (0.80) | 3.14 (0.40) | 3.16 (0.83) | 3.25 (0.42) | 3.36 (0.87) |
| Nup57 | - | 3.19 (0.10) | [13] | 3.30 (0.38) | 3.20 (0.75) | 3.34 (0.44) | 3.24 (0.84) | 3.48 (0.46) | 3.49 (0.91) |
| Nup60 | - | 3.13 (0.02) | [13] | 3.32 (0.39) | 3.81 (0.87) | 3.22 (0.37) | 3.59 (0.80) | 3.22 (0.38) | 3.60 (0.82) |
| P53 | 2.87 | - | [15] | 2.06 (0.18) | 1.99 (0.37) | 2.65 (0.32) | 3.12 (0.73) | 2.63 (0.33) | 3.08 (0.74) |
| ProTalpha | 3.79 | - | [15] | 3.37 (0.27) | 4.40 (0.79) | 3.06 (0.25) | 3.97 (0.75) | 3.06 (0.25) | 3.95 (0.75) |
| ProTalpha-C (56-110) | 3.70 (0.19) | - | [14] | 2.23 (0.12) | 3.11 (0.41) | 2.12 (0.13) | 2.94 (0.40) | 2.13 (0.13) | 2.94 (0.40) |
| ProTalpha-N (2-56) | 2.87 (0.14) | - | [14] | 2.08 (0.14) | 2.62 (0.42) | 1.96 (0.14) | 2.33 (0.38) | 1.95 (0.14) | 2.32 (0.38) |
| Protein-L | 1.65 (0.14) | - | [14] | 2.00 (0.20) | 2.35 (0.50) | 1.96 (0.19) | 2.25 (0.47) | 1.93 (0.20) | 2.18 (0.47) |
| R15 (6-99) | 1.72 (0.09) | - | [14] | 2.61 (0.24) | 3.08 (0.60) | 2.43 (0.27) | 2.73 (0.62) | 2.43 (0.27) | 2.74 (0.61) |
| R17 (6-99) | 2.29 (0.11) | - | [14] | 2.43 (0.21) | 2.87 (0.55) | 2.35 (0.29) | 2.55 (0.63) | 2.35 (0.28) | 2.54 (0.63) |
| SH4-UD | 2.90 | - | [15] | 2.34 (0.25) | 2.64 (0.56) | 2.31 (0.25) | 2.60 (0.57) | 2.32 (0.25) | 2.62 (0.58) |
| Sic1 | 3.21 | - | [15] | 2.09 (0.19) | 1.96 (0.36) | 2.43 (0.28) | 2.70 (0.64) | 2.48 (0.28) | 2.79 (0.64) |
| alpha-synuclein | 3.31 | - | [15] | 2.97 (0.26) | 3.14 (0.54) | 3.03 (0.34) | 3.29 (0.72) | 3.06 (0.34) | 3.34 (0.73) |
| alpha-synuclein | 3.30 (0.30) | - | [14] | 2.97 (0.26) | 3.14 (0.54) | 3.03 (0.34) | 3.29 (0.72) | 3.06 (0.34) | 3.34 (0.73) |
| cNup153 Human | - | 5.20 (3.20) | [17] | 4.94 (0.37) | 4.26 (0.64) | 4.88 (0.60) | 4.36 (1.19) | 4.87 (0.55) | 4.34 (1.04) |
| cNup214 Human | - | 3.40 (1.50) | [17] | 2.95 (0.23) | 2.49 (0.45) | 2.94 (0.27) | 2.48 (0.48) | 2.91 (0.27) | 2.46 (0.47) |
| cNup62 Human | - | 3.70 (1.70) | [17] | 2.33 (0.11) | 1.80 (0.13) | 2.55 (0.18) | 2.06 (0.26) | 2.56 (0.19) | 2.08 (0.27) |
| cNup98 Human | - | 5.60 (1.60) | [17] | 3.80 (0.29) | 3.14 (0.57) | 3.77 (0.35) | 3.10 (0.64) | 3.83 (0.36) | 3.18 (0.68) |
| hCyp (3,167) | 2.00 (0.10) | - | [14] | 3.00 (0.32) | 3.11 (0.68) | 3.11 (0.37) | 3.33 (0.78) | 2.97 (0.36) | 3.08 (0.73) |
| hNHE1cdt | 3.63 | - | [15] | 3.38 (0.34) | 4.11 (0.84) | 3.11 (0.34) | 3.57 (0.76) | 3.13 (0.34) | 3.62 (0.77) |
| hNHE1cdt | 3.63 (0.18) | - | [14] | 3.38 (0.34) | 4.11 (0.84) | 3.11 (0.34) | 3.57 (0.76) | 3.13 (0.34) | 3.62 (0.77) |
| sNase | 2.12 (0.10) | - | [14] | 3.08 (0.30) | 3.50 (0.78) | 2.98 (0.31) | 3.32 (0.70) | 2.92 (0.31) | 3.22 (0.69) |

Continued on next page

| Name | $R_{g,exp}$/ nm | $R_{h,exp}$/ nm | Source | V1 $R_h$/ nm | V1 $R_g$/ nm | V2 $R_h$/ nm | V2 $R_g$/ nm | V2.1 $R_h$/ nm | V2.1 $R_g$/ nm |
|---|---|---|---|---|---|---|---|---|---|

Table S8: Calculated radius of gyration, $R_g$ or hydrodynamic (Stokes) radius, $R_h$ for the original 1BPA-1.0 parameters (from [11]), labelled V1, and the 1BPA 2.0 parameters developed in this work, labelled V2. Experimental data (including information on data source) is also present for comparison with the calculated data. Standard deviations are in brackets. $R_h$ is calculated using the HullRad method [21]. Numbers in brackets next to the names of the molecules represent the amino acid ranges used in both experiments and simulations for $R_g$ measurements.

### 3 Simulation analysis using a molecular connectivity graph

To assess condensate stability in the droplet simulations, a molecular connectivity graph, consisting of nodes and edges, is used to describe the clustering and assess the time evolution of the molecular network during the simulation. Each simulation frame can be described by a connectivity graph. In the graph representation each node corresponds to a unique molecule, with an edge between the nodes denoting the number of non-zero interactions (if zero interactions are present no edge is added). Within an edge information can be stored about the number and type of contacts between molecules, based on residue categorisation. An interaction was determined to be present using a cutoff of 0.7 nm between 1BPA residues. Computation and processing of the contact matrix for a simulation frame allows the creation of the molecule graph. This process is shown in figure S2.

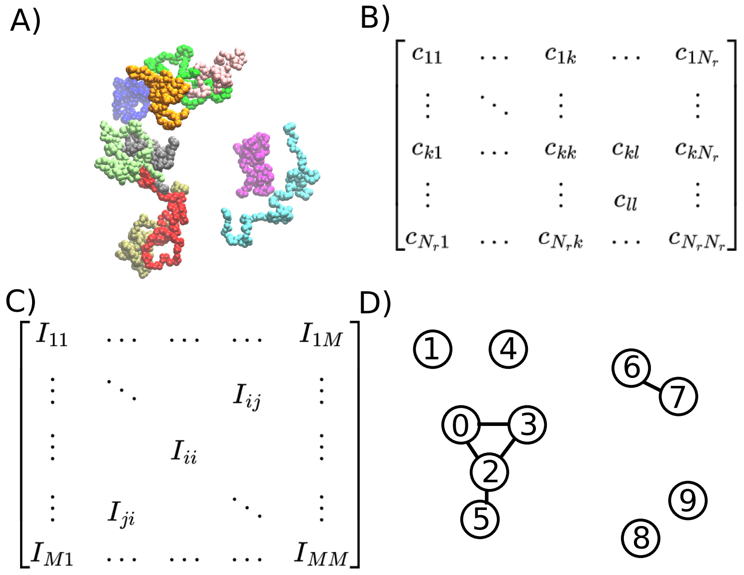

Figure S2: Processing of a contact matrix for one trajectory frame. (A) Visualisation of molecules in the trajectory frame. The particles of the 10 molecules are coloured based on the molecule type. (B) The contact matrix for a frame is computed between all residues, yielding an $N_r \times N_r$ matrix, with a value of 1 if a contact is present between particles $k$ and $l$, and 0 otherwise. (C) The contact matrix in (B) can be decomposed into distinct sub-matrices by grouping together residue interactions belonging to unique molecule pairs. This produces a new matrix representation, where element $I_{ij}$ is the contact matrix between molecules $i$ and $j$. (D) A molecular graph derived from the contact matrix in (C). When constructing the molecular graph we consider only the elements where $i \geq j$ to avoid double counting of interactions. We iterate over the $I_{ij}$ matrices ($i \neq j$), to add edges to the graph, storing information about the number of intermolecular contacts between molecules $i$ and $j$. Similarly, a summation of intramolecular contacts $I_{ii}$ for like molecules (case when $i = j$) is also stored. This process is repeated for all frames in a simulation with a sampling rate of once every 50 ns.

#### 3.1 Readouts from a molecular connectivity graph

A molecular connectivity graph computed for a simulation can be used to get several different readouts about a simulation system. Information on the size of clusters can be found by identifying the discrete sub-graphs of nodes with only internal connections (no contacts outside of the cluster). Through assessment of the time evolution of the number of clusters, grouped based on number of molecules, and the number of molecules in clusters within specific size categories it is possible to determine whether the system has reached equilibrium. After convergence is established it is possible to examine the distribution of components within the different clusters. The percentage of species $i$ in a cluster is computed using $N_{i \in C}/(N_{frames} N_i)$ where $N_{i \in C}$ is the number of molecules of $i$ in a cluster of size $C$.

#### 3.2 Contact map definition

From the system graph creation process, contact matrices for all molecule combinations, over all frames are computed. This information is also useful for understanding the specific residues which drive condensation. As such, this information is aggregated during graph computation. Contact maps can be divided into two categories: intramolecular contact maps (interactions between particles in the same molecule copy), corresponding to the diagonal sub-matrices ($I_{ii}$) in figure S2, and intermolecular contact maps (interactions between particles of two different molecules), corresponding to the off-diagonal sub-matrices ($I_{ij}$) in figure S2. Since individual molecules of the same type are indistinguishable at the macroscale, only unique combinations of molecules need to be stored in different matrices, such that a single contact map for each intermolecular pairing and intramolecular contact map is computed. These maps contain the data for all copies of the same molecule pairs and for all timeframes studied. The contact maps by particle index (Figures 3A-C and 4A-C in manuscript and Figures S23-S30 parts A,D,G- if D or G present) display the contact information as a contact probability: the contact information is normalised by the number of frames, $N_{frames}$ and $((N_iN_j)^{0.5})$ where $N_i$ and $N_j$ are the number of copies of molecule types $i$ and $j$ in the simulation.

When the contact information is presented based on residue type in a molecule (Figures 3D-F and 4D-F in manuscript and Figures S23-S30 parts B,E,H- if E or H present), a summation over the information in the contact maps by particle index to group interactions by residue type (equivalent to a matrix reduction operation). The presentation of normalised residue contact maps (Figures S23-S30 parts C,F,I- if F or I present) applies a second normalisation step to the contact map by residue type of $(N_{i,r1}N_{j,r2})^{0.5}$, with $N_{j,r1}$ the number of residue $r1$ in molecule $i$ and $N_{j,r2}$ the number of residue $r2$ in molecule $j$.

#### Interaction summary information

Data on the distribution of interactions between different interaction types is also extracted from the contact maps for a system. The contact data is aggregated by summation over the contact maps and grouped into five categories of attractive interactions: aromatic, aliphatic, aliphatic-aromatic, cation-$\pi$, electrostatic. To compute the fraction of interactions, $F_{int}$, between two species (Figures 3D-F and 4D-F in manuscript) the number of interactions is divided by the total interactions between the species. The fraction of interactions, $F_{int}$, for a system (Figures S3 and S4) is the sum of the interactions from all contact maps (intramolecular and intermolecular) in the system divided by the total number of all interactions in the system. The theoretical upper bound of the number of interactions that are possible between two classes of residue types is $N_iN_j$, where $N_i$ and $N_j$ are the number of residues of type $i$ and $j$, respectively, in the simulation. By normalising the number of actual interactions by the upper bound $N_iN_j$ it is possible to compute the mean interaction propensity for a ternary mixture. This was done for FUS-SP1-TAF15 mixtures to generate the data in Figure 7. From these mean interaction propensities in the table in Figure 7A, we compute the possible fraction of interactions we expect to see in any FUS-SP1-TAF15 stochiometry by multiplying the interaction propensities in the table of Figure 7A with the number of residue types in the composition, as shown in Figure 7B-F.

The propensity for a specific interaction type is the sum of the interactions from all contact maps (intramolecular and intermolecular) in the system divided by the theoretical upper limit for the number of interactions, $N_iN_j$, where $N_i$, $N_j$ are the number of residues of type $i$ and $j$ that form the interaction. The mean propensity (Figure 7A) is the average propensity from all systems at 150mM ion concentration (34 compositions). To be able to generate the interaction triangles displaying $F_{int}$ for all compositions (Figure 7B-F and Figures SI), the mean propensity in Figure 7A is multiplied by $N_iN_j$ in the system (to get the expected number of interactions for the given composition), and divided by the expected total number of interactions $(0.026N_{tot}N_{tot})$ .

Intermolecular contact propensity between two molecules of different types (in Figures S5 and S6, and Tables S9-S19) is computed by summation of the interactions in the corresponding intermolecular contact map for the species involved. To enable comparison between different compositions normalisation by the theortical upp $N_iL_iN_jL_j$ is used, where $N_i$ and $N_j$ are the number of copies of molecule types $i$ and $j$, and $L_i$ and $L_j$ are the number of simulation residues per molecule of $i$ and $j$, respectively.

### 4 Summary figures

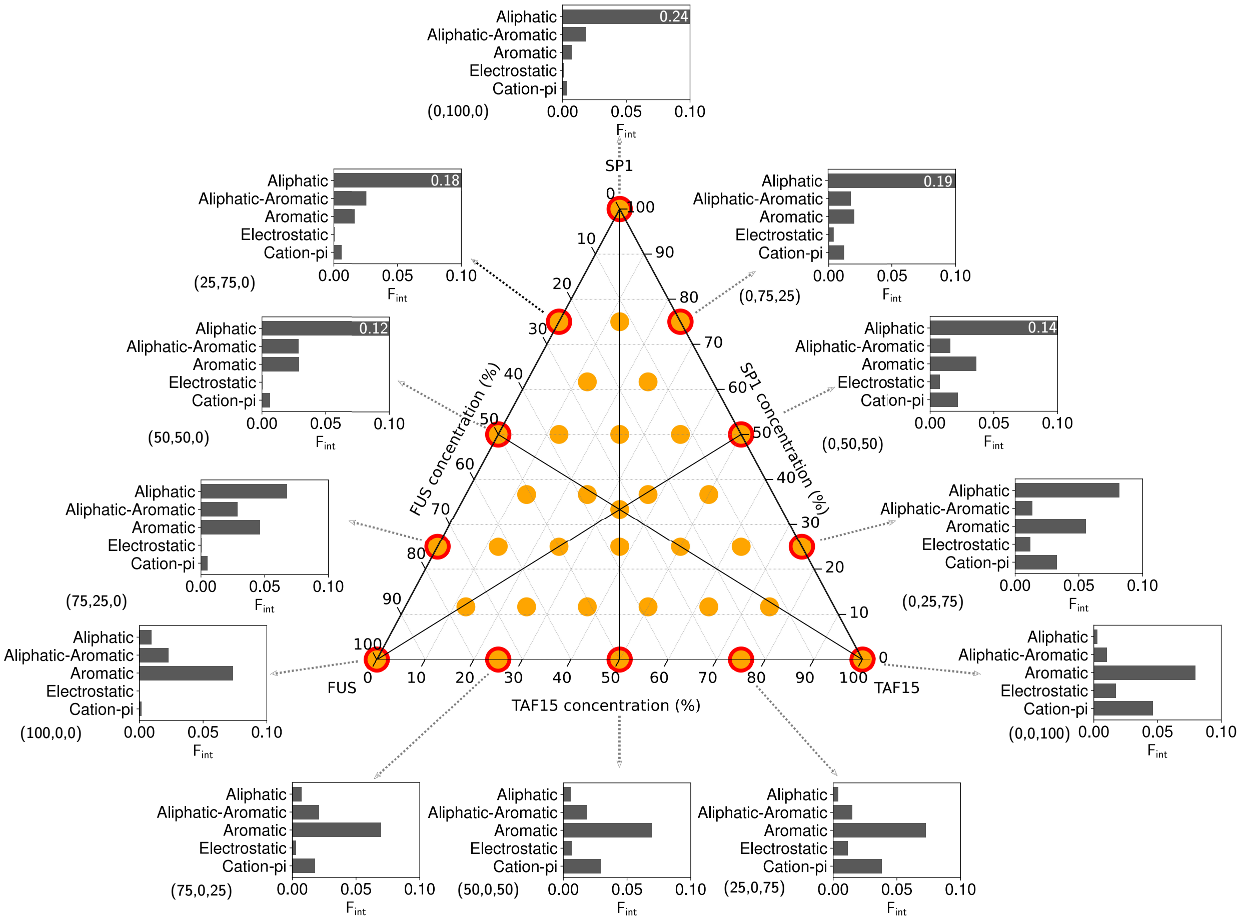

Figure S3: **Ternary phase diagram displaying the interaction type distribution of the two component droplets.** Ternary phase diagram showing the simulations undertaken in this work. The percentage compositions are described in the image. Interaction summaries for droplet simulations of one and two component systems at 150 mM and 300 K. Interactions are aggregated by type and normalised by the total number of interactions. Composition as a percentage is shown next to the droplets in brackets: (% FUS, % SP1, % TAF15).

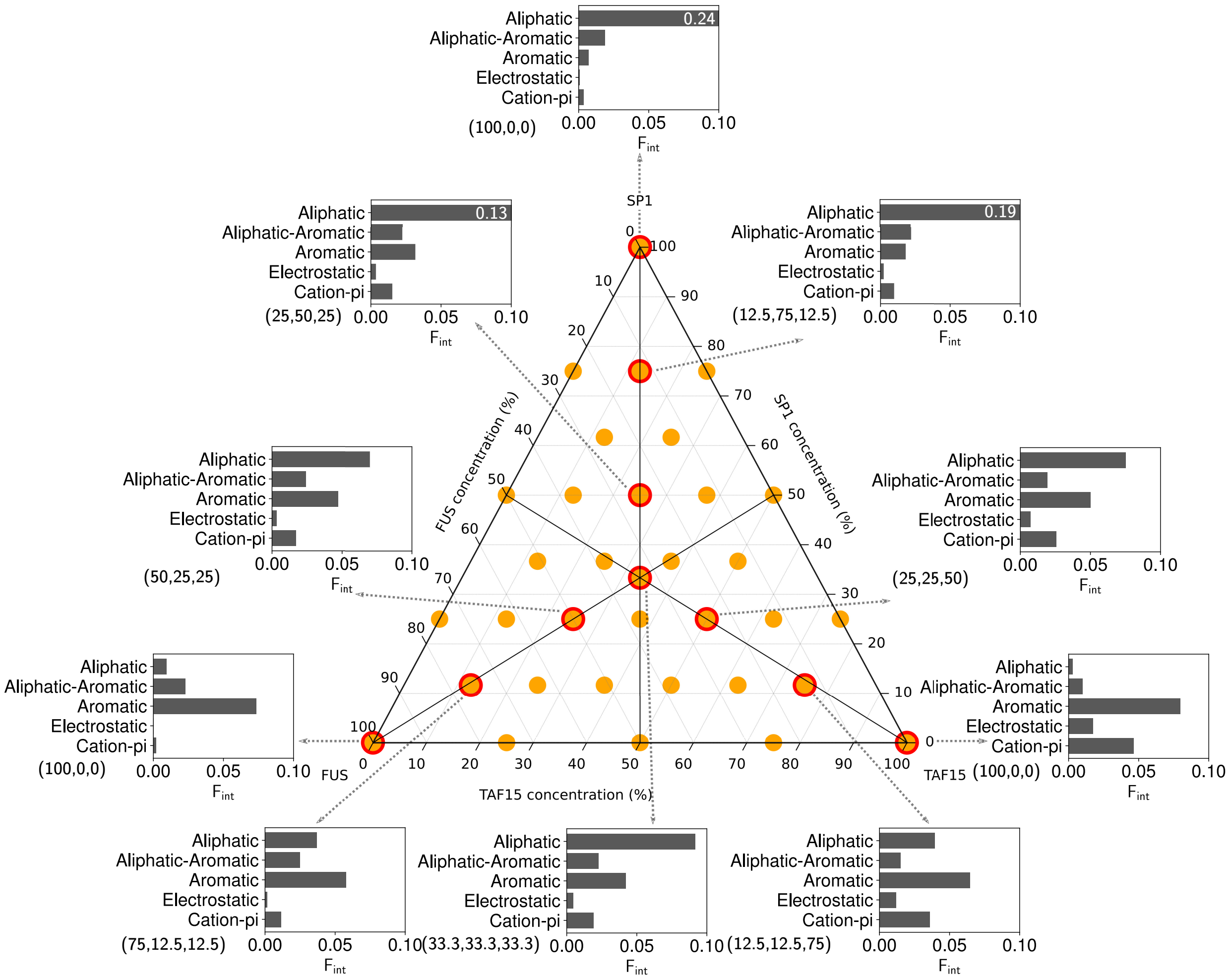

Figure S4: Ternary phase diagram showing the simulations undertaken in this work. Simulations were all run with an amino acid concentration of 80000 $\mu M$. The total composition of a species are defined relative to 120 FUS molecules, 120 TAF15 molecules, and 60 SP1 molecules, to give percentage compositions described in the image. Interaction summaries for droplet simulations of three component systems at 150 mM and 300 K. Interactions are aggregated by type and normalised by the total number of interactions. Composition as a percentage is shown next to the droplets in brackets: (% FUS, % SP1, % TAF15).

### 5 Simulation interaction data

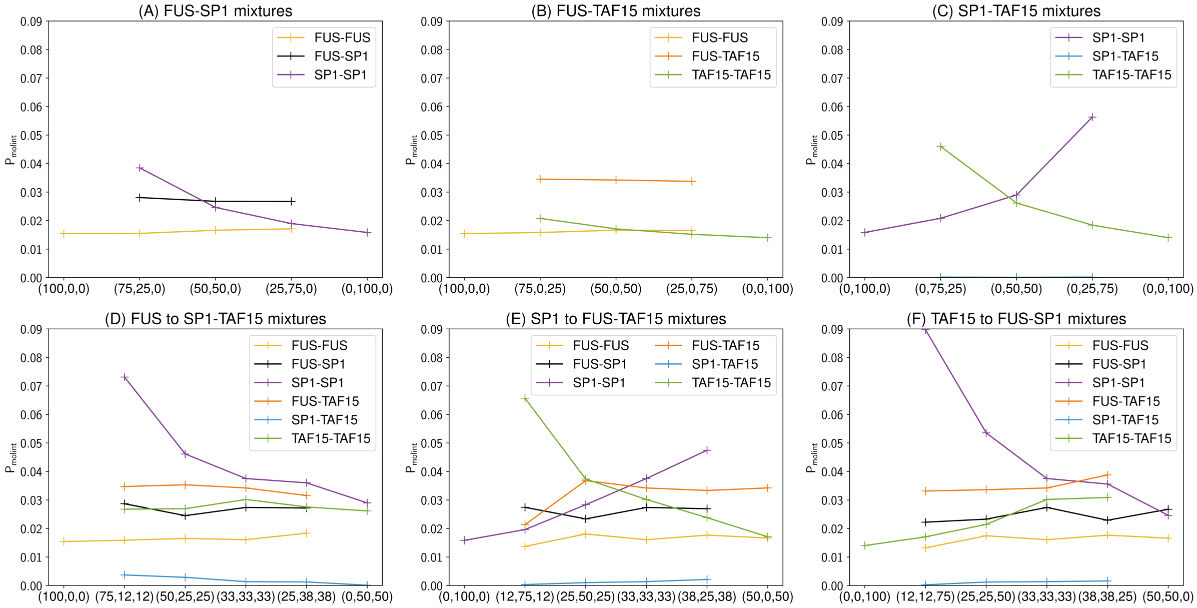

Figure S5: (A)-(F) Interaction propensity of molecule interactions, $P_{\mathsf{molint}}$, for (A) FUS-SP1 mixtures, (B) FUS-TAF15 mixtures, (C) SP1-TAF15 mixtures, (D) FUS to equal SP1-TAF15 mixtures, (E) SP1 to equal FUS-TAF15 mixtures, (F) TAF15 to equal FUS-SP1 mixtures. The total composition of a species are defined relative to 120 FUS molecules, 120 TAF15 molecules, and 60 SP1 molecules, to give percentage compositions. Composition as a percentage is shown on the $x$ axis in brackets: (% FUS, % SP1, % TAF15).

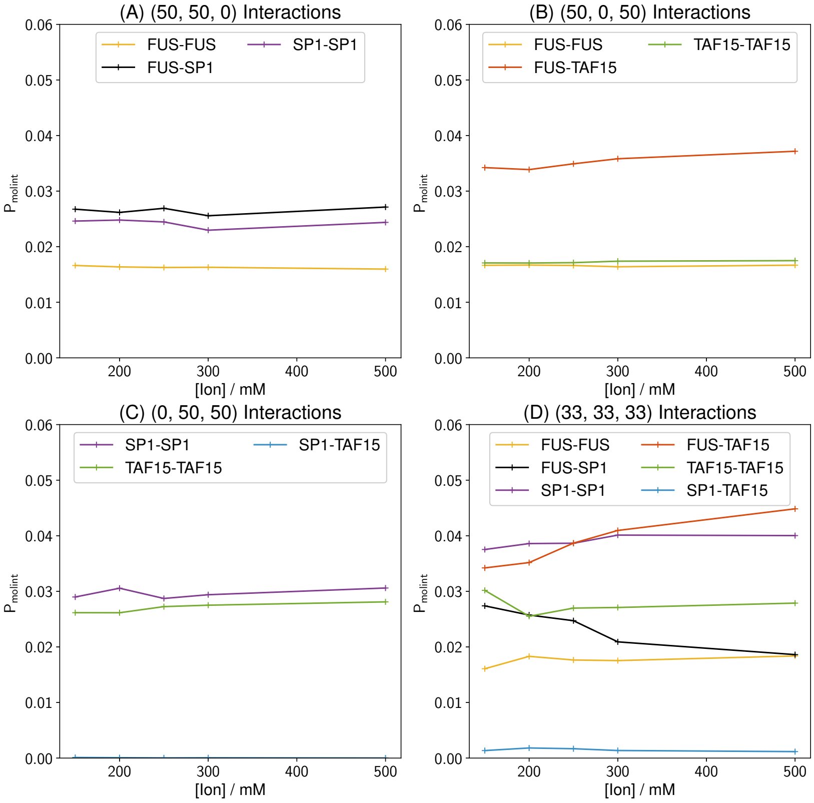

Figure S6: (A)-(D) Interaction propensity of molecule interactions, $P_{molint}$, as a function of ion concentration for (A) (50, 50, 0), (B) (50, 0, 50), (C) (0, 50, 50). (D) (33, 33, 33). The total composition of a species are defined relative to 120 FUS molecules, 120 TAF15 molecules, and 60 SP1 molecules, to give percentage compositions. Composition as a percentage is in brackets: (% FUS, % SP1, % TAF15).

Increased ion concentrations of up to 500 mM, compared to the 150 mM (used to represent biological conditions) were explored for a subset of compositions on the triangle in Fig. 2 (corners, midpoint of the edges, and centre of the triangle). The largest effect was observed in the ternary mixture, where the relative strength of FUS-TAF15 increased to become the strongest interaction at 500 mM, replacing SP1 homotypic interactions as the strongest intermolecular interactions in the system (Fig. S6D). Simultaneously the FUS-SP1 interactions became progressively weaker with increased ion concentration (Tables S9-S13 for data). This is due to the decreased repulsion between the anionic residues in FUS and TAF15, allowing easier formation of cation-$\pi$ interactions between the molecules. The result of the changing interaction strengths manifested as a greater exclusion of the SP1 from the merged droplet, leading to formation of a FUS-TAF15 condensate with an SP1 condensate interacting with FUS on the surface of the FUS-TAF15 droplet. As expected, no significant effect is seen in the FUS, SP1 and 50:50 FUS-SP1 compositions, where aromatic-aliphatic and aliphatic-aliphatic interactions are dominant since these are unperturbed by changes to ionic strength (Fig. S6). In TAF15, 50:50 FUS-TAF15, and 50:50 SP1-TAF15 a minor effect on TAF15 homotypic and FUS-TAF15 heterotypic interactions were observed due to the decreased repulsion between the anionic residues in TAF15 (Fig. S6). Intriguingly the effect of the presence of a third species that enables the utilisation of different interaction motifs leads to much more complexity in the emergent behaviour than the systems with only one or two components.

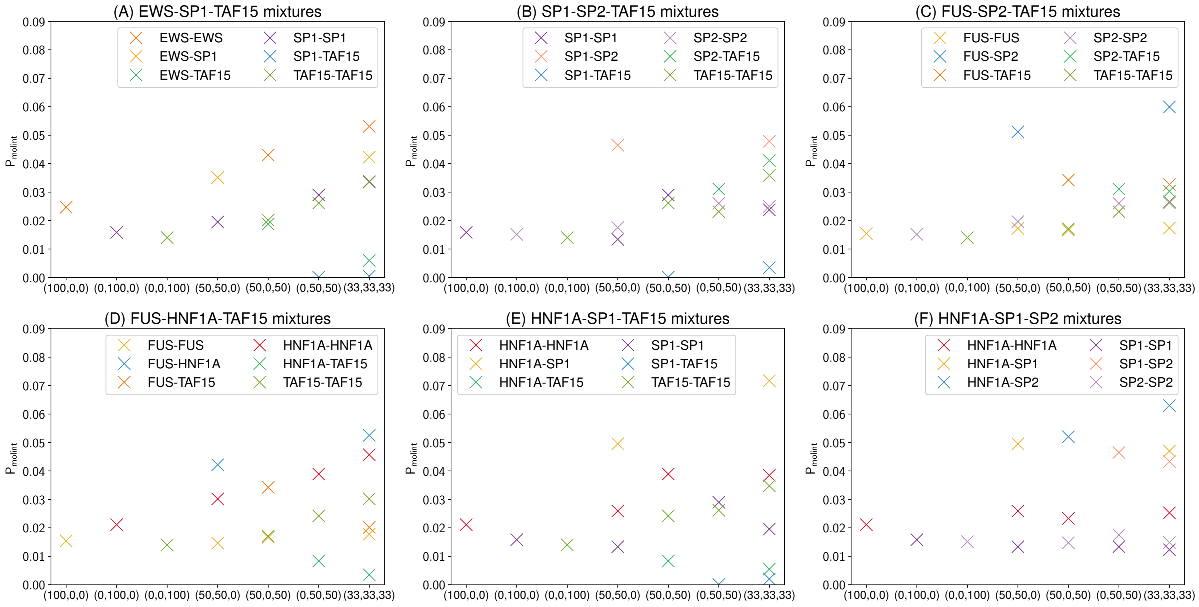

Figure S7: (A)-(F) Interaction propensity of molecule interactions, $P_{molint}$, for (A) EWS-SP1-TAF15 mixtures, (B) SP1-SP2-TAF15 mixtures, (C) FUS-SP2-TAF15 mixtures, (D) FUS-HNF1A-TAF15 mixtures, (E) HNF1A-SP1-TAF15 mixtures, (F) HNF1A-SP1-SP2 mixtures. The total composition of a species are defined relative to 120 FUS molecules, 120 TAF15 molecules, 120 EWS molecules, 60 SP1 molecules, 60 SP2 molecules, and 90 HNF1A molecules to give percentage compositions. Composition as a percentage is shown on the $x$ axis in brackets: (% species 1, % species 2, % species 3).

| Composition (% FUS, % SP1, % TAF15) | FUS-FUS | FUS-SP1 | FUS-TAF15 | SP1-SP1 | SP1-TAF15 | TAF15-TAF15 |
|---|---|---|---|---|---|---|
| (100, 0, 0) | 0.015 | 0.000 | 0.000 | 0.000 | 0.000 | 0.000 |
| (0, 100, 0) | 0.000 | 0.000 | 0.000 | 0.016 | 0.000 | 0.000 |
| (0, 0, 100) | 0.000 | 0.000 | 0.000 | 0.000 | 0.000 | 0.014 |
| (25, 75, 0) | 0.017 | 0.027 | 0.000 | 0.019 | 0.000 | 0.000 |
| (25, 0, 75) | 0.017 | 0.000 | 0.034 | 0.000 | 0.000 | 0.015 |
| (50, 50, 0) | 0.017 | 0.027 | 0.000 | 0.025 | 0.000 | 0.000 |
| (50, 0, 50) | 0.017 | 0.000 | 0.034 | 0.000 | 0.000 | 0.017 |
| (75, 25, 0) | 0.015 | 0.028 | 0.000 | 0.038 | 0.000 | 0.000 |
| (75, 0, 25) | 0.016 | 0.000 | 0.035 | 0.000 | 0.000 | 0.021 |
| (0, 25, 75) | 0.000 | 0.000 | 0.000 | 0.056 | 0.000 | 0.018 |
| (0, 50, 50) | 0.000 | 0.000 | 0.000 | 0.029 | 0.000 | 0.026 |
| (0, 75, 25) | 0.000 | 0.000 | 0.000 | 0.021 | 0.000 | 0.046 |
| (12, 25, 62) | 0.015 | 0.026 | 0.034 | 0.052 | 0.000 | 0.021 |
| (12, 75, 12) | 0.014 | 0.027 | 0.021 | 0.020 | 0.000 | 0.066 |
| (12, 12, 75) | 0.013 | 0.022 | 0.033 | 0.090 | 0.000 | 0.017 |
| (25, 25, 50) | 0.017 | 0.023 | 0.034 | 0.054 | 0.001 | 0.021 |
| (25, 50, 25) | 0.018 | 0.023 | 0.037 | 0.028 | 0.001 | 0.037 |
| (33, 33, 33) | 0.016 | 0.027 | 0.034 | 0.038 | 0.001 | 0.030 |
| (38, 25, 38) | 0.018 | 0.027 | 0.033 | 0.047 | 0.002 | 0.024 |
| (38, 12, 50) | 0.018 | 0.024 | 0.035 | 0.090 | 0.003 | 0.020 |
| (50, 25, 25) | 0.017 | 0.025 | 0.035 | 0.046 | 0.003 | 0.027 |
| (50, 12, 38) | 0.017 | 0.027 | 0.033 | 0.087 | 0.003 | 0.021 |
| (75, 12, 12) | 0.016 | 0.029 | 0.035 | 0.073 | 0.004 | 0.027 |
| (12, 37, 50) | 0.016 | 0.026 | 0.035 | 0.039 | 0.001 | 0.023 |
| (12, 50, 38) | 0.013 | 0.020 | 0.041 | 0.030 | 0.000 | 0.029 |
| (12, 62, 25) | 0.015 | 0.027 | 0.032 | 0.023 | 0.000 | 0.041 |
| (25, 37, 38) | 0.018 | 0.027 | 0.032 | 0.036 | 0.001 | 0.028 |
| (25, 62, 12) | 0.017 | 0.026 | 0.035 | 0.023 | 0.003 | 0.045 |
| (25, 12, 62) | 0.018 | 0.031 | 0.033 | 0.091 | 0.002 | 0.018 |
| (38, 37, 25) | 0.018 | 0.023 | 0.039 | 0.036 | 0.002 | 0.031 |
| (38, 50, 12) | 0.017 | 0.026 | 0.034 | 0.026 | 0.002 | 0.048 |
| (50, 37, 12) | 0.016 | 0.027 | 0.035 | 0.031 | 0.003 | 0.047 |
| (62, 25, 12) | 0.017 | 0.023 | 0.040 | 0.045 | 0.003 | 0.035 |
| (62, 12, 25) | 0.016 | 0.027 | 0.035 | 0.074 | 0.004 | 0.022 |

Table S9: Number of interactions between species for different compositions at a temperature of 300K and 150 mM ion concentration, normalised by $N_1L_1N_2L_2$, where $N_1$, $N_2$ are the number of copies of species 1 and species 2 in the system, and $L_1$, $L_2$ are the lengths of species 1 and species 2, respectively.

| Composition (% FUS, % SP1, % TAF15) | FUS-FUS | FUS-SP1 | FUS-TAF15 | SP1-SP1 | SP1-TAF15 | TAF15-TAF15 |
|---|---|---|---|---|---|---|
| (100, 0, 0) | 0.015 | 0.000 | 0.000 | 0.000 | 0.000 | 0.000 |
| (0, 100, 0) | 0.000 | 0.000 | 0.000 | 0.016 | 0.000 | 0.000 |
| (0, 0, 100) | 0.000 | 0.000 | 0.000 | 0.000 | 0.000 | 0.014 |
| (50, 50, 0) | 0.016 | 0.026 | 0.000 | 0.025 | 0.000 | 0.000 |
| (50, 0, 50) | 0.017 | 0.000 | 0.034 | 0.000 | 0.000 | 0.017 |
| (0, 50, 50) | 0.000 | 0.000 | 0.000 | 0.031 | 0.000 | 0.026 |
| (33, 33, 33) | 0.018 | 0.026 | 0.035 | 0.039 | 0.002 | 0.026 |

Table S10: Number of interactions between species for different compositions at a temperature of 300K and 200 mM ion concentration, normalised by $N_1L_1N_2L_2$, where $N_1$, $N_2$ are the number of copies of species 1 and species 2 in the system, and $L_1$, $L_2$ are the lengths of species 1 and species 2, respectively.

| Composition (% FUS, % SP1, % TAF15) | FUS-FUS | FUS-SP1 | FUS-TAF15 | SP1-SP1 | SP1-TAF15 | TAF15-TAF15 |
|---|---|---|---|---|---|---|
| (100, 0, 0) | 0.016 | 0.000 | 0.000 | 0.000 | 0.000 | 0.000 |
| (0, 100, 0) | 0.000 | 0.000 | 0.000 | 0.016 | 0.000 | 0.000 |
| (0, 0, 100) | 0.000 | 0.000 | 0.000 | 0.000 | 0.000 | 0.015 |
| (50, 50, 0) | 0.016 | 0.027 | 0.000 | 0.024 | 0.000 | 0.000 |
| (50, 0, 50) | 0.017 | 0.000 | 0.035 | 0.000 | 0.000 | 0.017 |
| (0, 50, 50) | 0.000 | 0.000 | 0.000 | 0.029 | 0.000 | 0.027 |
| (33, 33, 33) | 0.018 | 0.025 | 0.039 | 0.039 | 0.002 | 0.027 |

Table S11: Number of interactions between species for different compositions at a temperature of 300K and 250 mM ion concentration, normalised by $N_1L_1N_2L_2$, where $N_1$, $N_2$ are the number of copies of species 1 and species 2 in the system, and $L_1$, $L_2$ are the lengths of species 1 and species 2, respectively.

| Composition (% FUS, % SP1, % TAF15) | FUS-FUS | FUS-SP1 | FUS-TAF15 | SP1-SP1 | SP1-TAF15 | TAF15-TAF15 |
|---|---|---|---|---|---|---|
| (100, 0, 0) | 0.015 | 0.000 | 0.000 | 0.000 | 0.000 | 0.000 |
| (0, 100, 0) | 0.000 | 0.000 | 0.000 | 0.016 | 0.000 | 0.000 |
| (0, 0, 100) | 0.000 | 0.000 | 0.000 | 0.000 | 0.000 | 0.015 |
| (50, 50, 0) | 0.016 | 0.026 | 0.000 | 0.023 | 0.000 | 0.000 |
| (50, 0, 50) | 0.016 | 0.000 | 0.036 | 0.000 | 0.000 | 0.017 |
| (0, 50, 50) | 0.000 | 0.000 | 0.000 | 0.029 | 0.000 | 0.028 |
| (33, 33, 33) | 0.018 | 0.021 | 0.041 | 0.040 | 0.001 | 0.027 |

Table S12: Number of interactions between species for different compositions at a temperature of 300K and 300 mM ion concentration, normalised by $N_1L_1N_2L_2$, where $N_1$, $N_2$ are the number of copies of species 1 and species 2 in the system, and $L_1$, $L_2$ are the lengths of species 1 and species 2, respectively.

| Composition (% FUS, % SP1, % TAF15) | FUS-FUS | FUS-SP1 | FUS-TAF15 | SP1-SP1 | SP1-TAF15 | TAF15-TAF15 |
|---|---|---|---|---|---|---|
| (100, 0, 0) | 0.016 | 0.000 | 0.000 | 0.000 | 0.000 | 0.000 |
| (0, 100, 0) | 0.000 | 0.000 | 0.000 | 0.016 | 0.000 | 0.000 |
| (0, 0, 100) | 0.000 | 0.000 | 0.000 | 0.000 | 0.000 | 0.015 |
| (50, 50, 0) | 0.016 | 0.027 | 0.000 | 0.024 | 0.000 | 0.000 |
| (50, 0, 50) | 0.017 | 0.000 | 0.037 | 0.000 | 0.000 | 0.017 |
| (0, 50, 50) | 0.000 | 0.000 | 0.000 | 0.031 | 0.000 | 0.028 |
| (33, 33, 33) | 0.018 | 0.019 | 0.045 | 0.040 | 0.001 | 0.028 |

Table S13: Number of interactions between species for different compositions at a temperature of 300K and 500 mM ion concentration, normalised by $N_1L_1N_2L_2$, where $N_1$, $N_2$ are the number of copies of species 1 and species 2 in the system, and $L_1$, $L_2$ are the lengths of species 1 and species 2, respectively.

| Composition (% EWS, % SP1, % TAF15) | EWS-EWS | EWS-SP1 | EWS-TAF15 | SP1-SP1 | SP1-TAF15 | TAF15-TAF15 |
|---|---|---|---|---|---|---|
| (100, 0, 0) | 0.025 | 0.000 | 0.000 | 0.000 | 0.000 | 0.000 |
| (0, 100, 0) | 0.000 | 0.000 | 0.000 | 0.016 | 0.000 | 0.000 |
| (0, 0, 100) | 0.000 | 0.000 | 0.000 | 0.000 | 0.000 | 0.014 |
| (50, 50, 0) | 0.035 | 0.035 | 0.000 | 0.020 | 0.000 | 0.000 |
| (50, 0, 50) | 0.043 | 0.000 | 0.019 | 0.000 | 0.000 | 0.020 |
| (0, 50, 50) | 0.000 | 0.000 | 0.000 | 0.029 | 0.000 | 0.026 |
| (33, 33, 33) | 0.053 | 0.042 | 0.006 | 0.034 | 0.000 | 0.034 |

Table S14: Number of interactions between species for different compositions at a temperature of 300K and 150 mM ion concentration, normalised by $N_1L_1N_2L_2$, where $N_1$, $N_2$ are the number of copies of species 1 and species 2 in the system, and $L_1$, $L_2$ are the lengths of species 1 and species 2, respectively.

| Composition (% SP2, % TAF15, % SP1) | SP1-SP1 | SP1-SP2 | SP1-TAF15 | SP2-SP2 | SP2-TAF15 | TAF15-TAF15 |
|---|---|---|---|---|---|---|
| (0, 0, 100) | 0.016 | 0.000 | 0.000 | 0.000 | 0.000 | 0.000 |
| (100, 0, 0) | 0.000 | 0.000 | 0.000 | 0.015 | 0.000 | 0.000 |
| (0, 100, 0) | 0.000 | 0.000 | 0.000 | 0.000 | 0.000 | 0.014 |
| (50, 0, 50) | 0.013 | 0.046 | 0.000 | 0.018 | 0.000 | 0.000 |
| (0, 50, 50) | 0.029 | 0.000 | 0.000 | 0.000 | 0.000 | 0.026 |
| (50, 50, 0) | 0.000 | 0.000 | 0.000 | 0.026 | 0.031 | 0.023 |
| (33, 33, 33) | 0.024 | 0.048 | 0.003 | 0.025 | 0.041 | 0.036 |

Table S15: Number of interactions between species for different compositions at a temperature of 300K and 150 mM ion concentration, normalised by $N_1L_1N_2L_2$, where $N_1$, $N_2$ are the number of copies of species 1 and species 2 in the system, and $L_1$, $L_2$ are the lengths of species 1 and species 2, respectively.

| Composition (% FUS, % TAF15, % SP2) | FUS-FUS | FUS-SP2 | FUS-TAF15 | SP2-SP2 | SP2-TAF15 | TAF15-TAF15 |
|---|---|---|---|---|---|---|
| (100, 0, 0) | 0.015 | 0.000 | 0.000 | 0.000 | 0.000 | 0.000 |
| (0, 0, 100) | 0.000 | 0.000 | 0.000 | 0.015 | 0.000 | 0.000 |
| (0, 100, 0) | 0.000 | 0.000 | 0.000 | 0.000 | 0.000 | 0.014 |
| (50, 0, 50) | 0.017 | 0.051 | 0.000 | 0.020 | 0.000 | 0.000 |
| (50, 50, 0) | 0.017 | 0.000 | 0.034 | 0.000 | 0.000 | 0.017 |
| (0, 50, 50) | 0.000 | 0.000 | 0.000 | 0.026 | 0.031 | 0.023 |
| (33, 33, 33) | 0.017 | 0.060 | 0.033 | 0.026 | 0.030 | 0.027 |

Table S16: Number of interactions between species for different compositions at a temperature of 300K and 150 mM ion concentration, normalised by $N_1L_1N_2L_2$, where $N_1$, $N_2$ are the number of copies of species 1 and species 2 in the system, and $L_1$, $L_2$ are the lengths of species 1 and species 2, respectively.

| Composition (% FUS, % TAF15, % HNF1A) | FUS-FUS | FUS-HNF1A | FUS-TAF15 | HNF1A-HNF1A | HNF1A-TAF15 | TAF15-TAF15 |
|---|---|---|---|---|---|---|
| (100, 0, 0) | 0.015 | 0.000 | 0.000 | 0.000 | 0.000 | 0.000 |
| (0, 0, 100) | 0.000 | 0.000 | 0.000 | 0.021 | 0.000 | 0.000 |
| (0, 100, 0) | 0.000 | 0.000 | 0.000 | 0.000 | 0.000 | 0.014 |
| (50, 0, 50) | 0.015 | 0.042 | 0.000 | 0.030 | 0.000 | 0.000 |
| (50, 50, 0) | 0.017 | 0.000 | 0.034 | 0.000 | 0.000 | 0.017 |
| (0, 50, 50) | 0.000 | 0.000 | 0.000 | 0.039 | 0.008 | 0.024 |
| (33, 33, 33) | 0.018 | 0.053 | 0.020 | 0.046 | 0.003 | 0.030 |

Table S17: Number of interactions between species for different compositions at a temperature of 300K and 150 mM ion concentration, normalised by $N_1L_1N_2L_2$, where $N_1$, $N_2$ are the number of copies of species 1 and species 2 in the system, and $L_1$, $L_2$ are the lengths of species 1 and species 2, respectively.

| Composition (% HNF1A, % TAF15, % SP1) | HNF1A-HNF1A | HNF1A-SP1 | HNF1A-TAF15 | SP1-SP1 | SP1-TAF15 | TAF15-TAF15 |
|---|---|---|---|---|---|---|
| (100, 0, 0) | 0.021 | 0.000 | 0.000 | 0.000 | 0.000 | 0.000 |
| (0, 0, 100) | 0.000 | 0.000 | 0.000 | 0.016 | 0.000 | 0.000 |
| (0, 100, 0) | 0.000 | 0.000 | 0.000 | 0.000 | 0.000 | 0.014 |
| (50, 0, 50) | 0.026 | 0.050 | 0.000 | 0.013 | 0.000 | 0.000 |
| (50, 50, 0) | 0.039 | 0.000 | 0.008 | 0.000 | 0.000 | 0.024 |
| (0, 50, 50) | 0.000 | 0.000 | 0.000 | 0.029 | 0.000 | 0.026 |
| (33, 33, 33) | 0.038 | 0.072 | 0.005 | 0.020 | 0.002 | 0.035 |

Table S18: Number of interactions between species for different compositions at a temperature of 300K and 150 mM ion concentration, normalised by $N_1L_1N_2L_2$, where $N_1$, $N_2$ are the number of copies of species 1 and species 2 in the system, and $L_1$, $L_2$ are the lengths of species 1 and species 2, respectively.

| Composition (% HNF1A, % SP1, % SP2) | HNF1A-HNF1A | HNF1A-SP1 | HNF1A-SP2 | SP1-SP1 | SP1-SP2 | SP2-SP2 |
|---|---|---|---|---|---|---|
| (100, 0, 0) | 0.021 | 0.000 | 0.000 | 0.000 | 0.000 | 0.000 |
| (0, 100, 0) | 0.000 | 0.000 | 0.000 | 0.016 | 0.000 | 0.000 |
| (0, 0, 100) | 0.000 | 0.000 | 0.000 | 0.000 | 0.000 | 0.015 |
| (50, 50, 0) | 0.026 | 0.050 | 0.000 | 0.013 | 0.000 | 0.000 |
| (50, 0, 50) | 0.023 | 0.000 | 0.052 | 0.000 | 0.000 | 0.015 |
| (0, 50, 50) | 0.000 | 0.000 | 0.000 | 0.013 | 0.046 | 0.018 |
| (33, 33, 33) | 0.025 | 0.047 | 0.063 | 0.012 | 0.043 | 0.015 |

Table S19: Number of interactions between species for different compositions at a temperature of 300K and 150 mM ion concentration, normalised by $N_1L_1N_2L_2$, where $N_1$, $N_2$ are the number of copies of species 1 and species 2 in the system, and $L_1$, $L_2$ are the lengths of species 1 and species 2, respectively.

### 6 Simulation Images

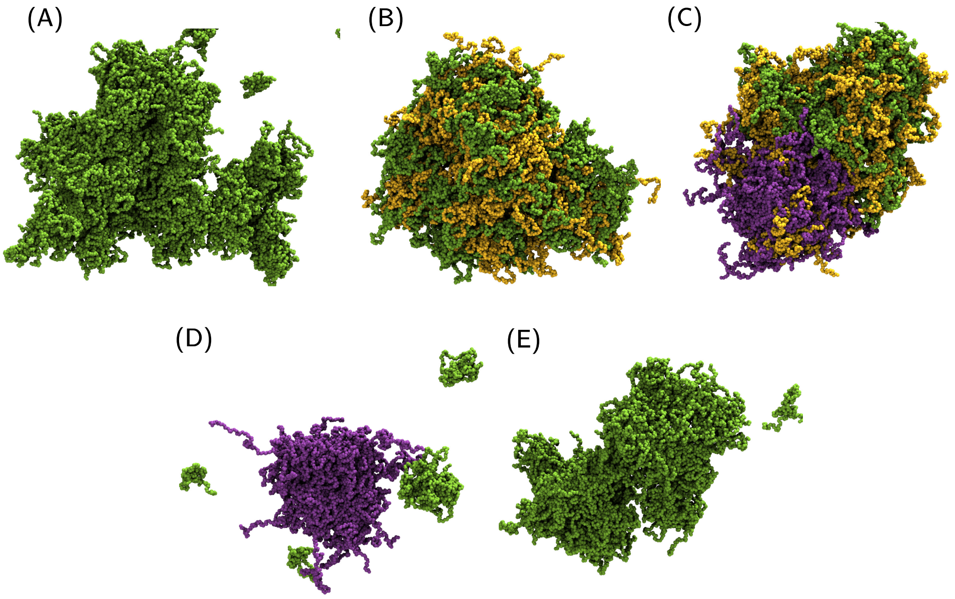

Figure S8: Snapshots of simulations completed with increased total particle number. Simulation were run using the same settings with a total amino acid concentration of 80000 $\mu M$. The total composition in these systems is defined relative to 240 FUS molecules, 240 TAF15 molecules, and 120 SP1 molecules, to give the percentage compositions (% FUS, % SP1, % TAF15). The end frame of 3 $\mu$s of simulation is displayed as a representative state for (A) (0, 0, 100), (B) (50,0,50), (C) (33, 33, 33), (D) (0, 25, 75) and (E) (0, 25, 75). (D) and (E) are from different regions of the same simulation as it requires a lower zoom to get both condensates in same image. FUS molecules are coloured in yellow, SP1 molecules are coloured in purple, and TAF15 molecules are coloured in green. Droplets are formed in all simulations of these molecules under these concentration conditions, irrespective of composition.

### 7 Contact Lifetimes

Contact lifetimes between residues have been computed for the different species using the double cutoff approach developed in [22]. The short cutoff of 0.6 nm for the formation of contacts and a long cutoff of 1.0 nm for the breaking of contacts were used in the analysis of a 5 ns simulation for different compositions, as shown in Figure S9. From Figure S9 we show that the contact lifetimes are fast, with peaks under 5ps (simulation time) for all interaction types.

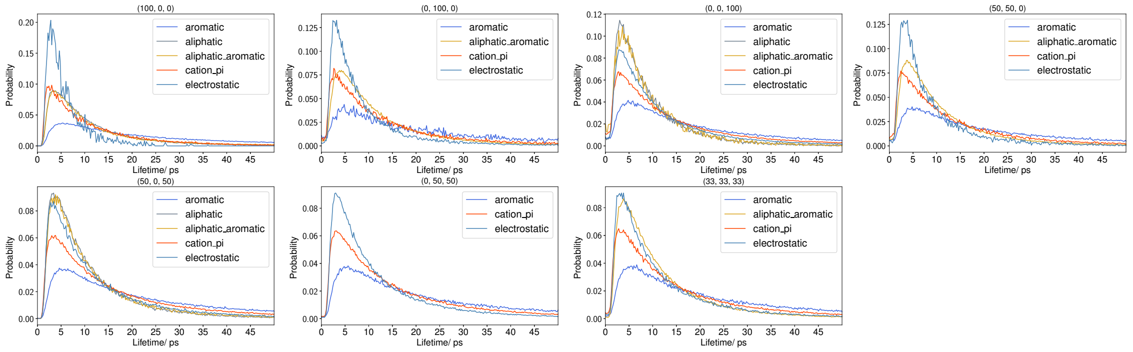

Figure S9: Contact lifetimes in simulation time for simulations of different compositions. The total composition in these systems is defined relative to 120 FUS molecules, 120 TAF15 molecules, and 60 SP1 molecules, to give the percentage compositions (% FUS, % SP1, % TAF15). Based on the coarse grained simulation settings the speed up factor (1000x) [12] results in contact lifetimes on the order of 5ns.

### 8 Additional ternary systems

Simulation outputs for ternary systems detailed in Table 1 in the main manuscript are in Figures S10-S15.

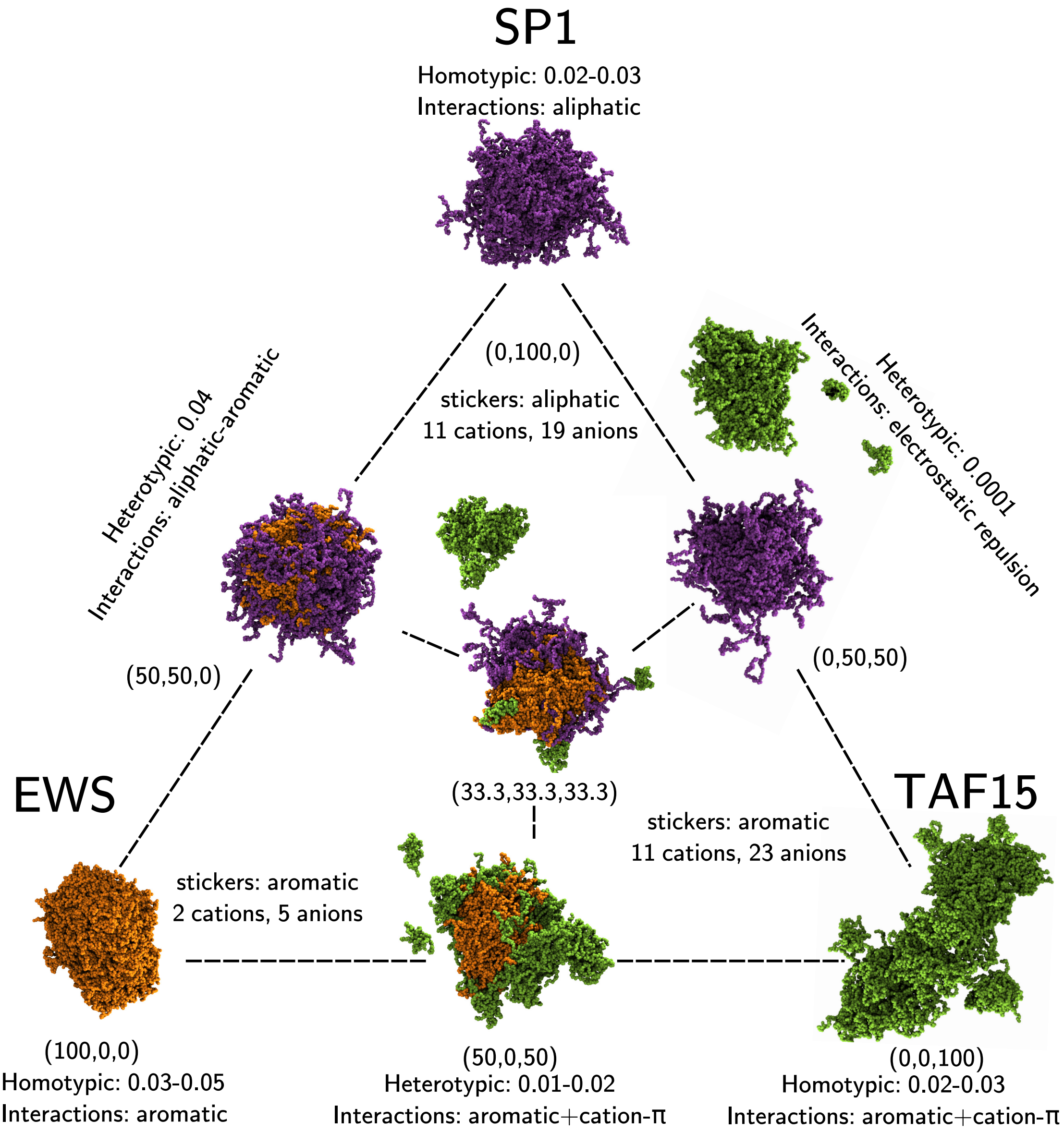

Figure S10: **Snapshots and interaction data of the ternary EWS-SP1-TAF15 system.** The total composition is defined relative to 120 EWS molecules, 120 TAF15 molecules, and 60 SP1 molecules, to give the percentage compositions (% EWS, % SP1, % TAF15). The end frame of 3 $\mu$s of simulation is displayed as a representative state. EWS molecules are coloured in orange, SP1 molecules are coloured in purple, and TAF15 molecules are coloured in green. Droplets are formed in all simulations of these molecules under these concentration conditions, irrespective of composition.

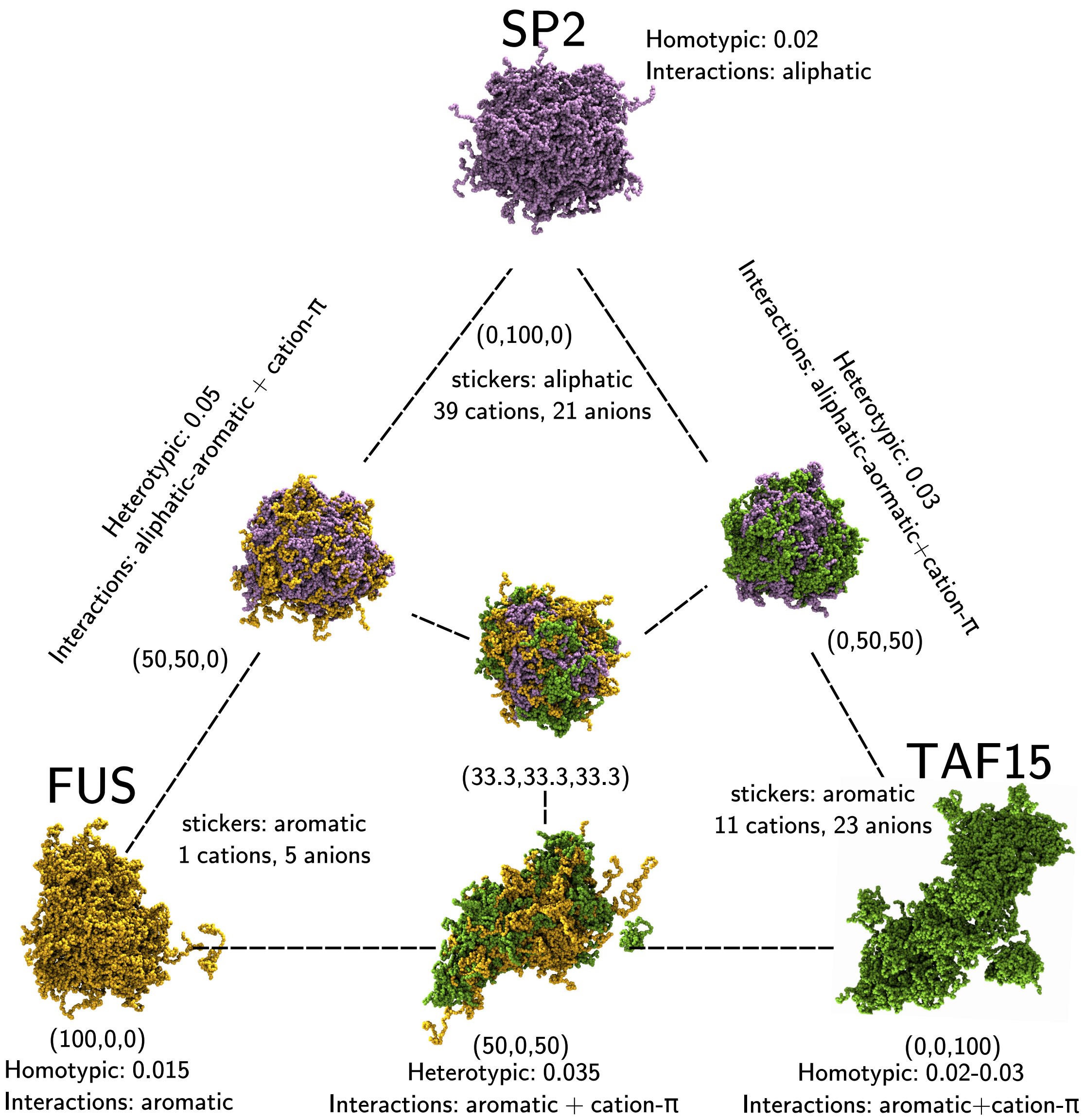

Figure S11: **Snapshots and interaction data of the ternary FUS-SP2-TAF15 system.** The total composition is defined relative to 120 FUS molecules, 120 TAF15 molecules, and 60 SP2 molecules, to give the percentage compositions (% FUS, % SP2, % TAF15). The end frame of 3 $\mu$s of simulation is displayed as a representative state. FUS molecules are coloured in yellow, SP2 molecules are coloured in light purple, and TAF15 molecules are coloured in green. Droplets are formed in all simulations of these molecules under these concentration conditions, irrespective of composition.

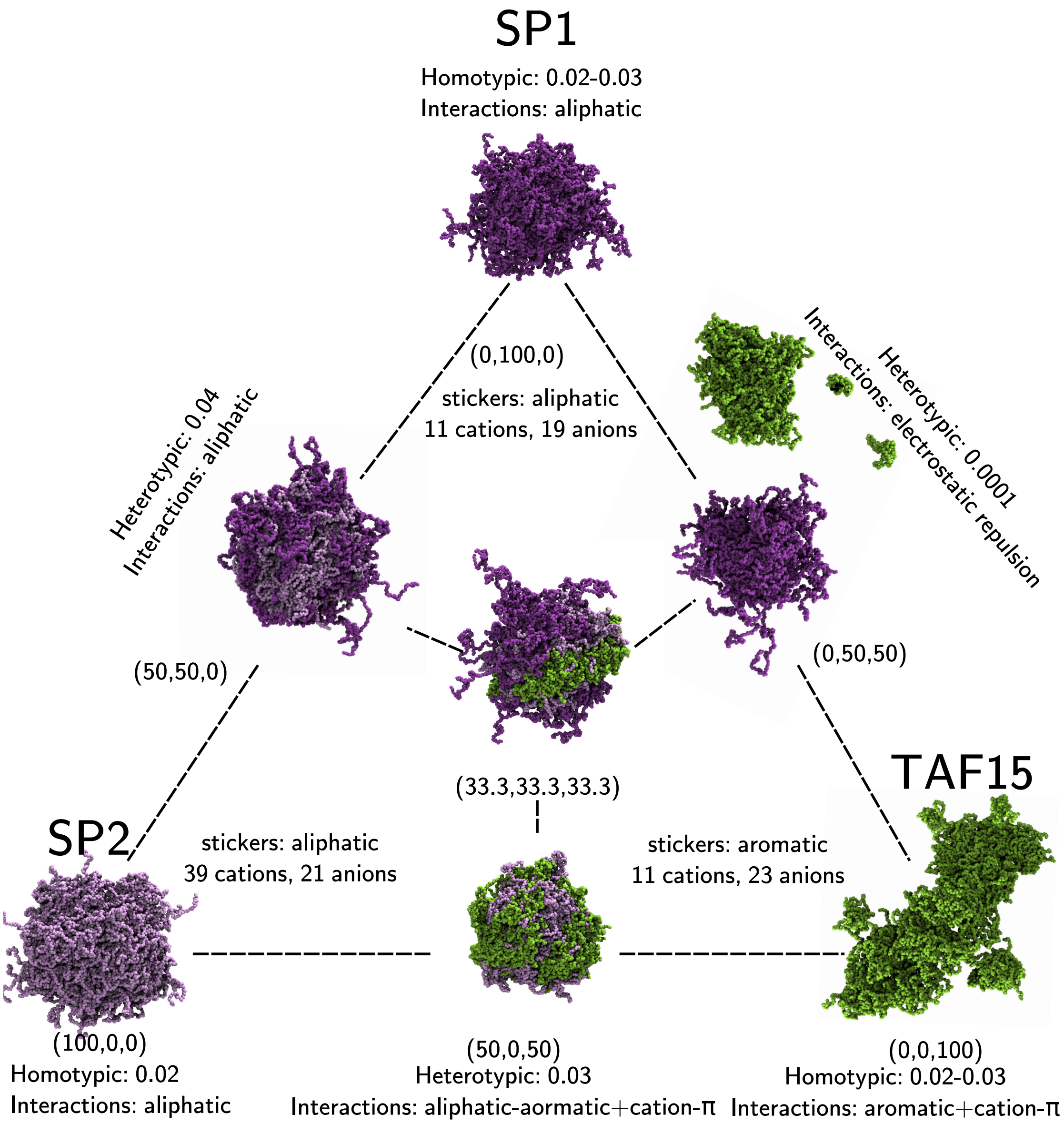

Figure S12: **Snapshots and interaction data of the ternary SP1-SP2-TAF15 system.** The total composition is defined relative to 60 SP1 molecules, 120 TAF15 molecules, and 60 SP2 molecules, to give the percentage compositions (% SP2, % SP1, % TAF15). The end frame of 3 $\mu$s of simulation is displayed as a representative state. SP2 molecules are coloured in light purple, SP1 molecules are coloured in purple, and TAF15 molecules are coloured in green. Droplets are formed in all simulations of these molecules under these concentration conditions, irrespective of composition.

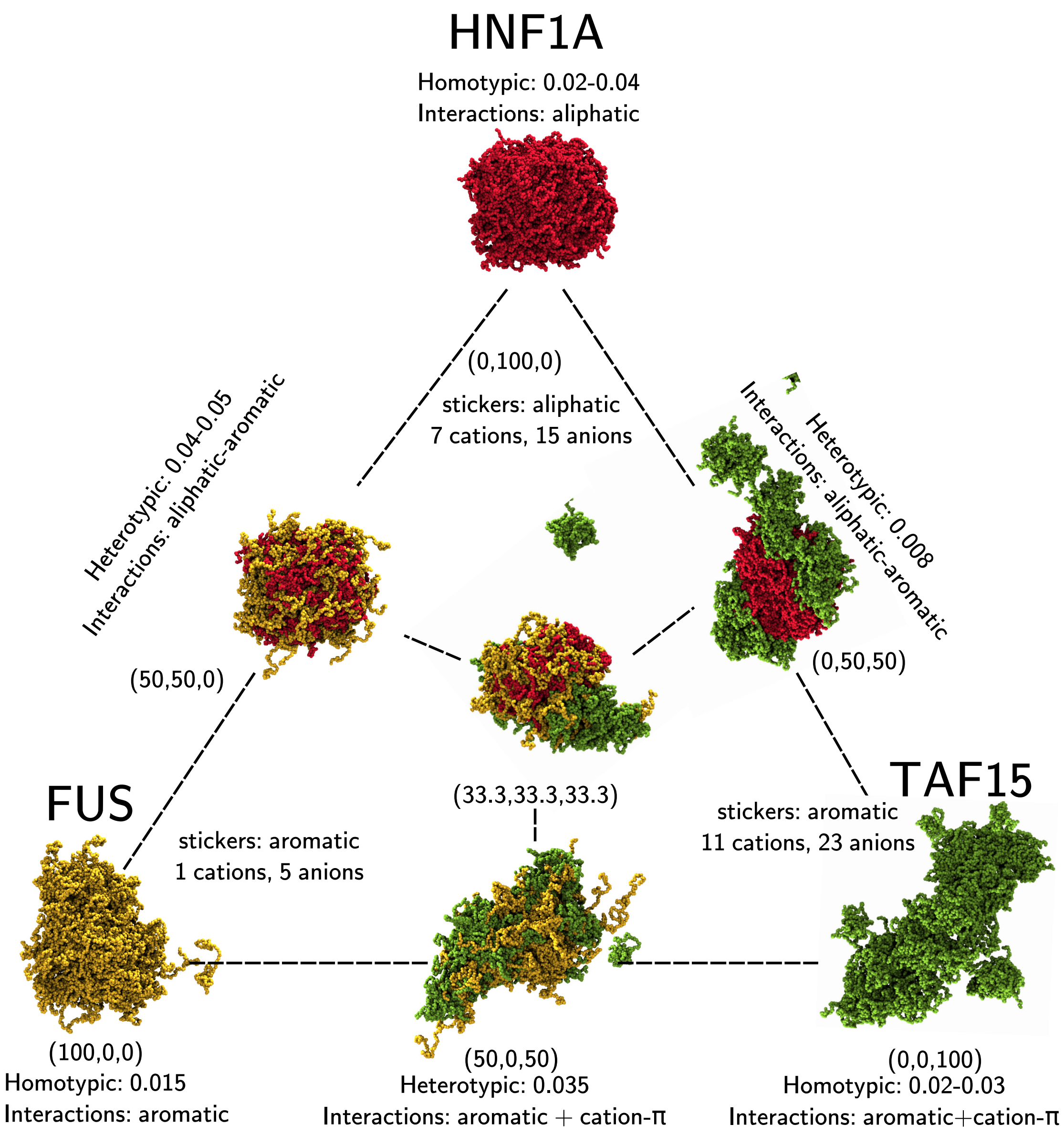

Figure S13: **Snapshots and interaction data of the ternary FUS-HNF1A-TAF15 system.** The total composition is defined relative to 120 FUS molecules, 120 TAF15 molecules, and 90 HNF1A molecules, to give the percentage compositions (% FUS, % HNF1A, % TAF15). The end frame of 3 $\mu$s of simulation is displayed as a representative state. FUS molecules are coloured in yellow, HNF1A molecules are coloured in red, and TAF15 molecules are coloured in green. Droplets are formed in all simulations of these molecules under these concentration conditions, irrespective of composition.

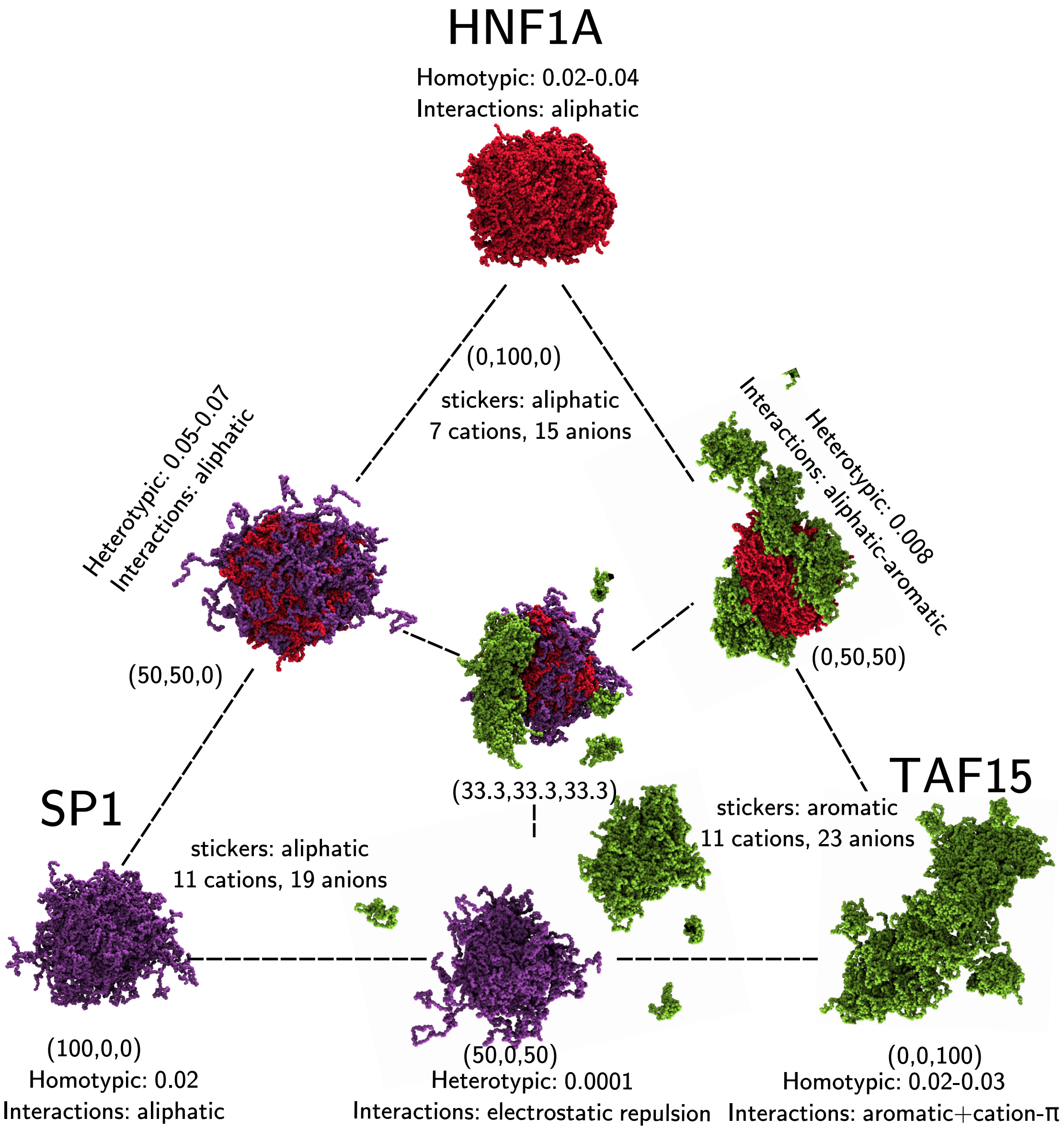

Figure S14: **Snapshots and interaction data of the ternary SP1-HNF1A-TAF15 wsystem.** The total composition is defined relative to 60 SP1 molecules, 120 TAF15 molecules, and 90 HNF1A molecules, to give the percentage compositions (% SP1, % HNF1A, % TAF15). The end frame of 3 $\mu$s of simulation is displayed as a representative state. HNF1A molecules are coloured in red, SP1 molecules are coloured in purple, and TAF15 molecules are coloured in green. Droplets are formed in all simulations of these molecules under these concentration conditions, irrespective of composition.

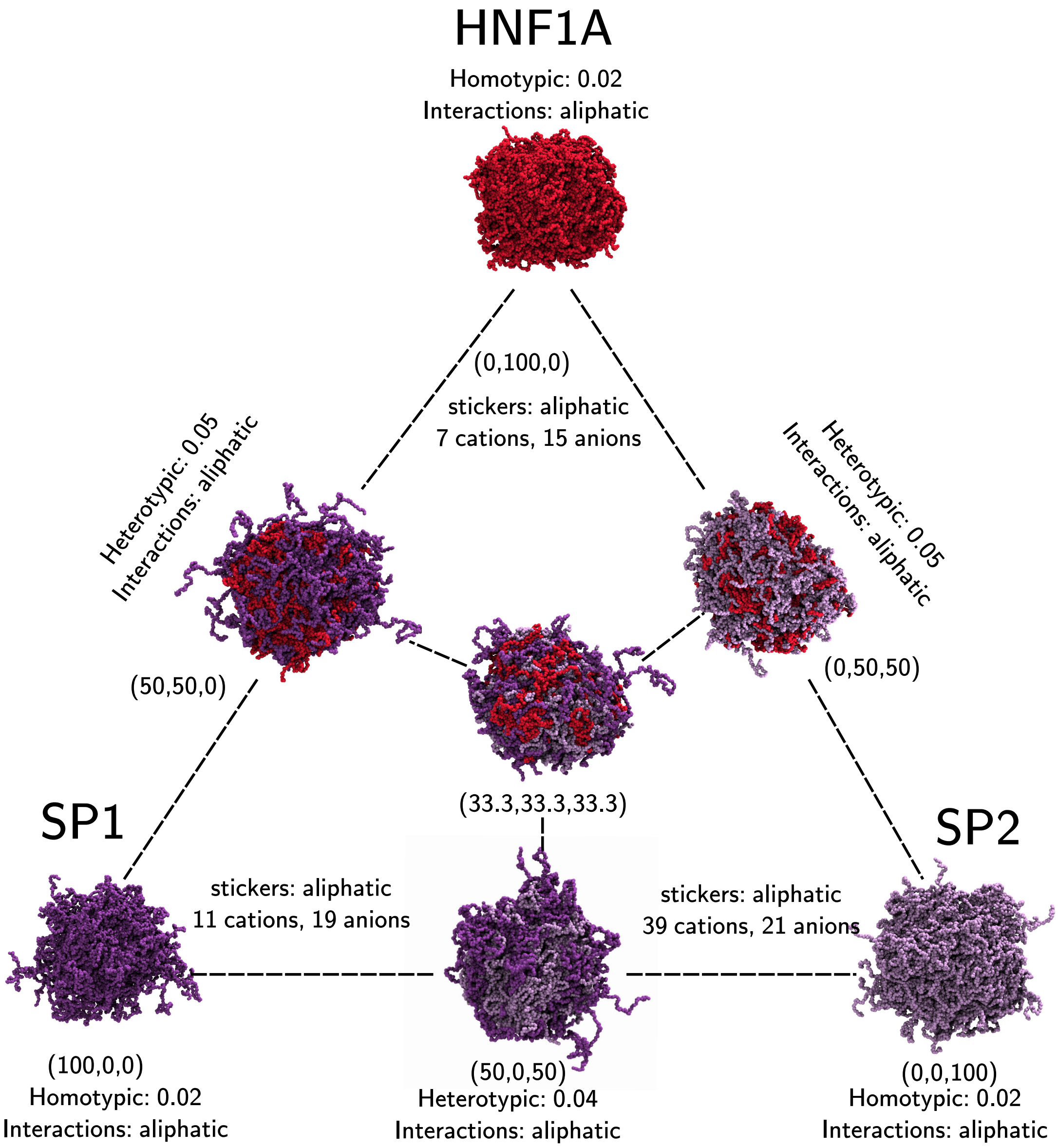

Figure S15: **Snapshots and interaction data of the ternary SP1-HNF1A-SP2 system.** The total composition is defined relative to 60 SP2 molecules, 90 HNF1A molecules, and 60 SP1 molecules, to give the percentage compositions (% SP1, % HNF1A, % SP2). The end frame of 3 $\mu$s of simulation is displayed as a representative state. SP2 molecules are coloured in light purple, SP1 molecules are coloured in purple, and HNF1A molecules are coloured in red. Droplets are formed in all simulations of these molecules under these concentration conditions, irrespective of composition.

### 9 Contact data

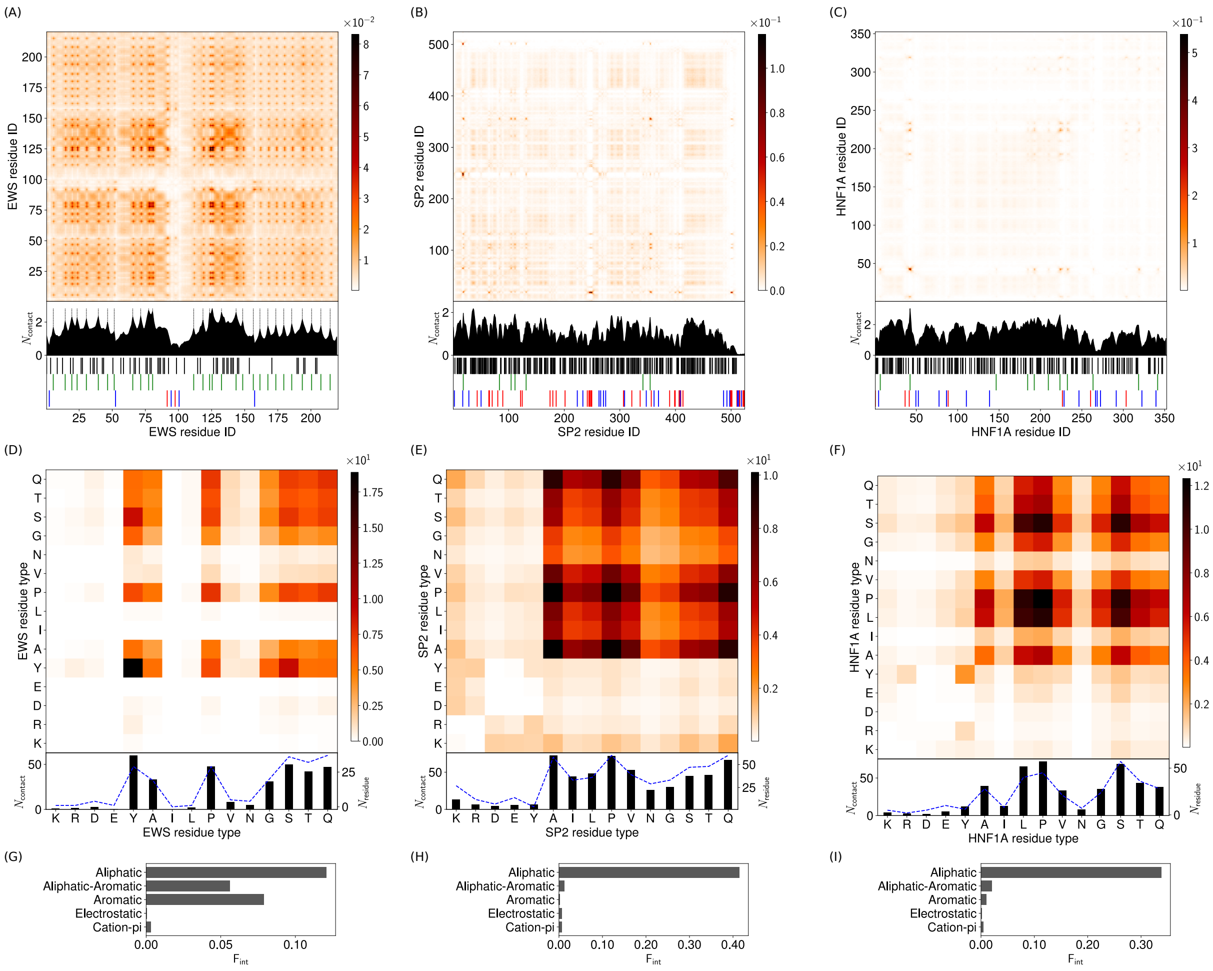

Figure S16: **Intermolecular contact maps for homotypic interactions in single-component droplets.** (A)-(C) Intermolecular contacts by residue index for (A) 100% EWS, (B) 100% SP2, and (C) 100% HNF1A, at 150 mM ion concentration and 300 K. The contacts are averaged in time and normalised by the number of molecules in the simulation (see section 3.2 for more details). A 1D contact profile (summation of the 2D map) is included below the contact map to show the total interactions per residue index ($N_{\text{contact}}$). The black dashed lines in (A) highlight the residues with the most contacts in EWS (which correspond to peaks in the 1D profiles). These are the aromatic residues in EWS driving $\pi$-$\pi$ aromatic contacts. Broad peaks are seen in (B) and (C) corresponding to the aliphatic residue stretches in SP1 and HNF1A, respectively. (D)-(F) Intermolecular contact map by residue type for (D) 100% EWS, (E) 100% SP2, and (F) 100% HNF1A. The contact maps in (D), (E) and (F) are similar to the contact maps by residue index in (A), (B), and (C), respectively, but aggregated by residue type. A 1D contact profile (summation of the 2D map), are also included below ($N_{\text{contact}}$) together with the abundance for the residues ($N_{\text{residue}}$) shown by blue dashed lines. (G)-(I) Intermolecular interaction summary for (G) EWS in 100% EWS (H) SP2 in 100% SP2, and (I) HNF1A in 100% HNF1A at 150 mM and 300 K. The fraction of interactions, $\mathsf{F}_{\text{int}}$, are aggregated by type and normalised by the total number of the intermolecular interactions in (A)-(C) respectively. Aromatic and aliphatic interactions denote aromatic-aromatic and aliphatic-aliphatic interactions respectively. This convention is used throughout this work. Details of the contact definitions can be found in section 3.2 of the SI.

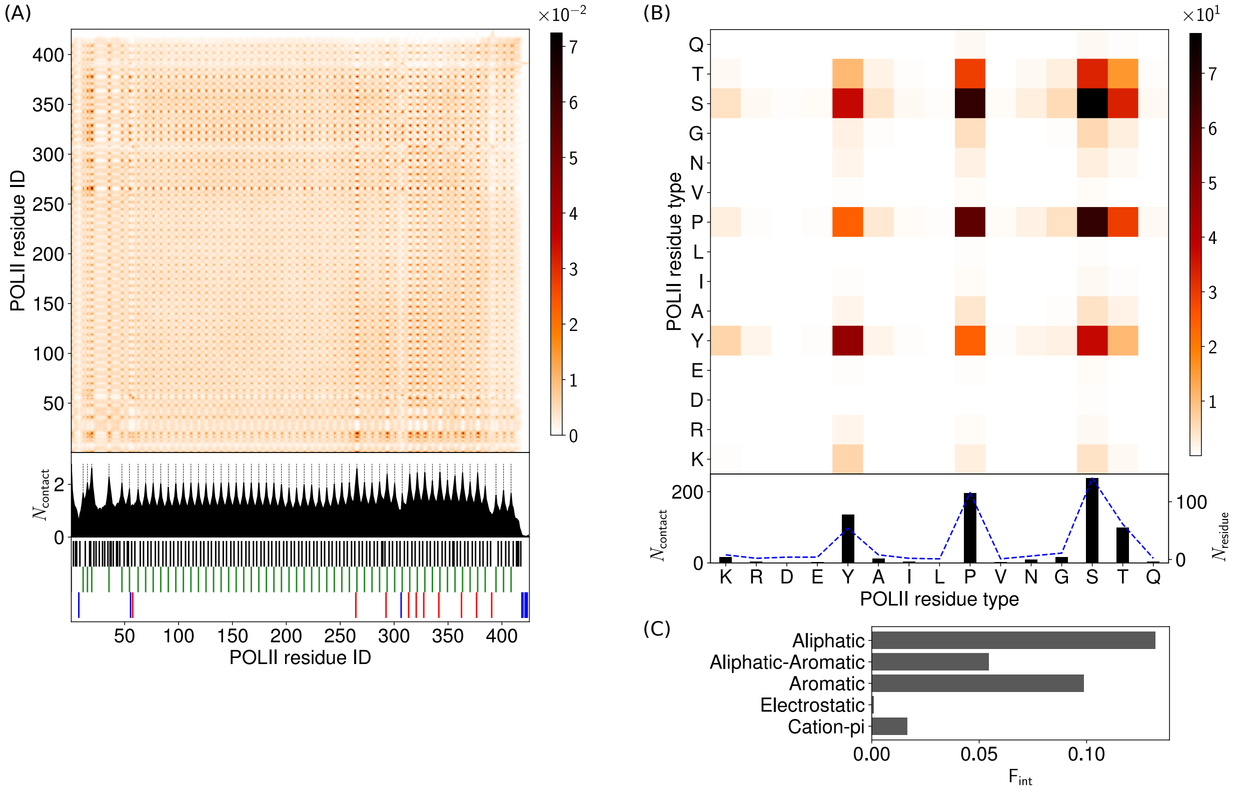

Figure S17: **Intermolecular contact maps for homotypic interactions in single-component POL II droplet.** (A) Intermolecular contacts by residue index for 100% POL II at 150 mM ion concentration and 300 K. The contacts are averaged in time and normalised by the number of molecules in the simulation (see section 3.2 for more details). A 1D contact profile (summation of the 2D map) is included below the contact map to show the total interactions per residue index ($N_{\text{contact}}$). The black dashed lines in (A) highlight the residues with the most contacts in POL II (which correspond to peaks in the 1D profiles). These are the aromatic residues in POL II driving $\pi$-$\pi$ aromatic contacts. (B) Intermolecular contact map by residue type for 100% POL II. The contact map in (B) are similar to the contact maps by residue index in (A) but aggregated by residue type. A 1D contact profile (summation of the 2D map), are also included below ($N_{\text{contact}}$) together with the abundance for the residues ($N_{\text{residue}}$) shown by blue dashed lines. (C) Intermolecular interaction summary for POL II in 100% POL II at 150 mM and 300 K. The fraction of interactions, $\mathsf{F}_{\text{int}}$, are aggregated by type and normalised by the total number of the intermolecular interactions in (A) respectively. Aromatic and aliphatic interactions denote aromatic-aromatic and aliphatic-aliphatic interactions respectively. This convention is used throughout this work. Details of the contact definitions can be found in section 3.2 of the SI.

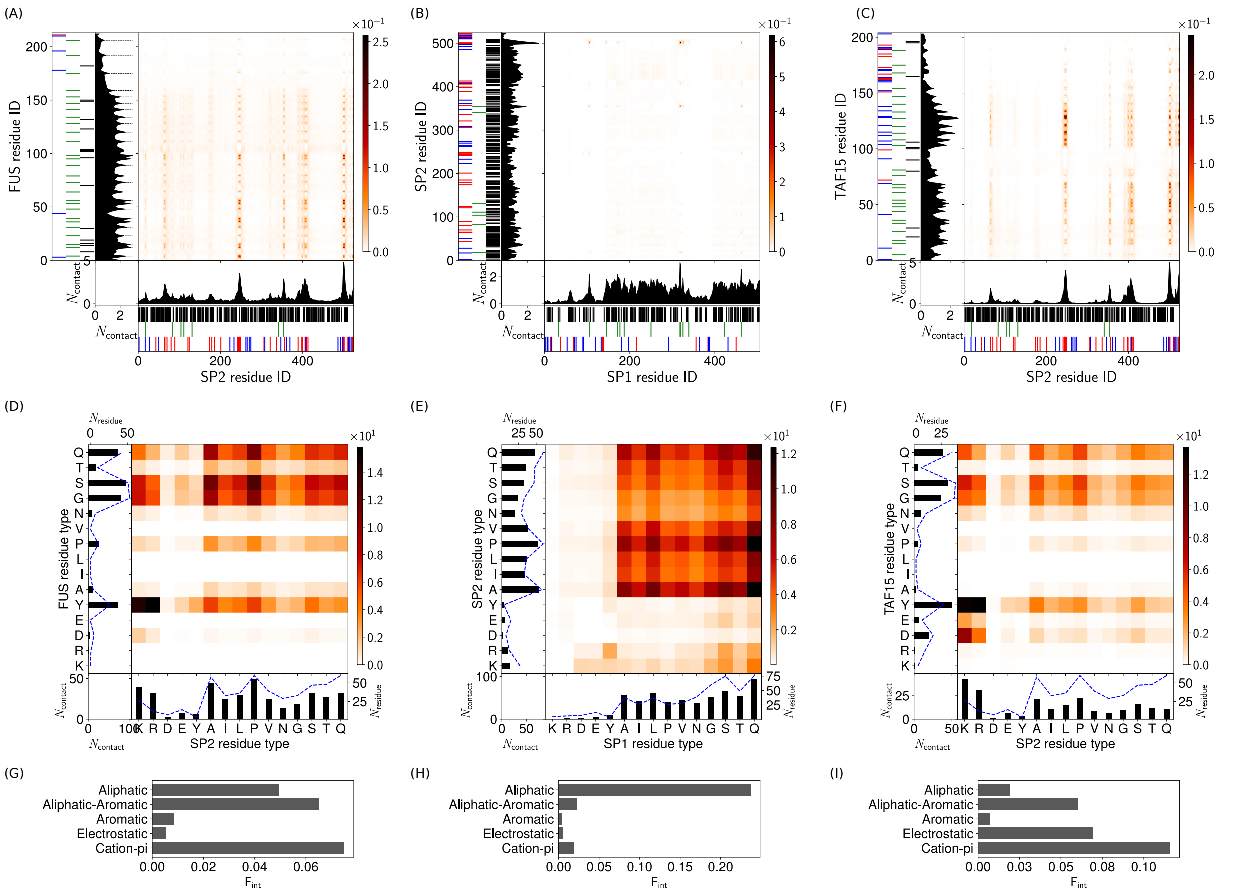

Figure S18: **Intermolecular contact maps for heterotypic interactions in two-component droplets.** (A)-(C) Intermolecular contact map by residue index for (A) FUS with SP2 (50%, 50%), (B) SP1 with SP2 (50%, 50%), and (C) SP2 with TAF15 (50%, 50%) at 150 mM and 300 K. For the definitions of the different contact types see the caption of Figure 3 and section 3.2. The 1D contact profiles denote a summation of the 2D map of the corresponding molecules. The black dashed lines highlight the key residues: (A) the aromatic residues in FUS. (D)-(F) Intermolecular contact map by residue type for (D) FUS with SP2 (50%, 50%), (E) SP1 with SP2 (50%, 50%), and (F) SP2 with TAF15 (50%, 50%) at 150 mM and 300 K. (G)-(I) Intermolecular interaction summary for (G) FUS-SP2 interactions in (50%, 50%), (H) SP1-SP2 interactions in (50%, 50%), and (I) TAF15-SP2 interactions in (50%, 50%) at 150 mM and 300 K. The fraction of interactions, $\mathsf{F}_{\mathsf{int}}$, are aggregated by type and normalised by the total number of the intermolecular interactions in (A)-(C) respectively.

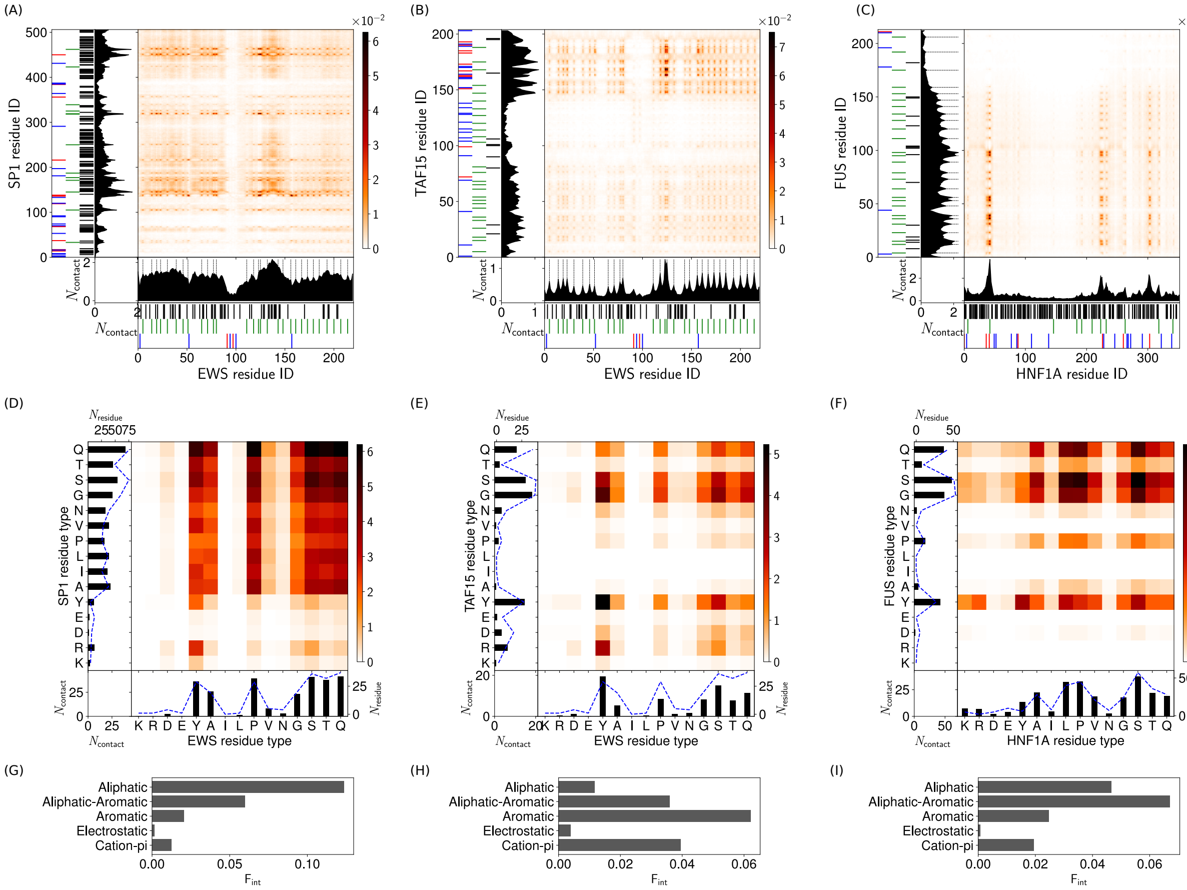

Figure S19: **Intermolecular contact maps for heterotypic interactions in two-component droplets.** (A)-(C) Intermolecular contact map by residue index for (A) EWS with SP1 (50%, 50%), (B) EWS with TAF15 (50%, 50%), and (C) FUS with HNF1A (50%, 50%) at 150 mM and 300 K. For the definitions of the different contact types see the caption of Figure 3 and section 3.2. The 1D contact profiles denote a summation of the 2D map of the corresponding molecules. The black dashed lines highlight the key residues: (A) the aromatic residues in EWS, (B) the aromatic residues in EWS, and (C) the aromatic residues in FUS. (D)-(F) Intermolecular contact map by residue type for (D) EWS with SP1 (50%, 50%), (E) EWS with TAF15 (50%, 50%), and (F) FUS with HNF1A (50%, 50%) at 150 mM and 300 K. (G)-(I) Intermolecular interaction summary for (G) EWS-SP1 interactions in (50%, 50%), (H) EWS-TAF15 interactions in (50%, 50%), and (I) FUS-HNF1A interactions in (50%, 50%) at 150 mM and 300 K. The fraction of interactions, $\mathsf{F}_{\mathsf{int}}$, are aggregated by type and normalised by the total number of the intermolecular interactions in (A)-(C) respectively.

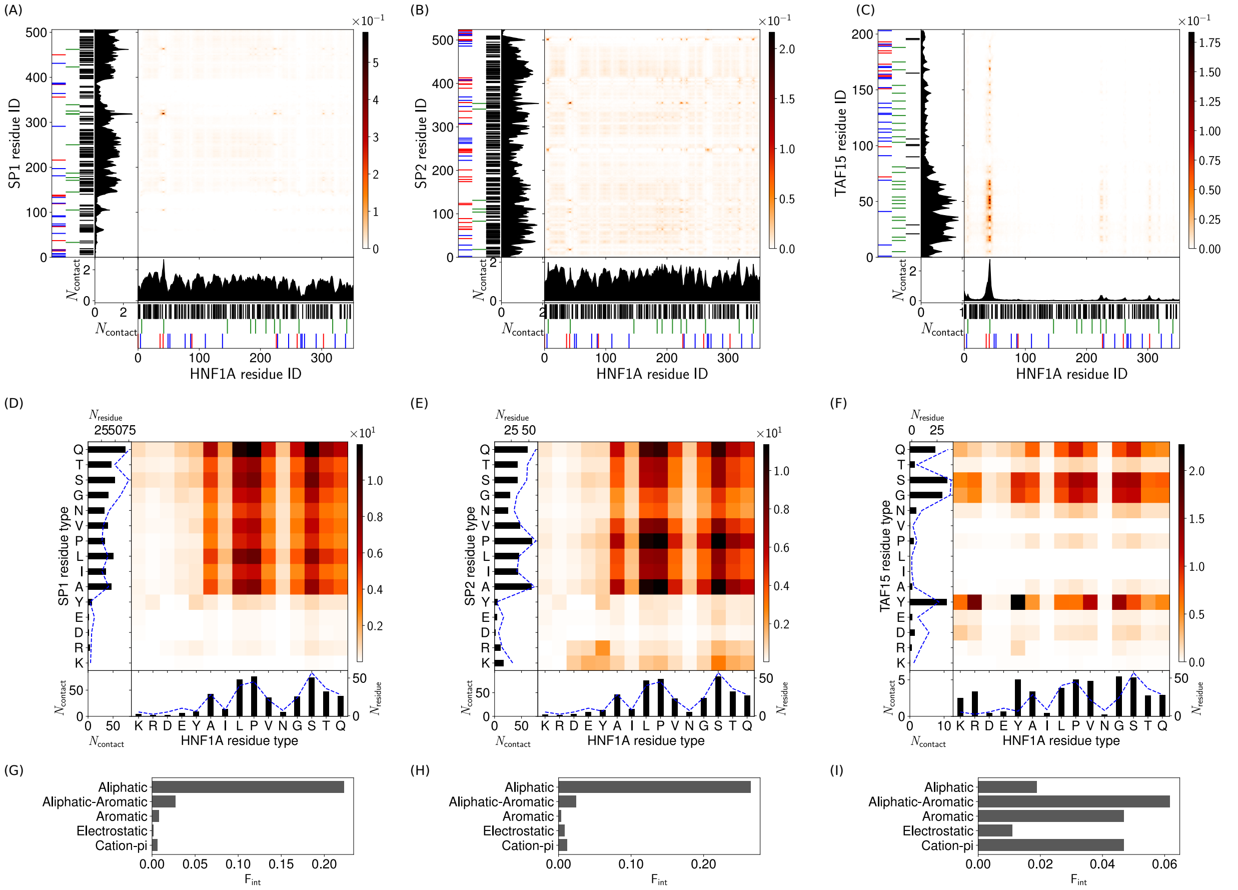

Figure S20: **Intermolecular contact maps for heterotypic interactions in two-component droplets.** (A)-(C) Intermolecular contact map by residue index for (A) HNF1A with SP1 (50%, 50%), (B) HNF1A with SP2 (50%, 50%), and (C) HNF1A with TAF15 (50%, 50%) at 150 mM and 300 K. For the definitions of the different contact types see the caption of Figure 3 and section 3.2. The 1D contact profiles denote a summation of the 2D map of the corresponding molecules. (D)-(F) Intermolecular contact map by residue type for (D) HNF1A with SP1 (50%, 50%), (E) HNF1A with SP2 (50%, 50%), and (F) HNF1A with TAF15 (50%, 50%) at 150 mM and 300 K. (G)-(I) Intermolecular interaction summary for (G) HNF1A-SP1 interactions in (50%, 50%), (H) HNF1A-SP2 interactions in (50%, 50%), and (I) HNF1A-TAF15 interactions in (50%, 50%) at 150 mM and 300 K. The fraction of interactions, $\mathsf{F_{int}}$, are aggregated by type and normalised by the total number of the intermolecular interactions in (A)-(C) respectively.

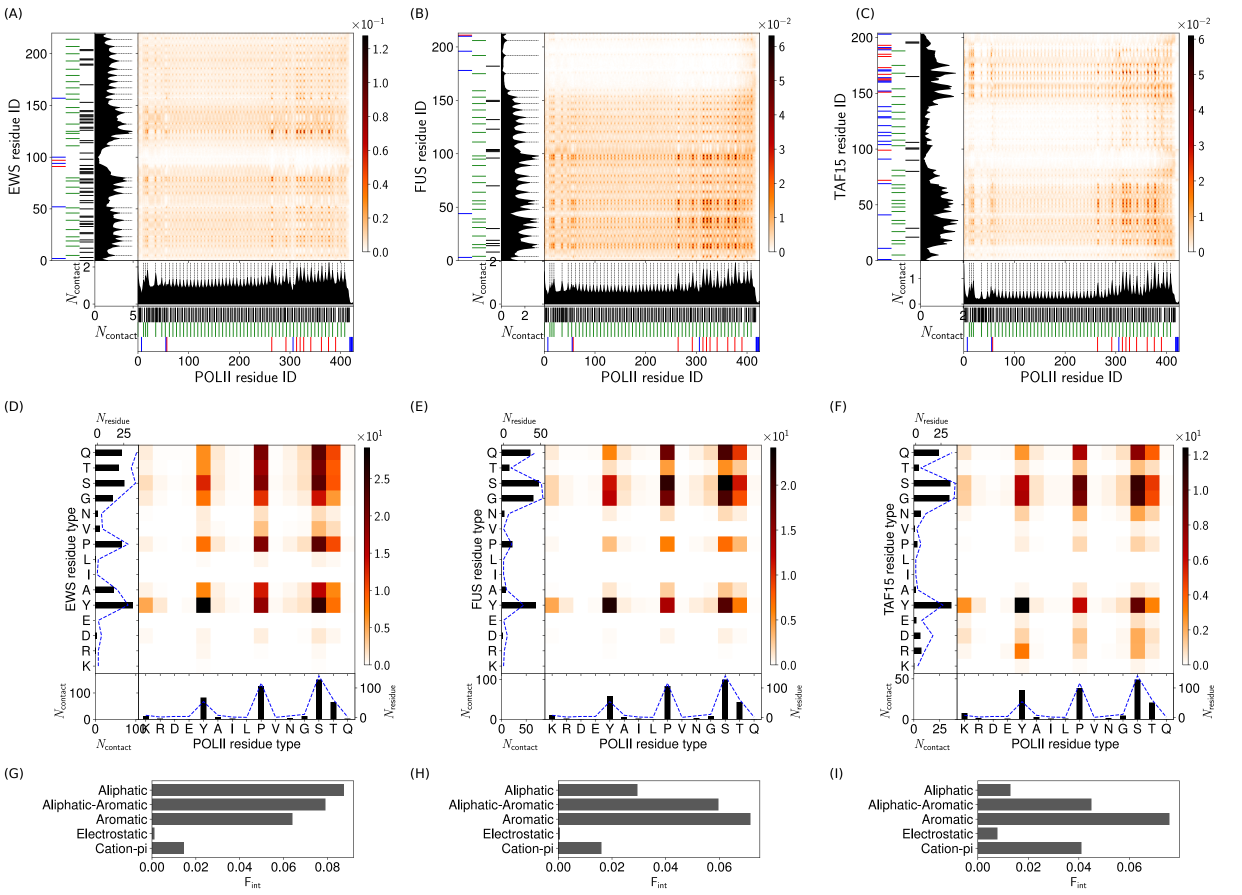

Figure S21: **Intermolecular contact maps for heterotypic interactions in two-component droplets.** (A)-(C) Intermolecular contact map by residue index for (A) EWS with POL II (50%, 50%), (B) FUS with POL II (50%, 50%), and (C) TAF15 with POL II (50%, 50%) at 150 mM and 300 K. For the definitions of the different contact types see the caption of Figure 3 and section 3.2. The 1D contact profiles denote a summation of the 2D map of the corresponding molecules. The black dashed lines highlight the key residues: (A) the aromatic residues in EWS and POL II, (B) the aromatic residues in FUS and POL II, and (C) the aromatic residues in POL II. (D)-(F) Intermolecular contact map by residue type for (D) EWS with POL II (50%, 50%), (E) FUS with POL II (50%, 50%), and (F) TAF15 with POL II (50%, 50%) at 150 mM and 300 K. (G)-(I) Intermolecular interaction summary for (G) EWS-POL II interactions in (50%, 50%), (H) FUS-POL II interactions in (50%, 50%), and (I) TAF15-POL II interactions in (50%, 50%) at 150 mM and 300 K. The fraction of interactions, $F_{int}$, are aggregated by type and normalised by the total number of the intermolecular interactions in (A)-(C) respectively.

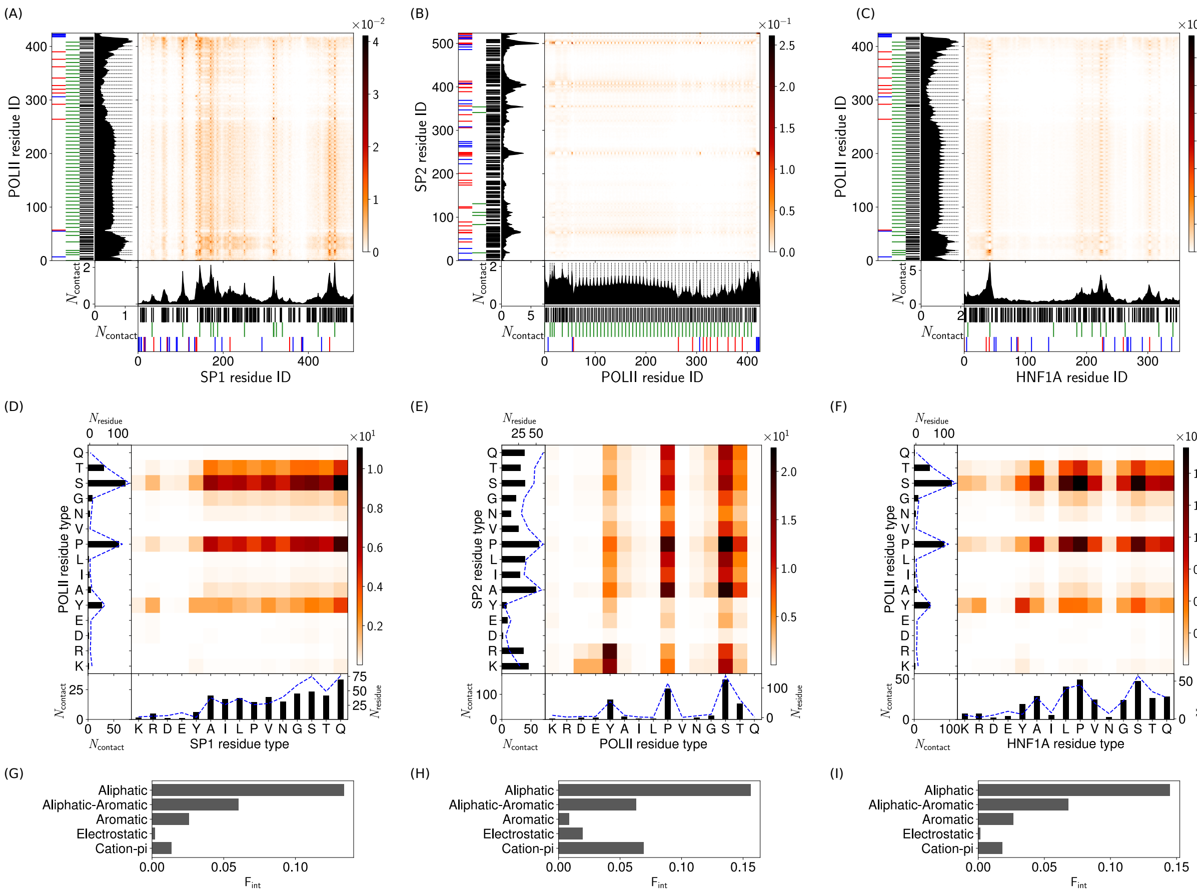

Figure S22: **Intermolecular contact maps for heterotypic interactions in two-component droplets.** (A)-(C) Intermolecular contact map by residue index for (A) SP1 with POL II (50%, 50%), (B) SP2 with POL II (50%, 50%), and (C) HNF1A with POL II (50%, 50%) at 150 mM and 300 K. For the definitions of the different contact types see the caption of Figure 3 and section 3.2. The 1D contact profiles denote a summation of the 2D map of the corresponding molecules. The black dashed lines highlight the key residues: (A) the aromatic residues in POL II, (B) the aromatic residues in POL II, and (C) the aromatic residues in POL II. (D)-(F) Intermolecular contact map by residue type for (D) SP1 with POL II (50%, 50%), (E) SP2 with POL II (50%, 50%), and (F) HNF1A with POL II (50%, 50%) at 150 mM and 300 K. (G)-(I) Intermolecular interaction summary for (G) SP1-POL II interactions in (50%, 50%), (H) SP2-POL II interactions in (50%, 50%), and (I) HNF1A-POL II interactions in (50%, 50%) at 150 mM and 300 K. The fraction of interactions, $\mathsf{F}_{\mathsf{int}}$, are aggregated by type and normalised by the total number of the intermolecular interactions in (A)-(C) respectively.

### 10 Simulation data on all TFs

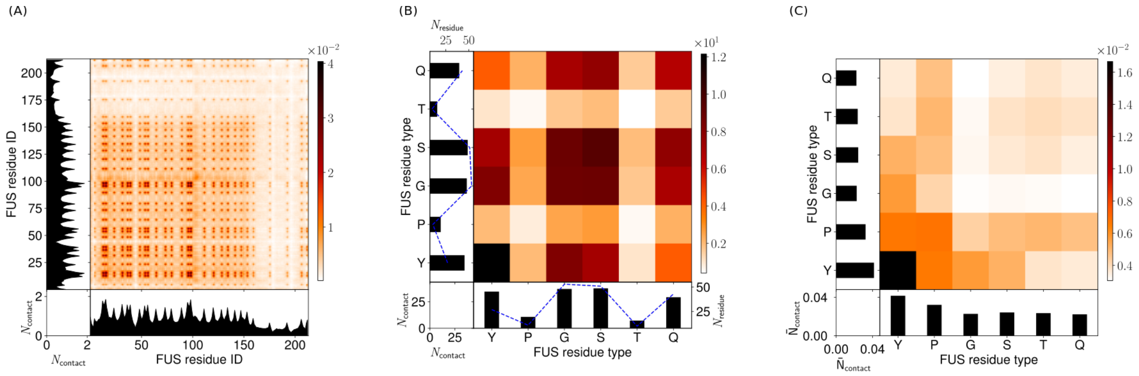

Figure S23: Contact maps for a droplet simulation with 120 FUS molecules (100%, 0%, 0%) at 150 mM and 300 K. (A) Intermolecular contact map by particle index for FUS. (B) Intermolecular contact map by particle type for FUS. (C) Intermolecular contact map by particle type, normalised by particle abundance for FUS. The contacts in the contact maps by particle index (left column) are normalised by the number of frames used (600) and $(N_{mol1}N_{mol2})^{0.5}$ where $N_{mol1}$, $N_{mol2}$ are the number of copies of molecule 1 and molecule 2. The contact maps aggregated by particle type (centre column) are a matrix reduction of the contact map by particle index. Bead abundance in a molecule is shown by a blue dashed line. The contact maps (right column) are normalised by the square root of the relative bead abundance in the molecules.

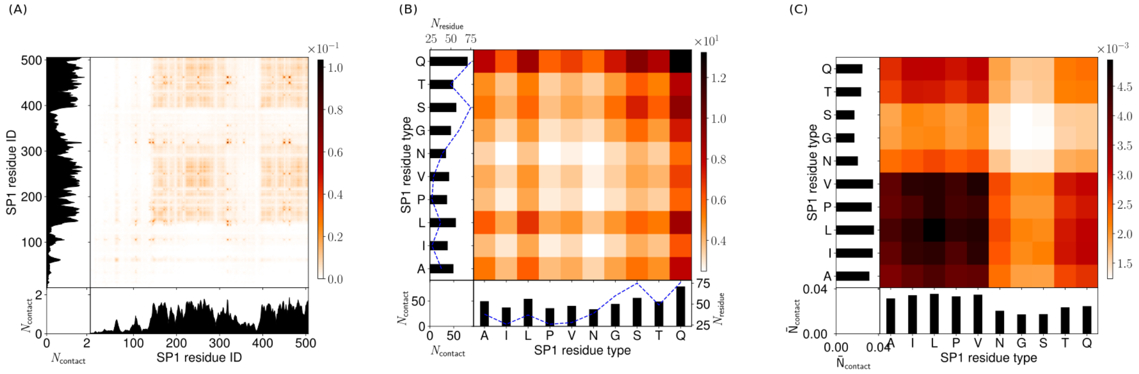

Figure S24: Contact maps for a droplet simulation with 60 SP1 molecules (0%, 100%, 0%) at 150 mM and 300 K. (A) Intermolecular contact map by particle index for SP1. (B) Intermolecular contact map by particle type for SP1. (C) Intermolecular contact map by particle type, normalised by particle abundance for SP1. The contacts in the contact maps by particle index (left column) are normalised by the number of frames used (600) and $(N_{mol1}N_{mol2})^{0.5}$ where $N_{mol1}$, $N_{mol2}$ are the number of copies of molecule 1 and molecule 2. The contact maps aggregated by particle type (centre column) are a matrix reduction of the contact map by particle index. Bead abundance in a molecule is shown by a blue dashed line. The contact maps (right column) are normalised by the square root of the relative bead abundance in the molecules.

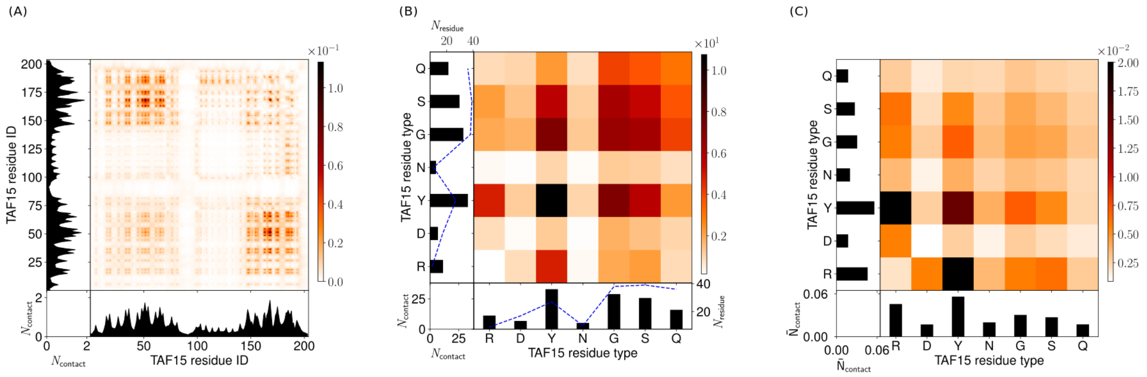

Figure S25: Contact maps for a droplet simulation with 120 TAF15 molecules (0%, 0%, 100%) at 150 mM and 300 K. (A) Intermolecular contact map by particle index for TAF15. (B) Intermolecular contact map by particle type for TAF15. (C) Intermolecular contact map by particle type, normalised by particle abundance for TAF15. The contacts in the contact maps by particle index (left column) are normalised by the number of frames used (600) and $(N_{mol1}N_{mol2})^{0.5}$ where $N_{mol1}$, $N_{mol2}$ are the number of copies of molecule 1 and molecule 2. The contact maps aggregated by particle type (centre column) are a matrix reduction of the contact map by particle index. Bead abundance in a molecule is shown by a blue dashed line. The contact maps (right column) are normalised by the square root of the relative bead abundance in the molecules.

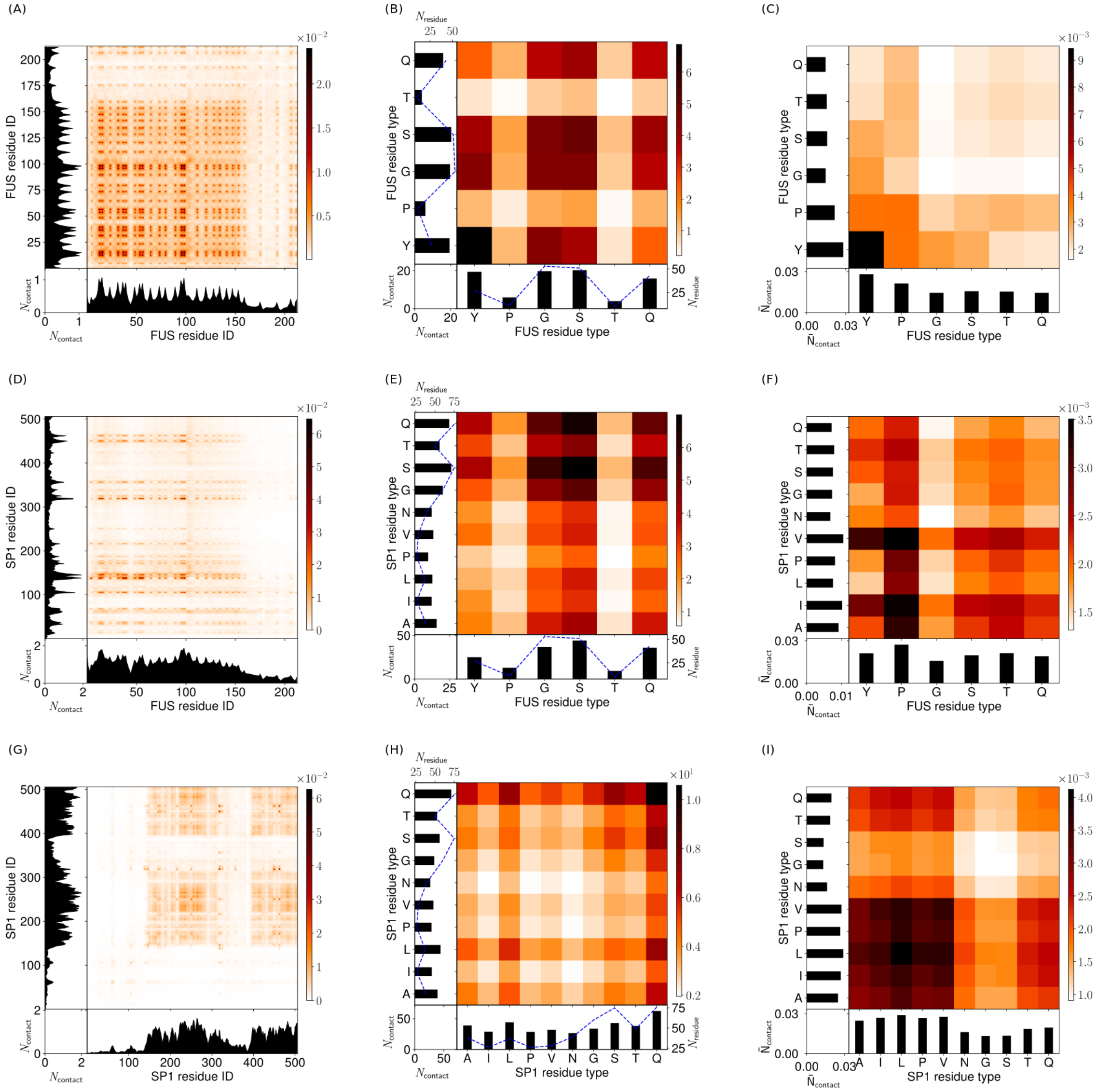

Figure S26: Contact maps for a droplet simulation with 60 FUS molecules and 30 SP1 molecules (50%, 50%, 0%) at 150 mM and 300 K. (A) Intermolecular contact map by particle index for FUS with other FUS. (B) Intermolecular contact map by particle type for FUS with other FUS. (C) Intermolecular contact map by particle type, normalised by particle abundance for FUS with other FUS. (D) Intermolecular contact map by particle index for FUS with SP1. (E) Intermolecular contact map by particle type for FUS with SP1. (F) Intermolecular contact map by particle type, normalised by particle abundance for FUS with SP1. (G) Intermolecular contact map by particle index for SP1 with other SP1. (H) Intermolecular contact map by particle type for SP1 with other SP1. (I) Intermolecular contact map by particle type, normalised by particle abundance for SP1 with other SP1. The contacts in the contact maps by particle index (left column) are normalised by the number of frames used (600) and $(N_{mol1}N_{mol2})^{0.5}$ where $N_{mol1}$, $N_{mol2}$ are the number of copies of molecule 1 and molecule 2. The contact maps aggregated by particle type (centre column) are a matrix reduction of the contact map by particle index. Bead abundance in a molecule is shown by a blue dashed line. The contact maps (right column) are normalised by the square root of the relative bead abundance in the molecules.

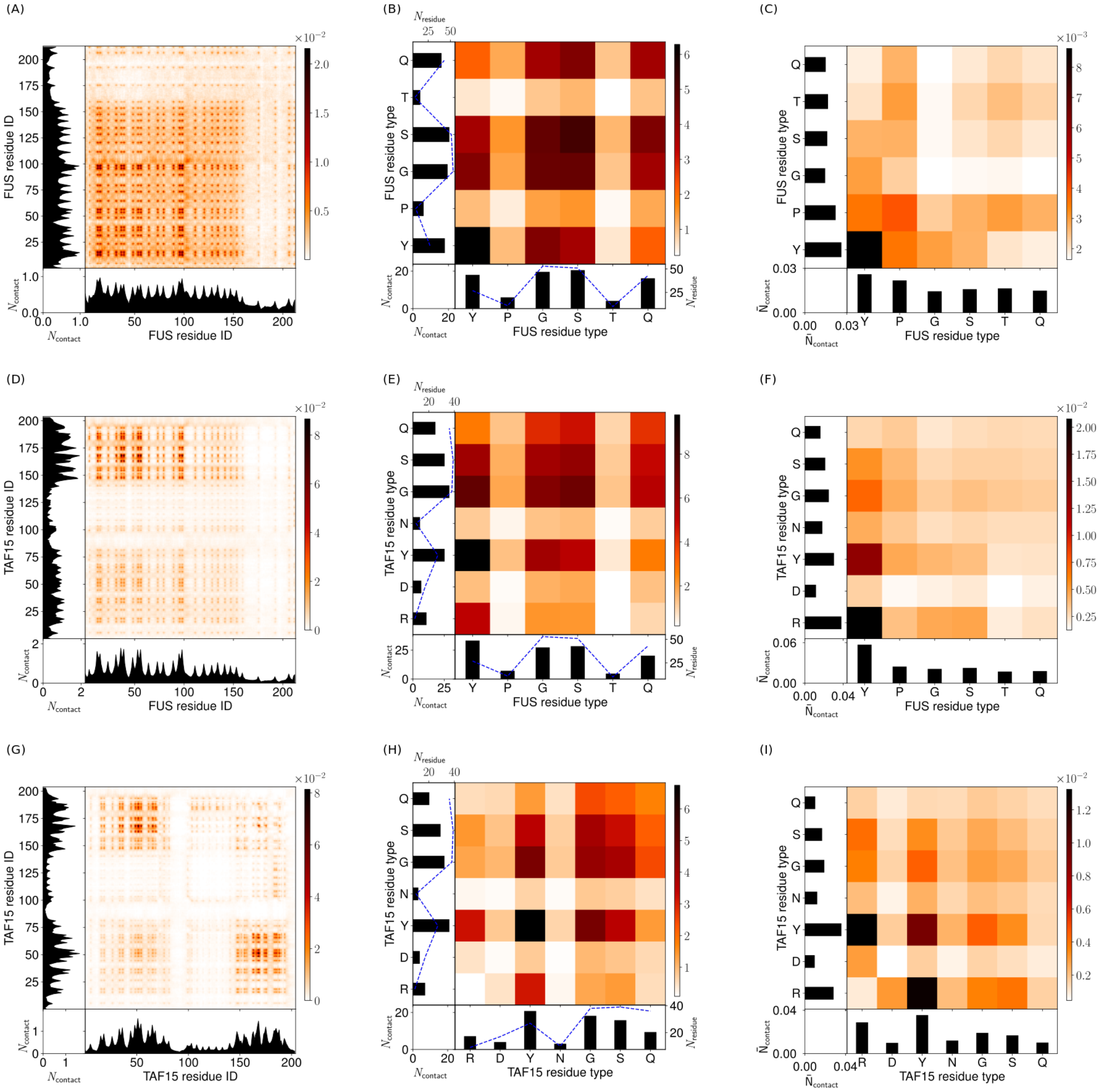

Figure S27: Contact maps for a droplet simulation with 60 FUS molecules and 60 TAF15 molecules (50%, 0%, 50%)at 150 mM and 300 K. (A) Intermolecular contact map by particle index for FUS with other FUS. (B) Intermolecular contact map by particle type for FUS with other FUS. (C) Intermolecular contact map by particle type, normalised by particle abundance for FUS with other FUS. (D) Intermolecular contact map by particle index for FUS with TAF15. (E) Intermolecular contact map by particle type for FUS with TAF15. (F) Intermolecular contact map by particle type, normalised by particle abundance for FUS with TAF15. (G) Intermolecular contact map by particle index for TAF15 with other TAF15. (H) Intermolecular contact map by particle type for TAF15 with other TAF15. (I) Intermolecular contact map by particle type, normalised by particle abundance for TAF15 with other TAF15. The contacts in the contact maps by particle index (left column) are normalised by the number of frames used (600) and $(N_{mol1}N_{mol2})^{0.5}$ where $N_{mol1}$, $N_{mol2}$ are the number of copies of molecule 1 and molecule 2. The contact maps aggregated by particle type (centre column) are a matrix reduction of the contact map by particle index. Bead abundance in a molecule is shown by a blue dashed line. The contact maps (right column) are normalised by the square root of the relative bead abundance in the molecules.

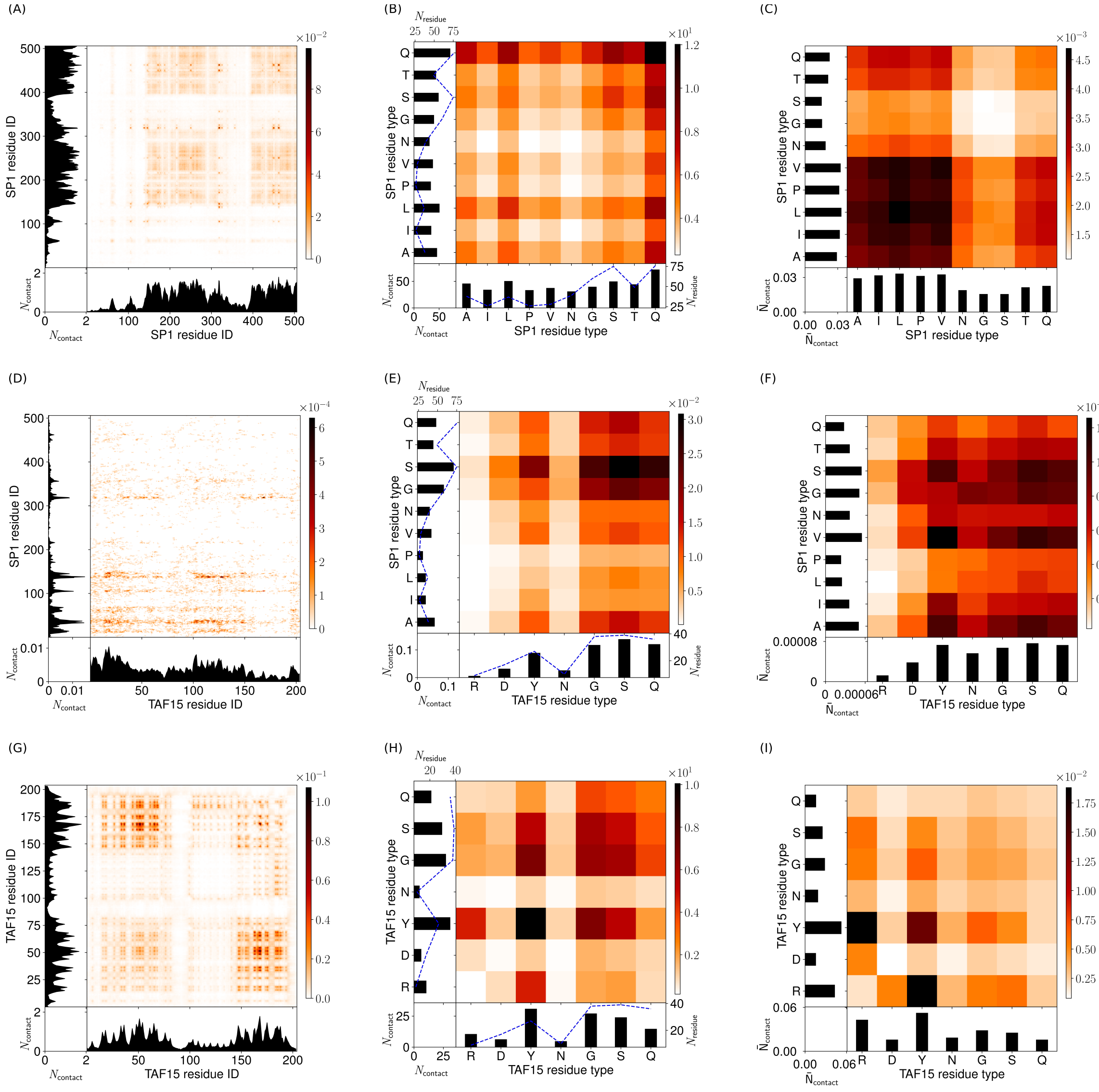

Figure S28: Contact maps for a droplet simulation with 30 SP1 molecules and 60 TAF15 molecules (0%, 50%, 50%) at 150 mM and 300 K. (A) Intermolecular contact map by particle index for SP1 with other SP1. (B) Intermolecular contact map by particle type for SP1 with other SP1. (C) Intermolecular contact map by particle type, normalised by particle abundance for SP1 with other SP1. (D) Intermolecular contact map by particle index for SP1 with TAF15. (E) Intermolecular contact map by particle type for SP1 with TAF15. (F) Intermolecular contact map by particle type, normalised by particle abundance for SP1 with TAF15. (G) Intermolecular contact map by particle index for TAF15 with other TAF15. (H) Intermolecular contact map by particle type for TAF15 with other TAF15. (I) Intermolecular contact map by particle type, normalised by particle abundance for TAF15 with other TAF15. The contacts in the contact maps by particle index (left column) are normalised by the number of frames used (600) and $(N_{mol1}N_{mol2})^{0.5}$ where $N_{mol1}$, $N_{mol2}$ are the number of copies of molecule 1 and molecule 2. The contact maps aggregated by particle type (centre column) are a matrix reduction of the contact map by particle index. Bead abundance in a molecule is shown by a blue dashed line. The contact maps (right column) are normalised by the square root of the relative bead abundance in the molecules.

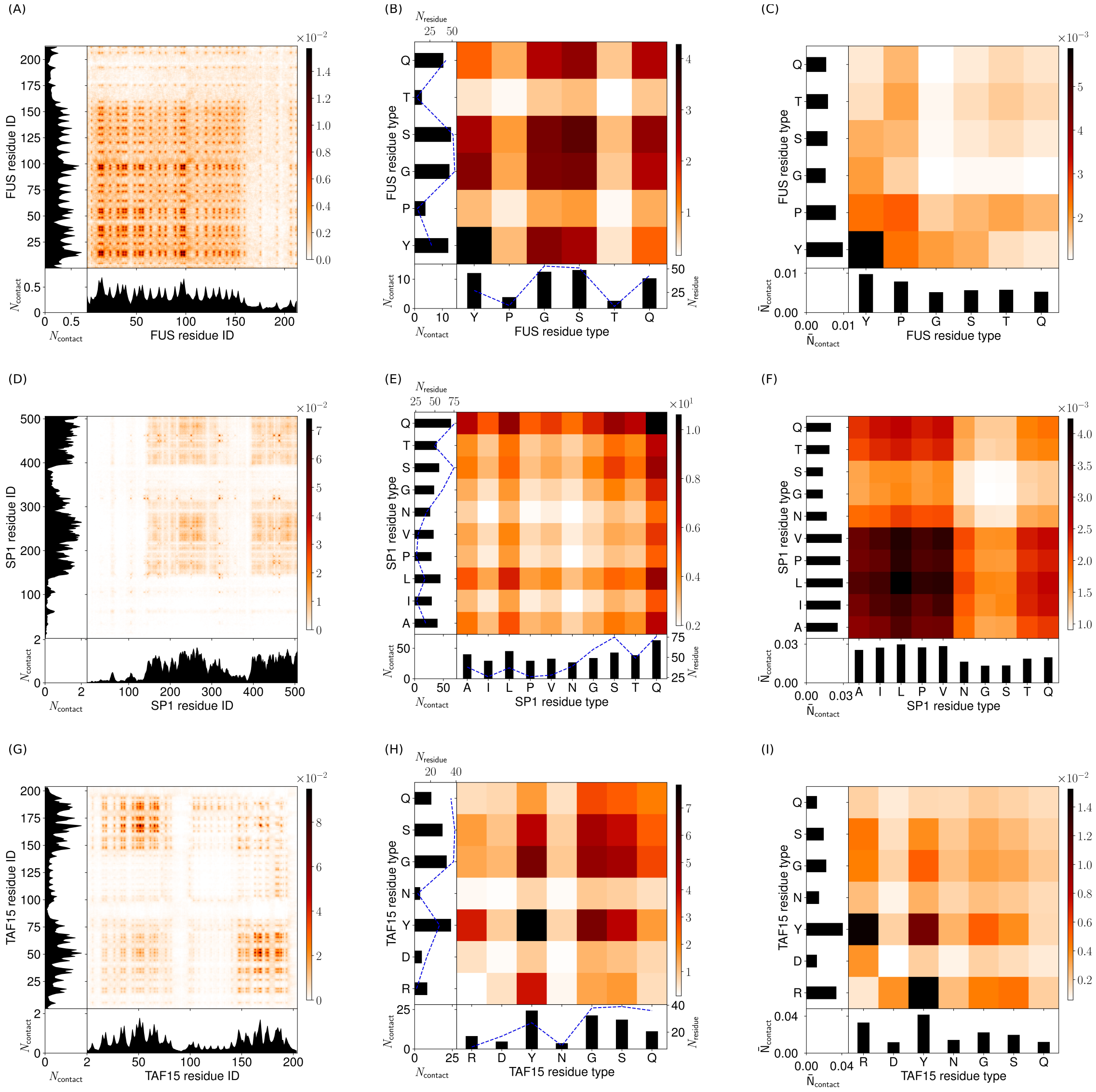

Figure S29: Contact maps for a droplet simulation with 40 FUS molecules, 20 SP1 molecules and 40 TAF15 molecules (33%, 33%, 33%) at 150 mM and 300 K. (A) Intermolecular contact map by particle index for FUS with other FUS. (B) Intermolecular contact map by particle type for FUS with other FUS. (C) Intermolecular contact map by particle type, normalised by particle abundance for FUS with other FUS. (D) Intermolecular contact map by particle index for SP1 with other SP1. (E) Intermolecular contact map by particle type for SP1 with other SP1. (F) Intermolecular contact map by particle type, normalised by particle abundance for SP1 with other SP1. (G) Intermolecular contact map by particle index for TAF15 with other TAF15. (H) Intermolecular contact map by particle type for TAF15 with other TAF15. (I) Intermolecular contact map by particle type, normalised by particle abundance for TAF15 with other TAF15. The contacts in the contact maps by particle index (left column) are normalised by the number of frames used (600) and $(N_{mol1}N_{mol2})^{0.5}$ where $N_{mol1}$, $N_{mol2}$ are the number of copies of molecule 1 and molecule 2. The contact maps aggregated by particle type (centre column) are a matrix reduction of the contact map by particle index. Bead abundance in a molecule is shown by a blue dashed line. The contact maps (right column) are normalised by the square root of the relative bead abundance in the molecules.

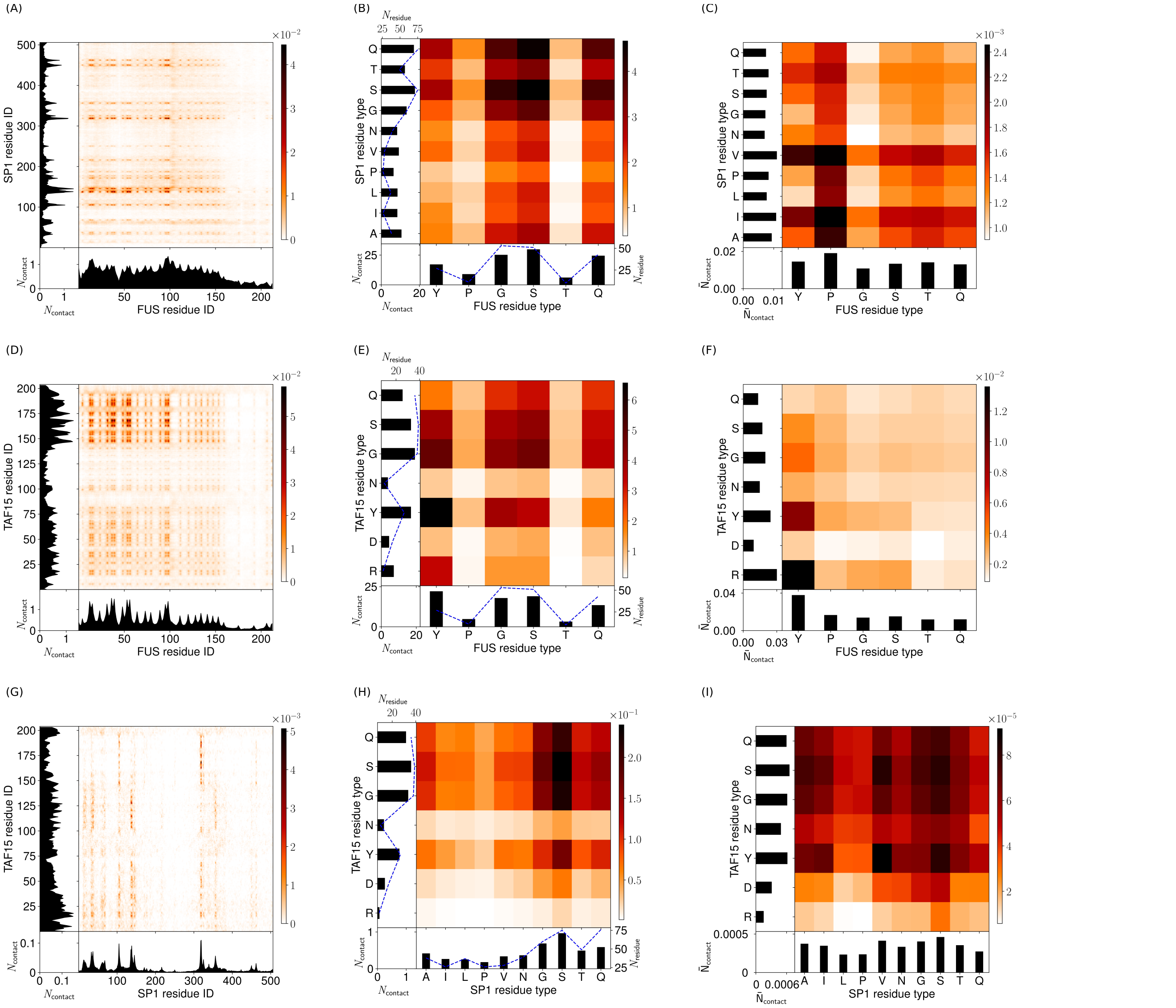

Figure S30: Contact maps for a droplet simulation with 40 FUS molecules, 20 SP1 molecules and 40 TAF15 molecules (33%, 33%, 33%) at 150 mM and 300 K. (A) Intermolecular contact map by particle index for FUS with SP1. (B) Intermolecular contact map by particle type for FUS with SP1. (C) Intermolecular contact map by particle type, normalised by particle abundance for FUS with SP1. (D) Intermolecular contact map by particle index for FUS with TAF15. (E) Intermolecular contact map by particle type for FUS with TAF15. (F) Intermolecular contact map by particle type, normalised by particle abundance for FUS with TAF15. (G) Intermolecular contact map by particle index for SP1 with TAF15. (H) Intermolecular contact map by particle type for SP1 with TAF15. (I) Intermolecular contact map by particle type, normalised by particle abundance for SP1 with TAF15. The contacts in the contact maps by particle index (left column) are normalised by the number of frames used (600) and $(N_{mol1}N_{mol2})^{0.5}$ where $N_{mol1}$, $N_{mol2}$ are the number of copies of molecule 1 and molecule 2. The contact maps aggregated by particle type (centre column) are a matrix reduction of the contact map by particle index. Bead abundance in a molecule is shown by a blue dashed line. The contact maps (right column) are normalised by the square root of the relative bead abundance in the molecules.

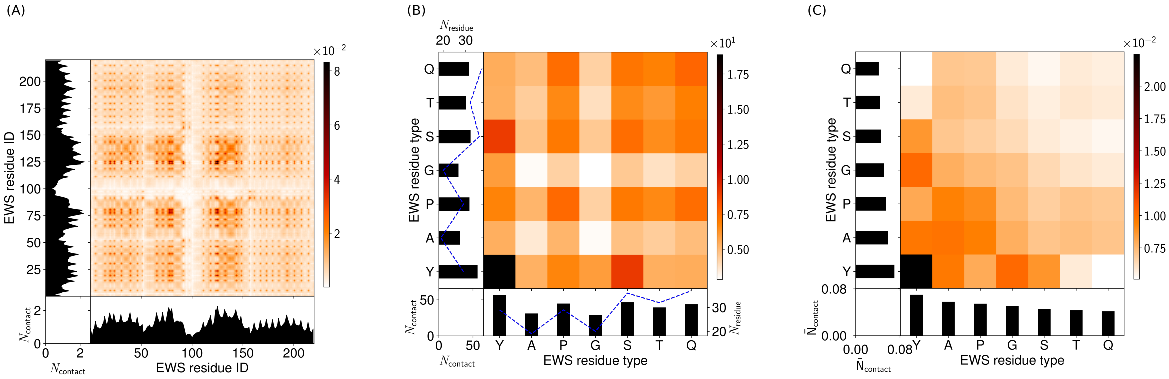

Figure S31: Molecule contact maps for a droplet simulation with 120 EWS molecules at 150 mM and 300 K. (A) Intermolecular contact map by particle index for EWS. (B) Intermolecular contact map by particle type for EWS. (C) Intermolecular contact map by particle type, normalised by particle abundance for EWS. The contacts in the contact maps by particle index (left column) are normalised by the number of frames used (600) and $(N_{mol1}N_{mol2})^{0.5}$ where $N_{mol1}$, $N_{mol2}$ are the number of copies of molecule 1 and molecule 2 in the simulation. The contact maps aggregated by particle type (centre column) are a matrix reduction of the contact map by particle index. Bead abundance in a molecule is shown by a blue dashed line. The contact maps (right column) are normalised by the square root of the relative bead abundance in the molecules.

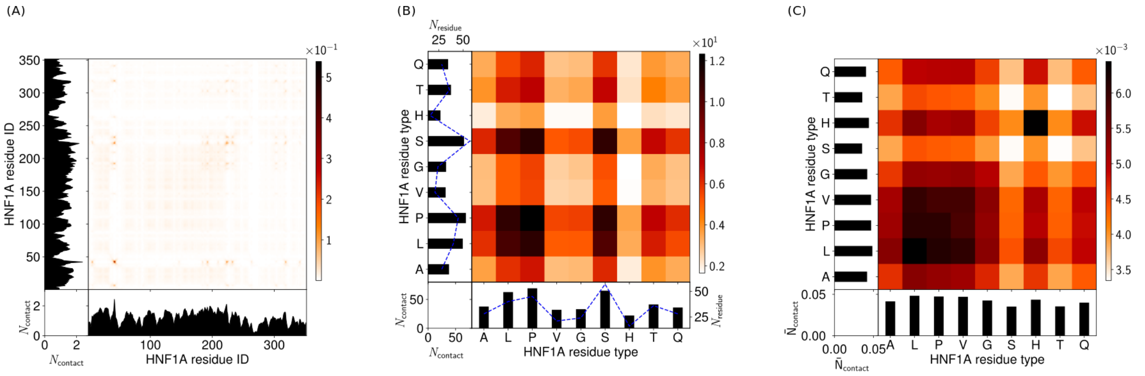

Figure S32: Molecule contact maps for a droplet simulation with 90 HNF1A molecules at 150 mM and 300 K. (A) Intermolecular contact map by particle index for HNF1A. (B) Intermolecular contact map by particle type for HNF1A. (C) Intermolecular contact map by particle type, normalised by particle abundance for HNF1A. The contacts in the contact maps by particle index (left column) are normalised by the number of frames used (600) and $(N_{mol1}N_{mol2})^{0.5}$ where $N_{mol1}$, $N_{mol2}$ are the number of copies of molecule 1 and molecule 2 in the simulation. The contact maps aggregated by particle type (centre column) are a matrix reduction of the contact map by particle index. Bead abundance in a molecule is shown by a blue dashed line. The contact maps (right column) are normalised by the square root of the relative bead abundance in the molecules.

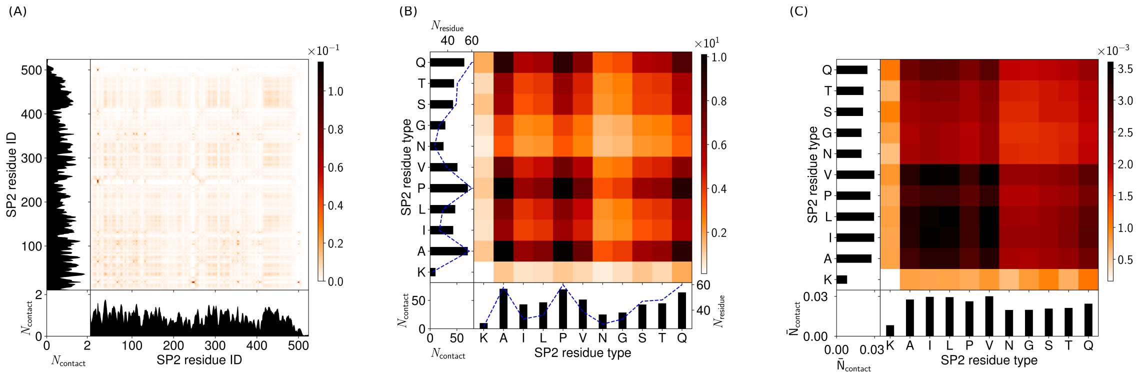

Figure S33: Molecule contact maps for a droplet simulation with 60 SP2 molecules at 150 mM and 300 K. (A) Intermolecular contact map by particle index for SP2. (B) Intermolecular contact map by particle type for SP2. (C) Intermolecular contact map by particle type, normalised by particle abundance for SP2. The contacts in the contact maps by particle index (left column) are normalised by the number of frames used (600) and $(N_{mol1}N_{mol2})^{0.5}$ where $N_{mol1}$, $N_{mol2}$ are the number of copies of molecule 1 and molecule 2 in the simulation. The contact maps aggregated by particle type (centre column) are a matrix reduction of the contact map by particle index. Bead abundance in a molecule is shown by a blue dashed line. The contact maps (right column) are normalised by the square root of the relative bead abundance in the molecules.

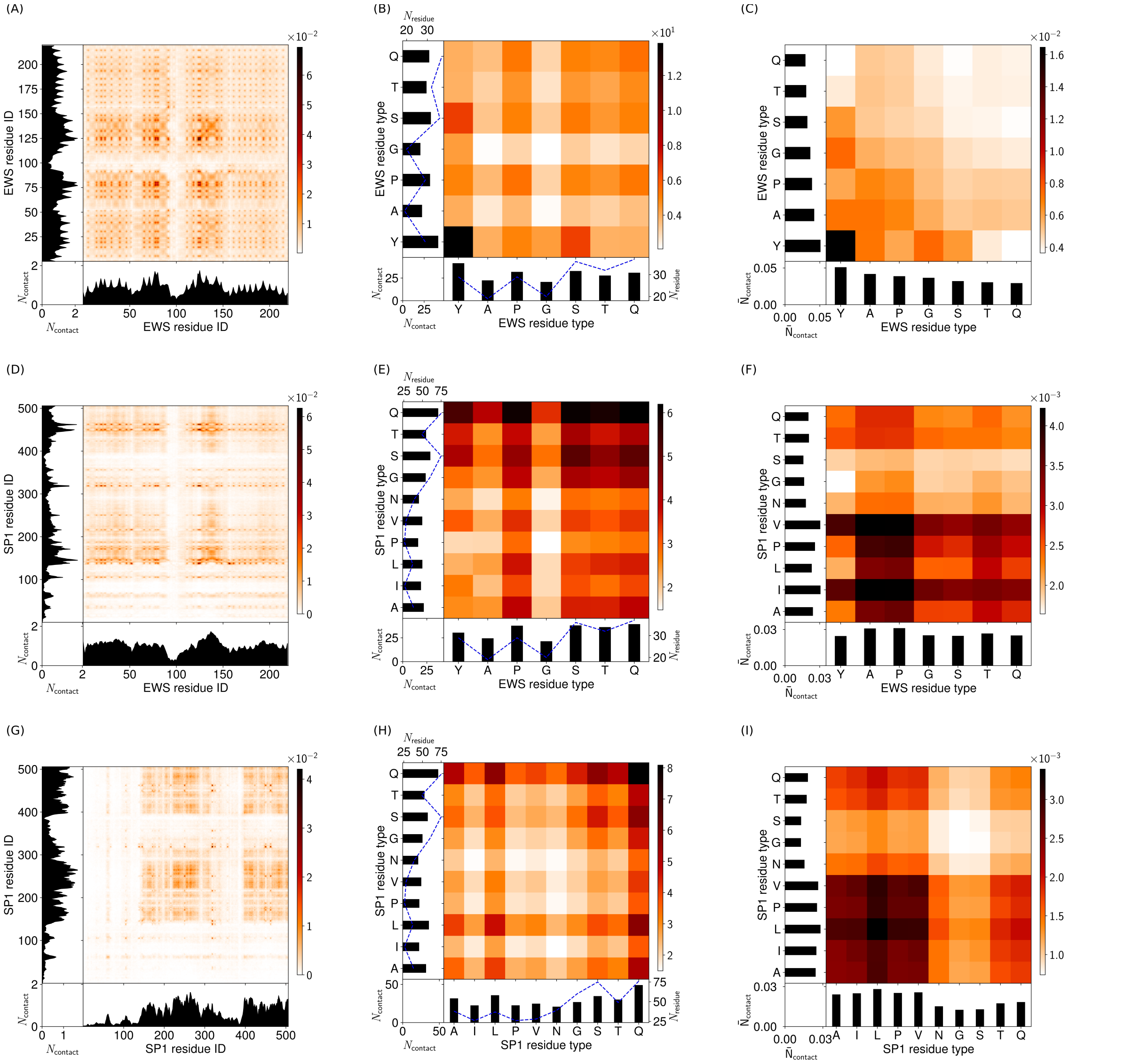

Figure S34: Molecule contact maps for a droplet simulation with 60 EWS molecules and 30 SP1 molecules at 150 mM and 300 K. (A) Intermolecular contact map by particle index for EWS with other EWS. (B) Intermolecular contact map by particle type for EWS with other EWS. (C) Intermolecular contact map by particle type, normalised by particle abundance for EWS with other EWS. (D) Intermolecular contact map by particle index for EWS with SP1. (E) Intermolecular contact map by particle type for EWS with SP1. (F) Intermolecular contact map by particle type, normalised by particle abundance for EWS with SP1. (G) Intermolecular contact map by particle index for SP1 with other SP1. (H) Intermolecular contact map by particle type for SP1 with other SP1. (I) Intermolecular contact map by particle type, normalised by particle abundance for SP1 with other SP1. The contacts in the contact maps by particle index (left column) are normalised by the number of frames used (600) and $(N_{mol1}N_{mol2})^{0.5}$ where $N_{mol1}$, $N_{mol2}$ are the number of copies of molecule 1 and molecule 2 in the simulation. The contact maps aggregated by particle type (centre column) are a matrix reduction of the contact map by particle index. Bead abundance in a molecule is shown by a blue dashed line. The contact maps (right column) are normalised by the square root of the relative bead abundance in the molecules.

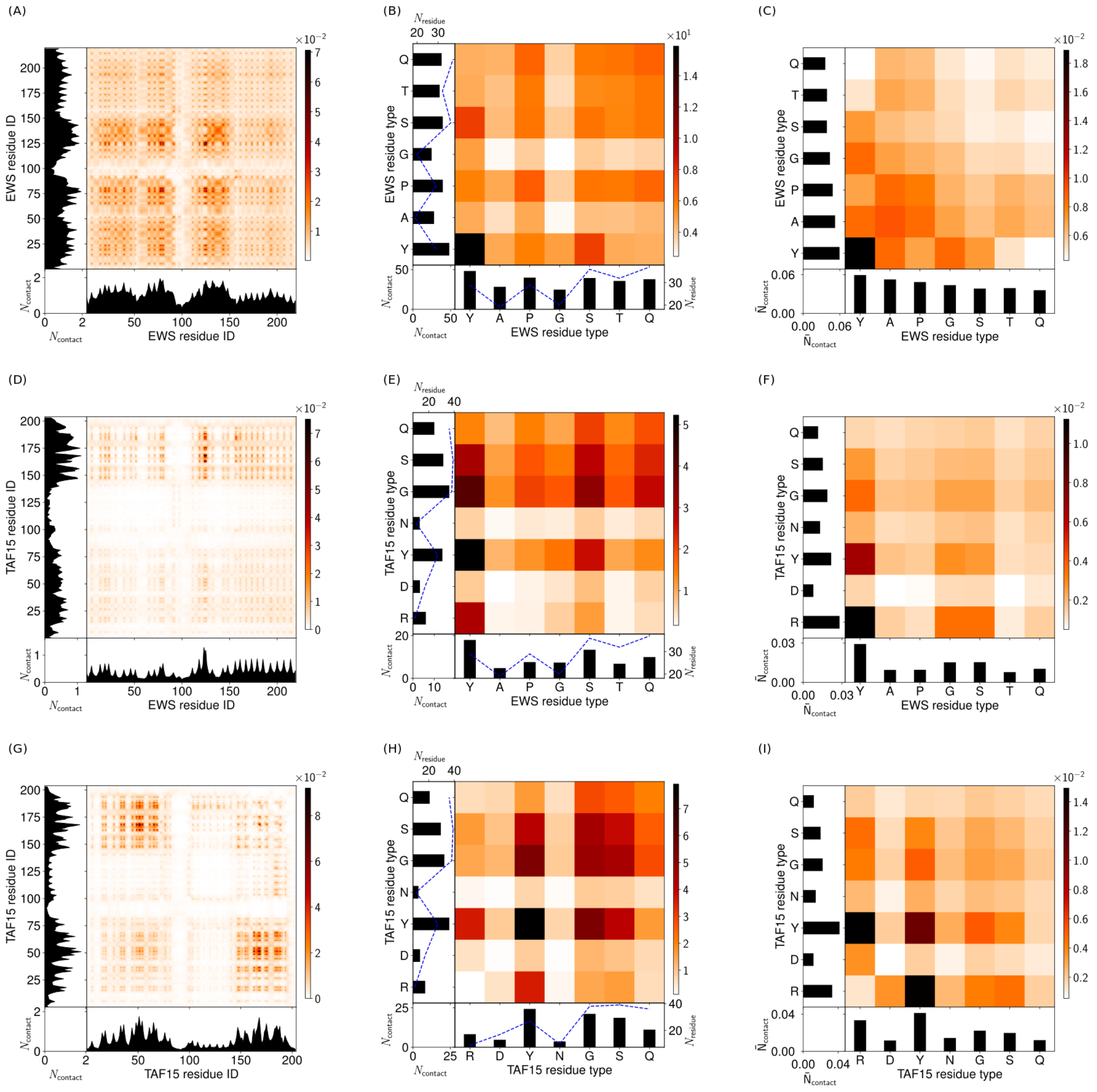

Figure S35: Molecule contact maps for a droplet simulation with 60 EWS molecules and 60 TAF15 molecules at 150 mM and 300 K. (A) Intermolecular contact map by particle index for EWS with other EWS. (B) Intermolecular contact map by particle type for EWS with other EWS. (C) Intermolecular contact map by particle type, normalised by particle abundance for EWS with other EWS. (D) Intermolecular contact map by particle index for EWS with TAF15. (E) Intermolecular contact map by particle type for EWS with TAF15. (F) Intermolecular contact map by particle type, normalised by particle abundance for EWS with TAF15. (G) Intermolecular contact map by particle index for TAF15 with other TAF15. (H) Intermolecular contact map by particle type for TAF15 with other TAF15. (I) Intermolecular contact map by particle type, normalised by particle abundance for TAF15 with other TAF15. The contacts in the contact maps by particle index (left column) are normalised by the number of frames used (600) and $(N_{mol1}N_{mol2})^{0.5}$ where $N_{mol1}$, $N_{mol2}$ are the number of copies of molecule 1 and molecule 2 in the simulation. The contact maps aggregated by particle type (centre column) are a matrix reduction of the contact map by particle index. Bead abundance in a molecule is shown by a blue dashed line. The contact maps (right column) are normalised by the square root of the relative bead abundance in the molecules.

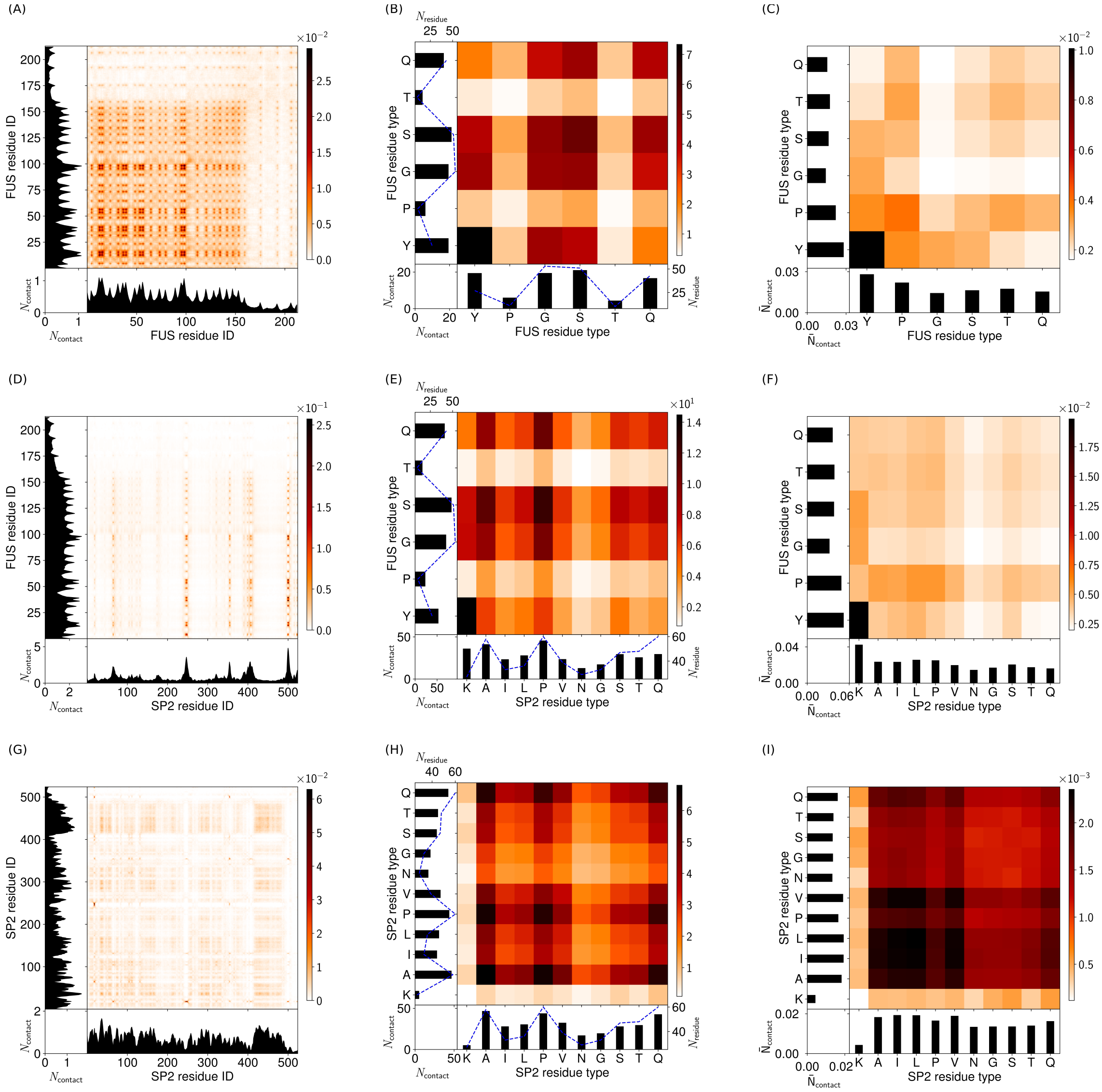

Figure S36: Molecule contact maps for a droplet simulation with 60 FUS molecules and 30 SP2 molecules at 150 mM and 300 K. (A) Intermolecular contact map by particle index for FUS with other FUS. (B) Intermolecular contact map by particle type for FUS with other FUS. (C) Intermolecular contact map by particle type, normalised by particle abundance for FUS with other FUS. (D) Intermolecular contact map by particle index for FUS with SP11. (E) Intermolecular contact map by particle type for FUS with SP2. (F) Intermolecular contact map by particle type, normalised by particle abundance for FUS with SP2. (G) Intermolecular contact map by particle index for SP2 with other SP2. (H) Intermolecular contact map by particle type for SP2 with other SP2. (I) Intermolecular contact map by particle type, normalised by particle abundance for SP2 with other SP2. The contacts in the contact maps by particle index (left column) are normalised by the number of frames used (600) and $(N_{mol1}N_{mol2})^{0.5}$ where $N_{mol1}$, $N_{mol2}$ are the number of copies of molecule 1 and molecule 2 in the simulation. The contact maps aggregated by particle type (centre column) are a matrix reduction of the contact map by particle index. Bead abundance in a molecule is shown by a blue dashed line. The contact maps (right column) are normalised by the square root of the relative bead abundance in the molecules.

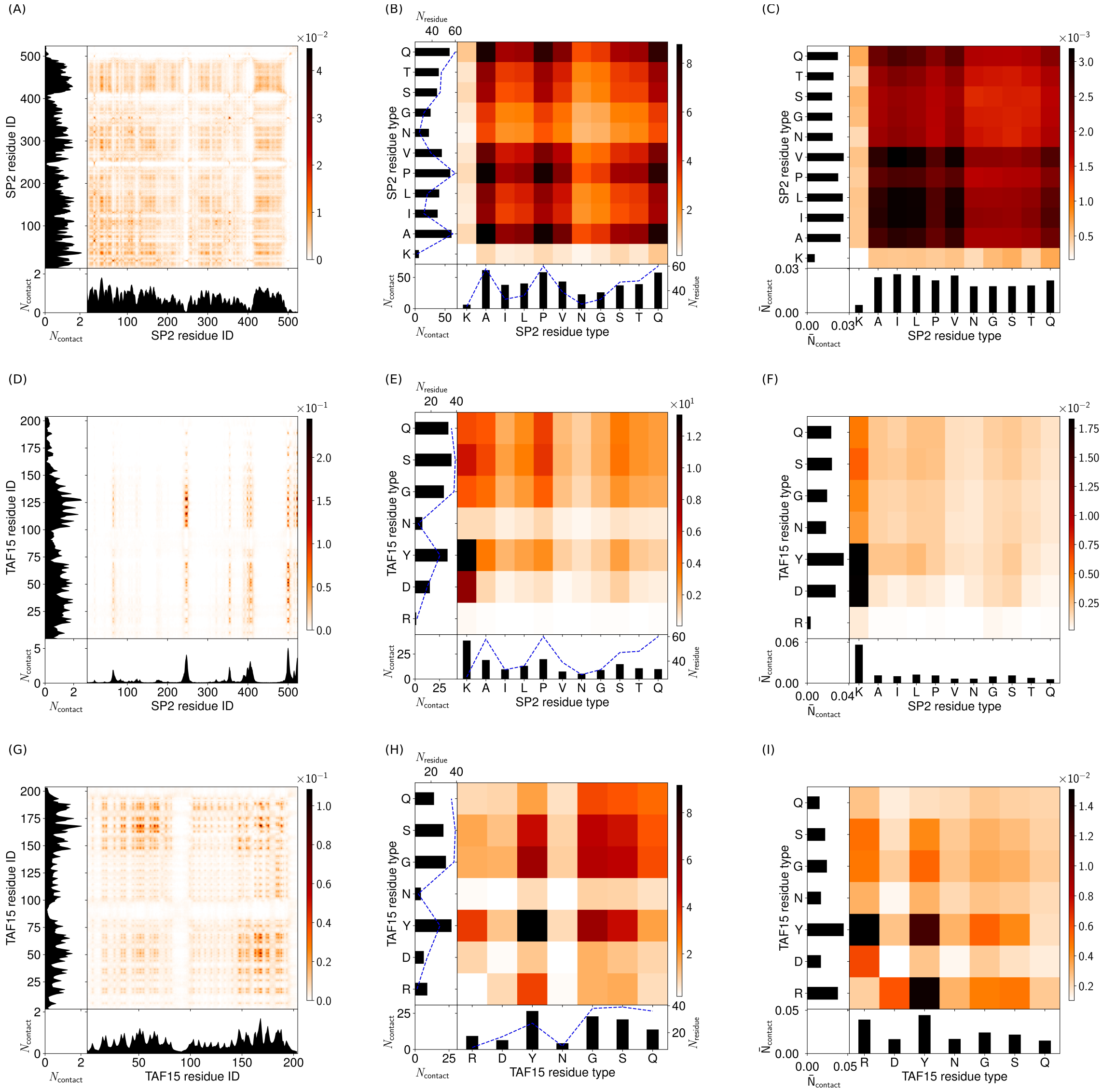

Figure S37: Molecule contact maps for a droplet simulation with 30 SP2 molecules and 60 TAF15 molecules at 150 mM and 300 K. (A) Intermolecular contact map by particle index for SP2 with other SP2. (B) Intermolecular contact map by particle type for SP2 with other SP2. (C) Intermolecular contact map by particle type, normalised by particle abundance for SP2 with other SP2. (D) Intermolecular contact map by particle index for SP2 with TAF15. (E) Intermolecular contact map by particle type for SP2 with TAF15. (F) Intermolecular contact map by particle type, normalised by particle abundance for SP2 with TAF15. (G) Intermolecular contact map by particle index for TAF15 with other TAF15. (H) Intermolecular contact map by particle type for TAF15 with other TAF15. (I) Intermolecular contact map by particle type, normalised by particle abundance for TAF15 with other TAF15. The contacts in the contact maps by particle index (left column) are normalised by the number of frames used (600) and $(N_{mol1}N_{mol2})^{0.5}$ where $N_{mol1}$, $N_{mol2}$ are the number of copies of molecule 1 and molecule 2 in the simulation. The contact maps aggregated by particle type (centre column) are a matrix reduction of the contact map by particle index. Bead abundance in a molecule is shown by a blue dashed line. The contact maps (right column) are normalised by the square root of the relative bead abundance in the molecules.

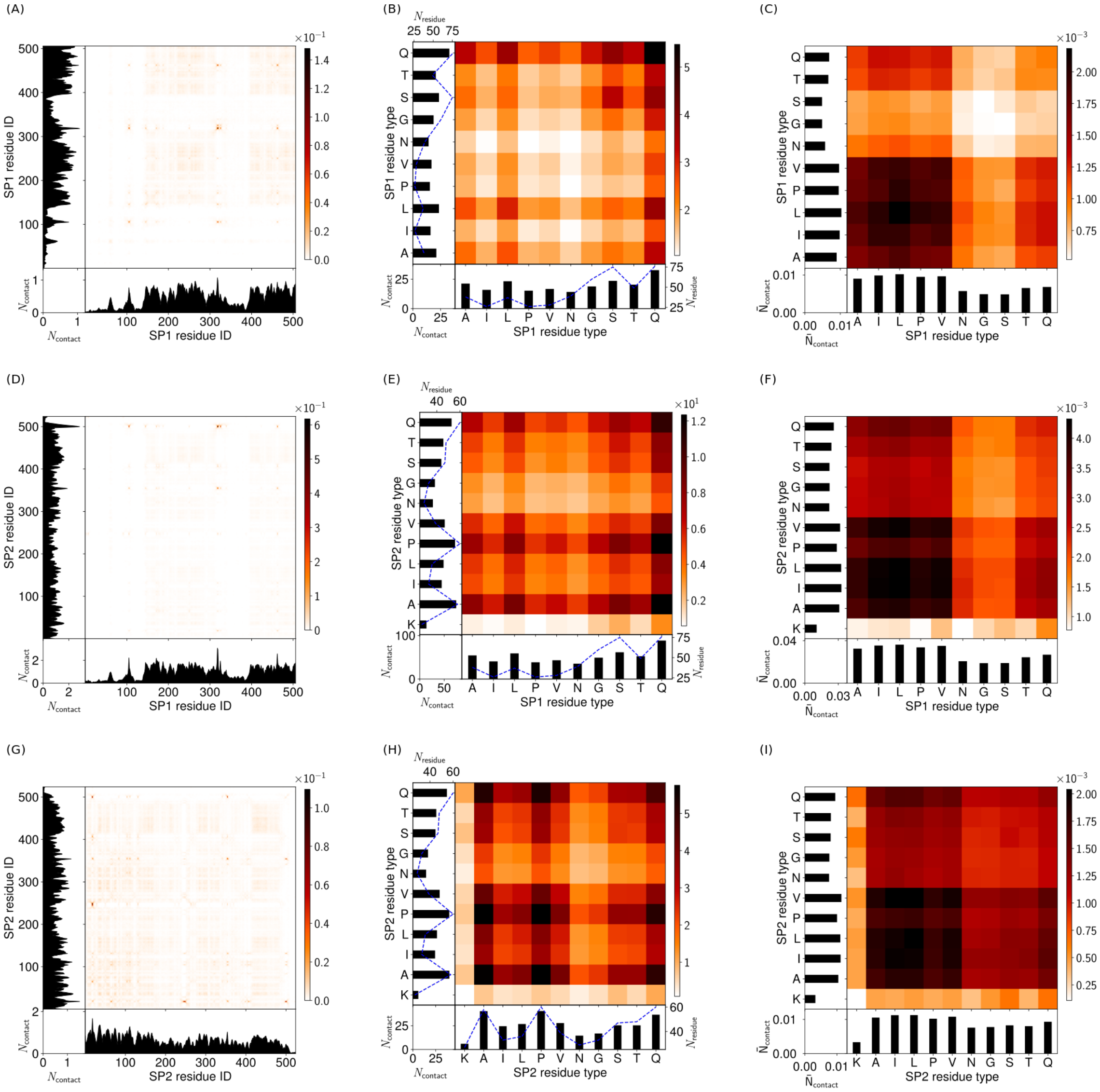

Figure S38: Molecule contact maps for a droplet simulation with 30 SP1 molecules and 30 SP2 molecules at 150 mM and 300 K. (A) Intermolecular contact map by particle index for SP1 with other SP1. (B) Intermolecular contact map by particle type for SP1 with other SP1. (C) Intermolecular contact map by particle type, normalised by particle abundance for SP1 with other SP1. (D) Intermolecular contact map by particle index for SP1 with SP2. (E) Intermolecular contact map by particle type for SP1 with SP2. (F) Intermolecular contact map by particle type, normalised by particle abundance for SP1 with SP2. (G) Intermolecular contact map by particle index for SP2 with other SP2. (H) Intermolecular contact map by particle type for SP2 with other SP2. (I) Intermolecular contact map by particle type, normalised by particle abundance for SP2 with other SP2. The contacts in the contact maps by particle index (left column) are normalised by the number of frames used (600) and $(N_{mol1}N_{mol2})^{0.5}$ where $N_{mol1}$, $N_{mol2}$ are the number of copies of molecule 1 and molecule 2 in the simulation. The contact maps aggregated by particle type (centre column) are a matrix reduction of the contact map by particle index. Bead abundance in a molecule is shown by a blue dashed line. The contact maps (right column) are normalised by the square root of the relative bead abundance in the molecules.

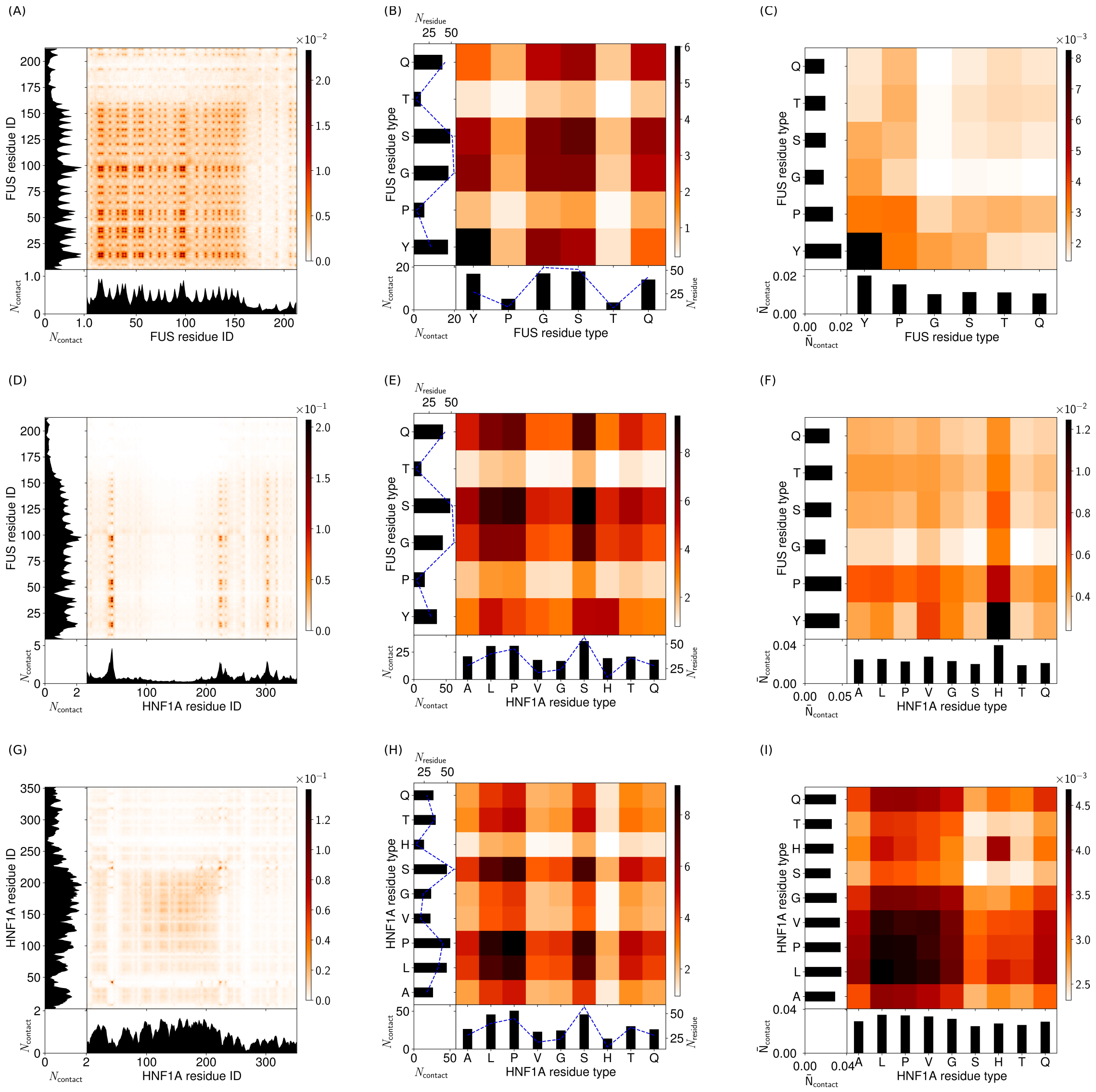

Figure S39: Molecule contact maps for a droplet simulation with 60 FUS molecules and 45 HNF1A molecules at 150 mM and 300 K. (A) Intermolecular contact map by particle index for FUS with other FUS. (B) Intermolecular contact map by particle type for FUS with other FUS. (C) Intermolecular contact map by particle type, normalised by particle abundance for FUS with other FUS. (D) Intermolecular contact map by particle index for FUS with HNF1A. (E) Intermolecular contact map by particle type for FUS with HNF1A. (F) Intermolecular contact map by particle type, normalised by particle abundance for FUS with HNF1A. (G) Intermolecular contact map by particle index for HNF1A with other HNF1A. (H) Intermolecular contact map by particle type for HNF1A with other HNF1A. (I) Intermolecular contact map by particle type, normalised by particle abundance for HNF1A with other HNF1A. The contacts in the contact maps by particle index (left column) are normalised by the number of frames used (600) and $(N_{mol1}N_{mol2})^{0.5}$ where $N_{mol1}$, $N_{mol2}$ are the number of copies of molecule 1 and molecule 2 in the simulation. The contact maps aggregated by particle type (centre column) are a matrix reduction of the contact map by particle index. Bead abundance in a molecule is shown by a blue dashed line. The contact maps (right column) are normalised by the square root of the relative bead abundance in the molecules.

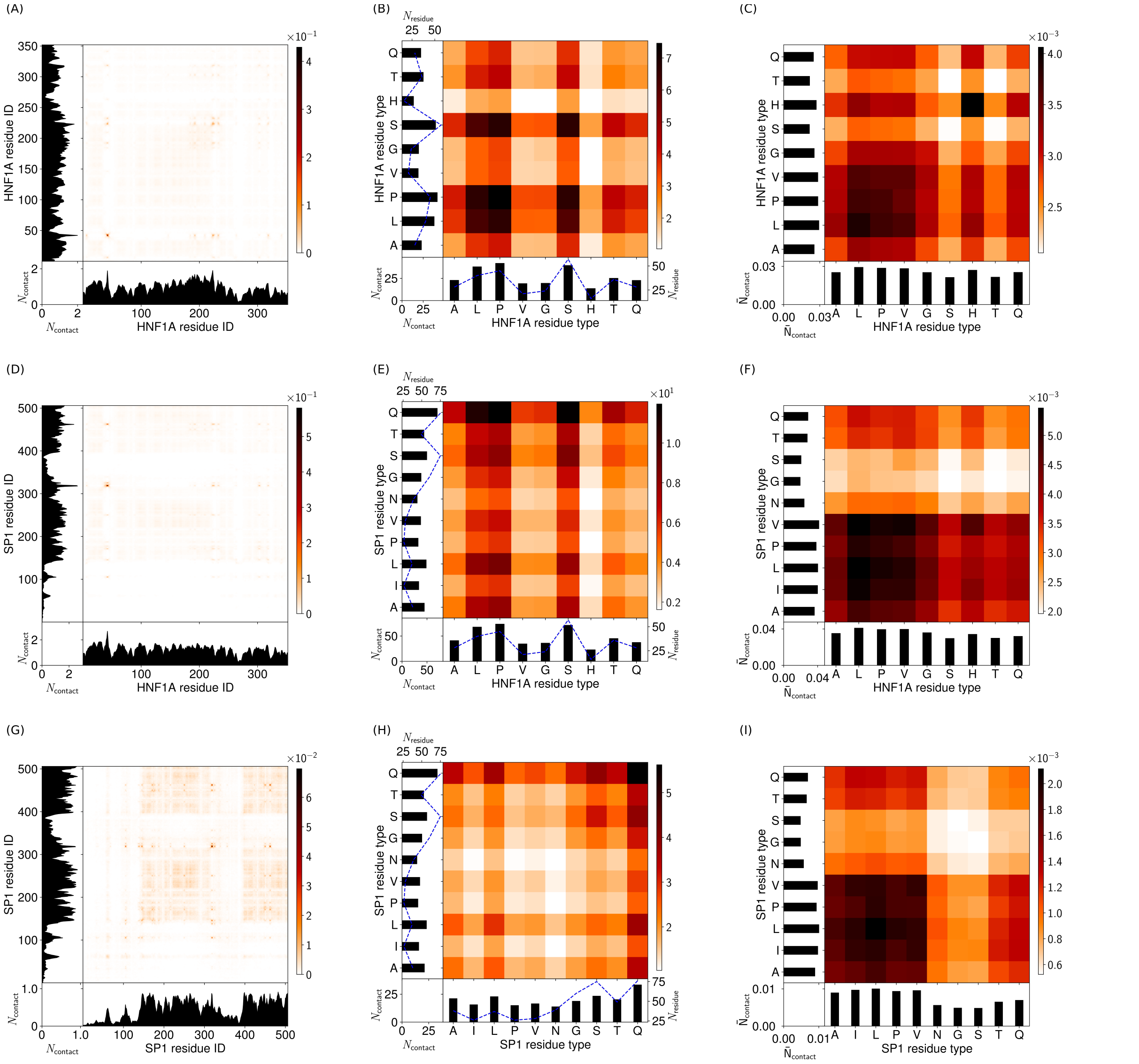

Figure S40: Molecule contact maps for a droplet simulation with 45 HNF1A molecules and 30 SP1 molecules at 150 mM and 300 K. (A) Intermolecular contact map by particle index for HNF1A with other HNF1A. (B) Intermolecular contact map by particle type for HNF1A with other HNF1A. (C) Intermolecular contact map by particle type, normalised by particle abundance for HNF1A with other HNF1A. (D) Intermolecular contact map by particle index for HNF1A with SP1. (E) Intermolecular contact map by particle type for HNF1A with SP1. (F) Intermolecular contact map by particle type, normalised by particle abundance for HNF1A with SP1. (G) Intermolecular contact map by particle index for SP1 with other SP1. (H) Intermolecular contact map by particle type for SP1 with other SP1. (I) Intermolecular contact map by particle type, normalised by particle abundance for SP1 with other SP1. The contacts in the contact maps by particle index (left column) are normalised by the number of frames used (600) and $(N_{mol1}N_{mol2})^{0.5}$ where $N_{mol1}$, $N_{mol2}$ are the number of copies of molecule 1 and molecule 2 in the simulation. The contact maps aggregated by particle type (centre column) are a matrix reduction of the contact map by particle index. Bead abundance in a molecule is shown by a blue dashed line. The contact maps (right column) are normalised by the square root of the relative bead abundance in the molecules.

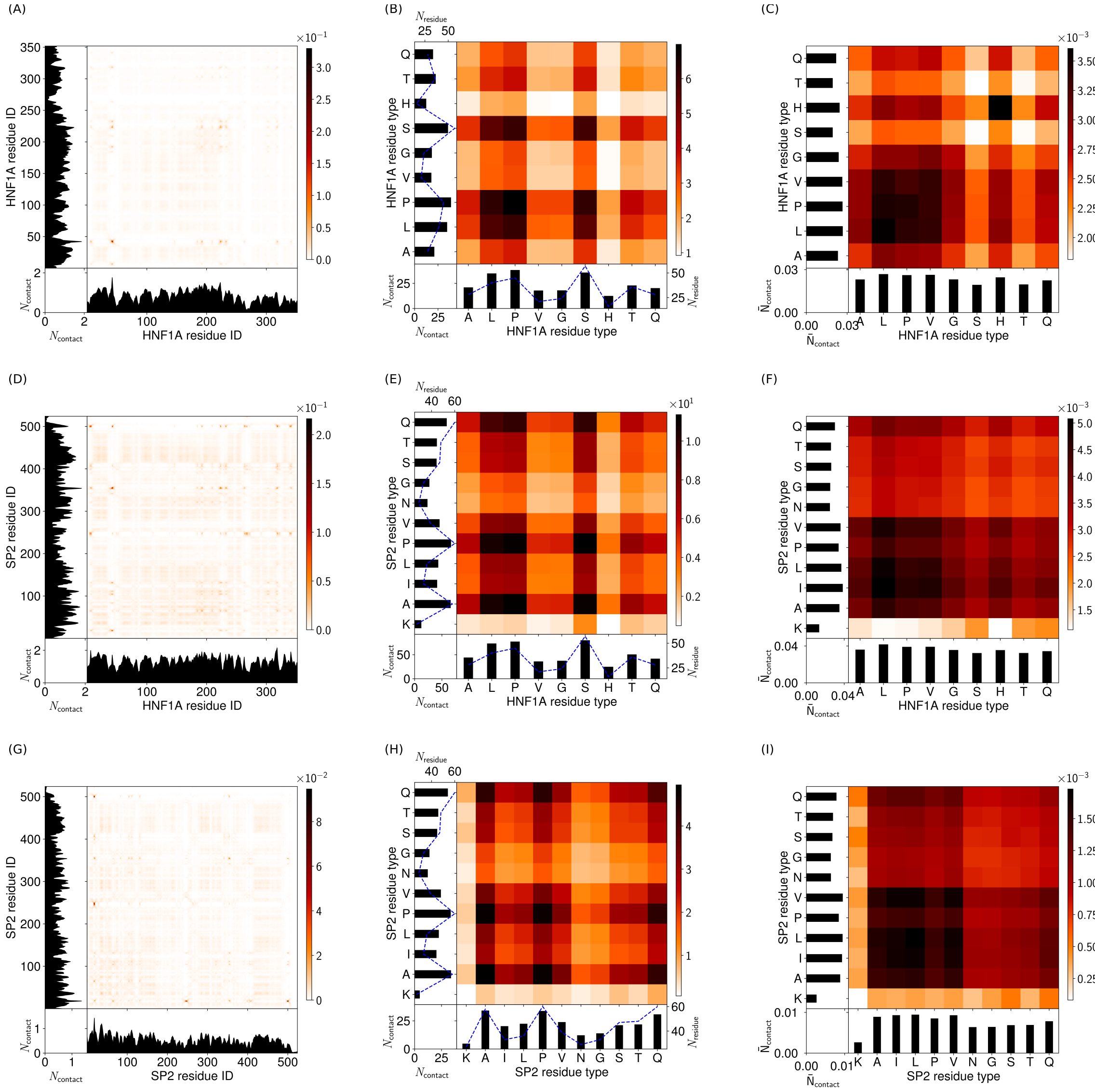

Figure S41: Molecule contact maps for a droplet simulation with 45 HNF1A molecules and 30 SP2 molecules at 150 mM and 300 K. (A) Intermolecular contact map by particle index for HNF1A with other HNF1A. (B) Intermolecular contact map by particle type for HNF1A with other HNF1A. (C) Intermolecular contact map by particle type, normalised by particle abundance for HNF1A with other HNF1A. (D) Intermolecular contact map by particle index for HNF1A with SP2. (E) Intermolecular contact map by particle type for HNF1A with SP2. (F) Intermolecular contact map by particle type, normalised by particle abundance for HNF1A with SP2. (G) Intermolecular contact map by particle index for SP2 with other SP2. (H) Intermolecular contact map by particle type for SP2 with other SP2. (I) Intermolecular contact map by particle type, normalised by particle abundance for SP2 with other SP2. The contacts in the contact maps by particle index (left column) are normalised by the number of frames used (600) and $(N_{mol1}N_{mol2})^{0.5}$ where $N_{mol1}$, $N_{mol2}$ are the number of copies of molecule 1 and molecule 2 in the simulation. The contact maps aggregated by particle type (centre column) are a matrix reduction of the contact map by particle index. Bead abundance in a molecule is shown by a blue dashed line. The contact maps (right column) are normalised by the square root of the relative bead abundance in the molecules.

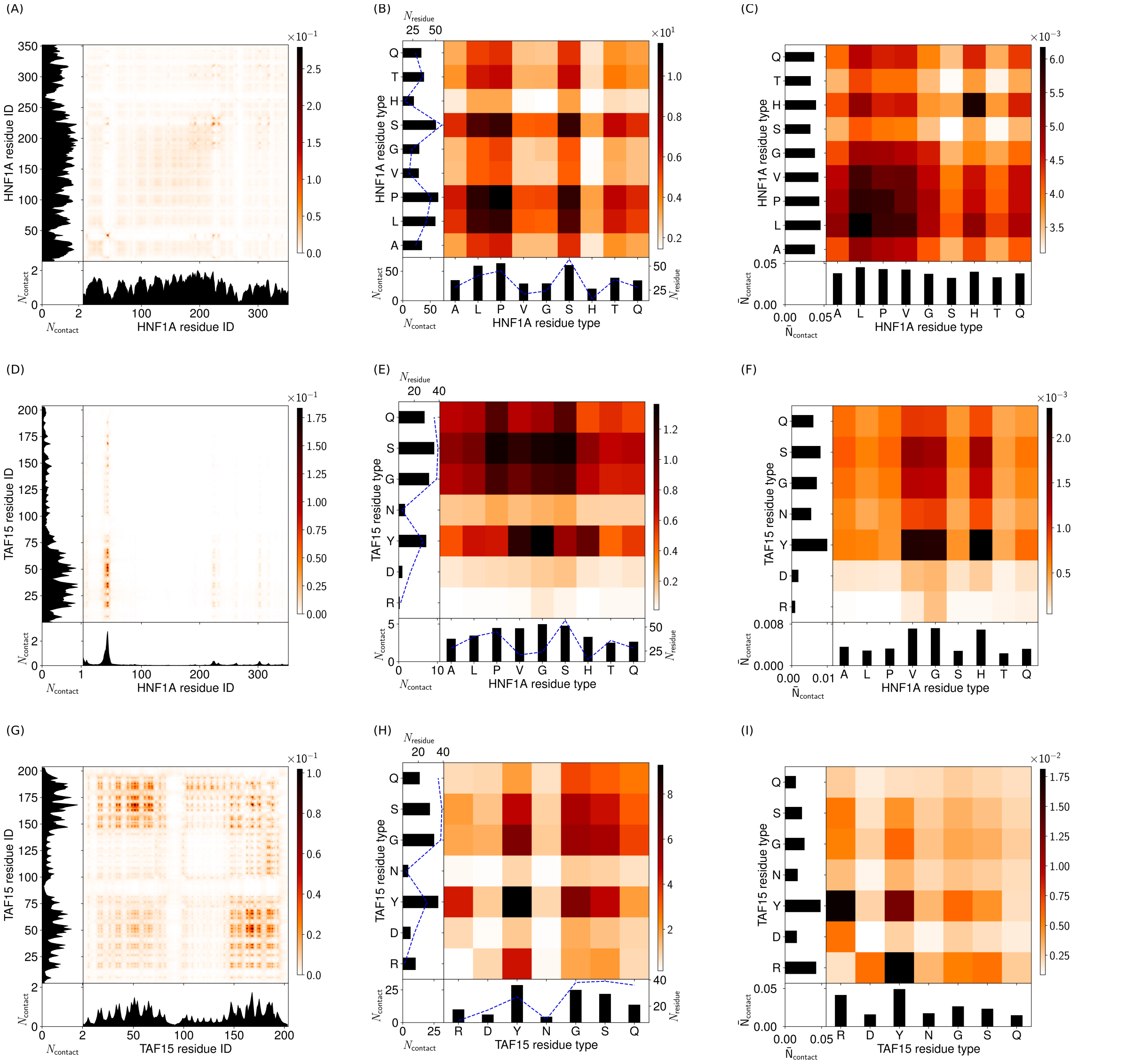

Figure S42: Molecule contact maps for a droplet simulation with 45 HNF1A molecules and 60 TAF15 molecules at 150 mM and 300 K. (A) Intermolecular contact map by particle index for HNF1A with other HNF1A. (B) Intermolecular contact map by particle type for HNF1A with other HNF1A. (C) Intermolecular contact map by particle type, normalised by particle abundance for HNF1A with other HNF1A. (D) Intermolecular contact map by particle index for HNF1A with TAF15. (E) Intermolecular contact map by particle type for HNF1A with TAF15. (F) Intermolecular contact map by particle type, normalised by particle abundance for HNF1A with TAF15. (G) Intermolecular contact map by particle index for TAF15 with other TAF15. (H) Intermolecular contact map by particle type for TAF15 with other TAF15. (I) Intermolecular contact map by particle type, normalised by particle abundance for TAF15 with other TAF15. The contacts in the contact maps by particle index (left column) are normalised by the number of frames used (600) and $(N_{mol1}N_{mol2})^{0.5}$ where $N_{mol1}$, $N_{mol2}$ are the number of copies of molecule 1 and molecule 2 in the simulation. The contact maps aggregated by particle type (centre column) are a matrix reduction of the contact map by particle index. Bead abundance in a molecule is shown by a blue dashed line. The contact maps (right column) are normalised by the square root of the relative bead abundance in the molecules.

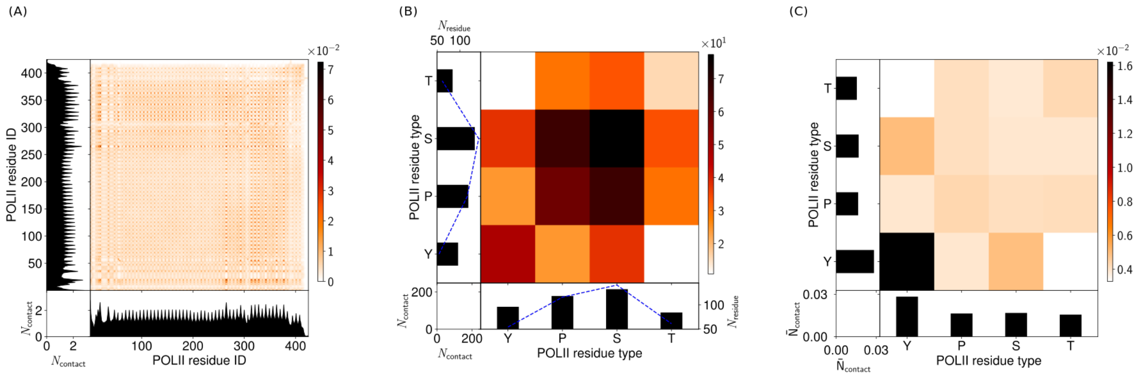

Figure S43: Molecule contact maps for a droplet simulation with 60 POLII molecules at 150 mM and 300 K. (A) Intermolecular contact map by particle index for POLII. (B) Intermolecular contact map by particle type for POLII. (C) Intermolecular contact map by particle type, normalised by particle abundance for POLII. The contacts in the contact maps by particle index (left column) are normalised by the number of frames used (600) and $(N_{mol1}N_{mol2})^{0.5}$ where $N_{mol1}$, $N_{mol2}$ are the number of copies of molecule 1 and molecule 2 in the simulation. The contact maps aggregated by particle type (centre column) are a matrix reduction of the contact map by particle index. Bead abundance in a molecule is shown by a blue dashed line. The contact maps (right column) are normalised by the square root of the relative bead abundance in the molecules.

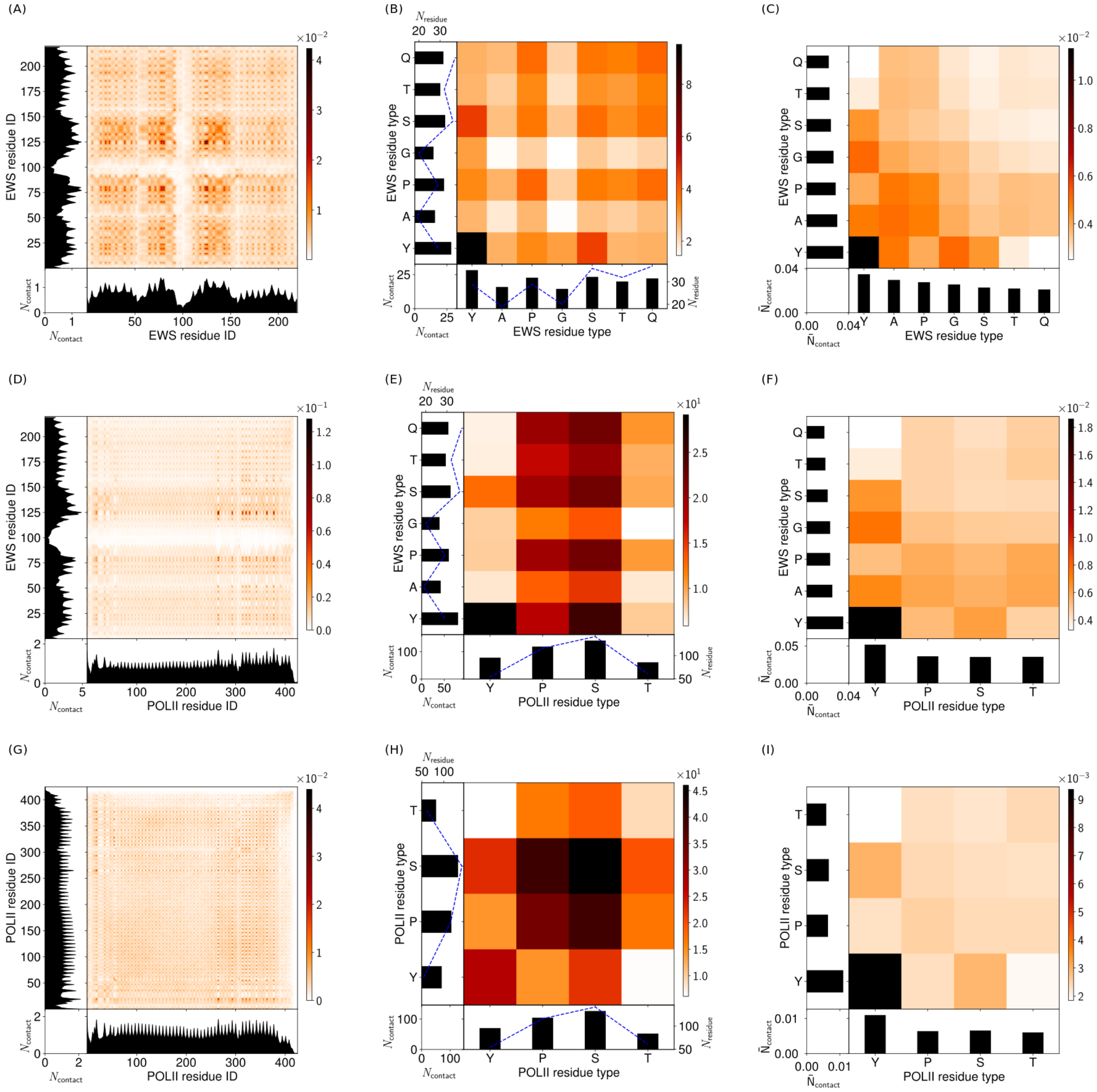

Figure S44: Molecule contact maps for a droplet simulation with 60 EWS molecules and 30 POLII molecules at 150 mM and 300 K. (A) Intermolecular contact map by particle index for EWS with other EWS. (B) Intermolecular contact map by particle type for EWS with other EWS. (C) Intermolecular contact map by particle type, normalised by particle abundance for EWS with other EWS. (D) Intermolecular contact map by particle index for EWS with POLII. (E) Intermolecular contact map by particle type for EWS with POLII. (F) Intermolecular contact map by particle type, normalised by particle abundance for EWS with POLII. (G) Intermolecular contact map by particle index for POLII with other POLII. (H) Intermolecular contact map by particle type for POLII with other POLII. (I) Intermolecular contact map by particle type, normalised by particle abundance for POLII with other POLII. The contacts in the contact maps by particle index (left column) are normalised by the number of frames used (600) and $(N_{mol1}N_{mol2})^{0.5}$ where $N_{mol1}$, $N_{mol2}$ are the number of copies of molecule 1 and molecule 2 in the simulation. The contact maps aggregated by particle type (centre column) are a matrix reduction of the contact map by particle index. Bead abundance in a molecule is shown by a blue dashed line. The contact maps (right column) are normalised by the square root of the relative bead abundance in the molecules.

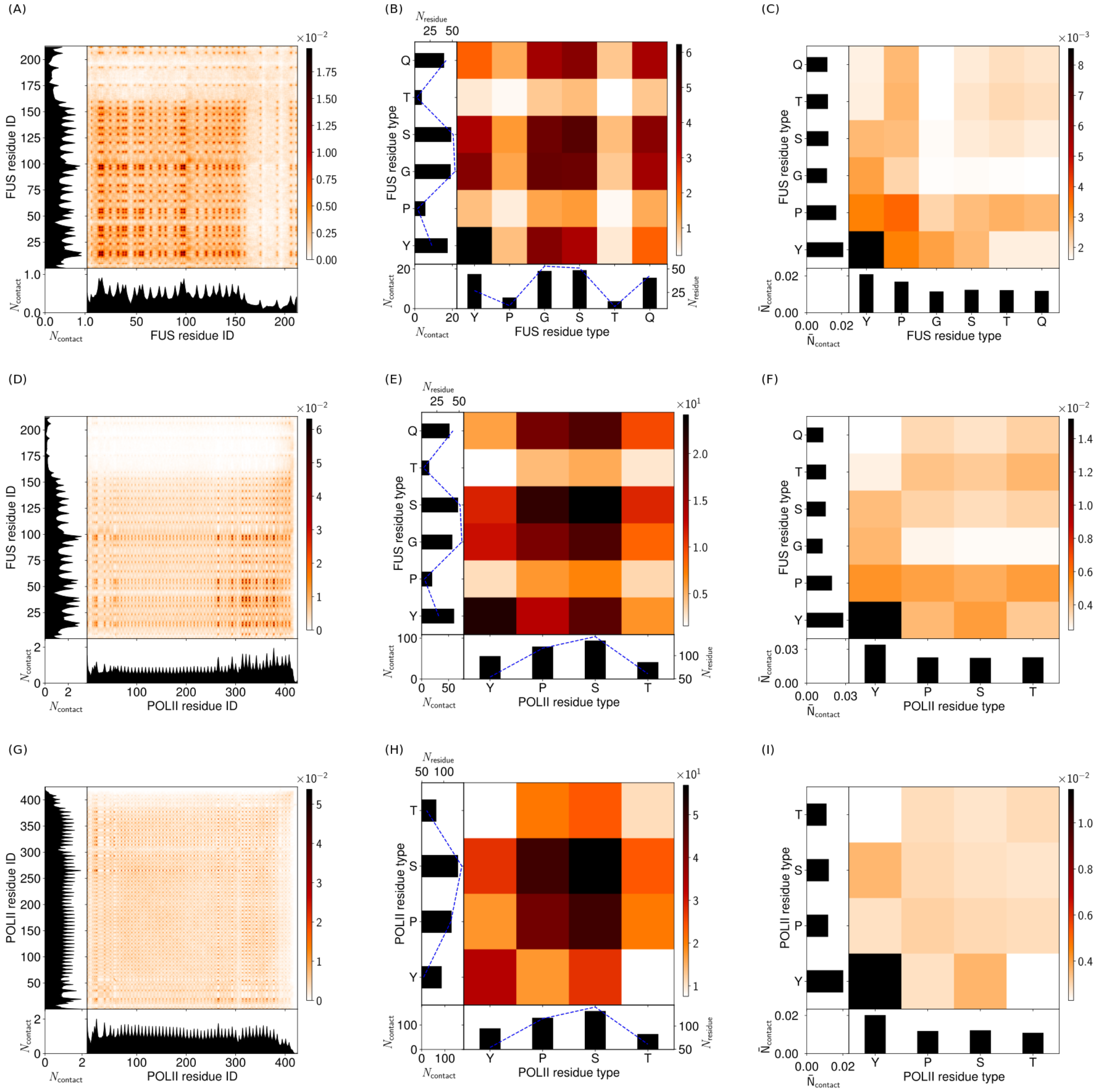

Figure S45: Molecule contact maps for a droplet simulation with 60 FUS molecules and 30 POLII molecules at 150 mM and 300 K. (A) Intermolecular contact map by particle index for FUS with other FUS. (B) Intermolecular contact map by particle type for FUS with other FUS. (C) Intermolecular contact map by particle type, normalised by particle abundance for FUS with other FUS. (D) Intermolecular contact map by particle index for FUS with POLII. (E) Intermolecular contact map by particle type for FUS with POLII. (F) Intermolecular contact map by particle type, normalised by particle abundance for FUS with POLII. (G) Intermolecular contact map by particle index for POLII with other POLII. (H) Intermolecular contact map by particle type for POLII with other POLII. (I) Intermolecular contact map by particle type, normalised by particle abundance for POLII with other POLII. The contacts in the contact maps by particle index (left column) are normalised by the number of frames used (600) and $(N_{mol1}N_{mol2})^{0.5}$ where $N_{mol1}$, $N_{mol2}$ are the number of copies of molecule 1 and molecule 2 in the simulation. The contact maps aggregated by particle type (centre column) are a matrix reduction of the contact map by particle index. Bead abundance in a molecule is shown by a blue dashed line. The contact maps (right column) are normalised by the square root of the relative bead abundance in the molecules.

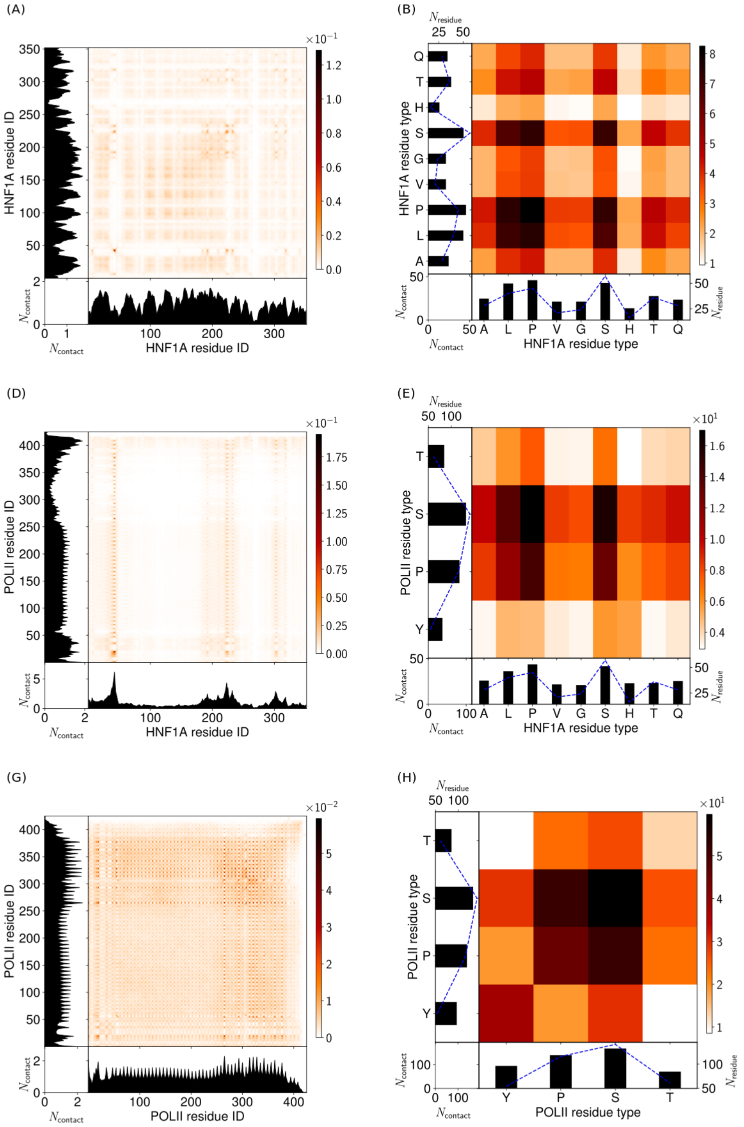

Figure S46: Molecule contact maps for a droplet simulation with 45 HNF1A molecules and 30 POLII molecules at 150 mM and 300 K. (A) Intermolecular contact map by particle index for HNF1A with other HNF1A. (B) Intermolecular contact map by particle type for HNF1A with other HNF1A. (C) Intermolecular contact map by particle type, normalised by particle abundance for HNF1A with other HNF1A. (D) Intermolecular contact map by particle index for HNF1A with POLII. (E) Intermolecular contact map by particle type for HNF1A with POLII. (F) Intermolecular contact map by particle type, normalised by particle abundance for HNF1A with POLII. (G) Intermolecular contact map by particle index for POLII with other POLII. (H) Intermolecular contact map by particle type for POLII with other POLII. (I) Intermolecular contact map by particle type, normalised by particle abundance for POLII with other POLII. The contacts in the contact maps by particle index (left column) are normalised by the number of frames used (600) and $(N_{mol1}N_{mol2})^{0.5}$ where $N_{mol1}$, $N_{mol2}$ are the number of copies of molecule 1 and molecule 2 in the simulation. The contact maps aggregated by particle type (centre column) are a matrix reduction of the contact map by particle index. Bead abundance in a molecule is shown by a blue dashed line. The contact maps (right column) are normalised by the square root of the relative bead abundance in the molecules.

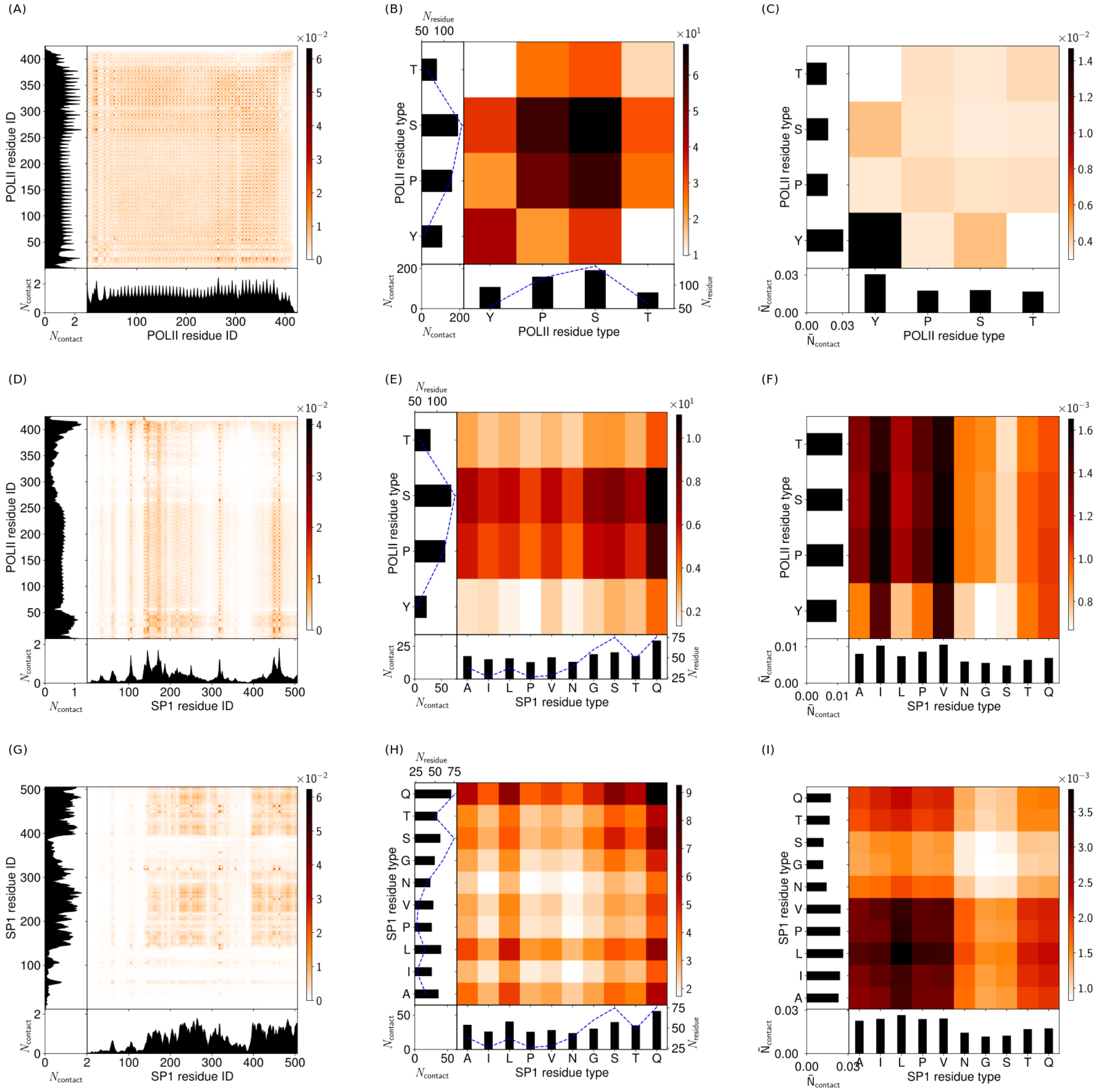

Figure S47: Molecule contact maps for a droplet simulation with 30 POLII molecules and 30 SP1 molecules at 150 mM and 300 K. (A) Intermolecular contact map by particle index for POLII with other POLII. (B) Intermolecular contact map by particle type for POLII with other POLII. (C) Intermolecular contact map by particle type, normalised by particle abundance for POLII with other POLII. (D) Intermolecular contact map by particle index for POLII with SP1. (E) Intermolecular contact map by particle type for POLII with SP1. (F) Intermolecular contact map by particle type, normalised by particle abundance for POLII with SP1. (G) Intermolecular contact map by particle index for SP1 with other SP1. (H) Intermolecular contact map by particle type for SP1 with other SP1. (I) Intermolecular contact map by particle type, normalised by particle abundance for SP1 with other SP1. The contacts in the contact maps by particle index (left column) are normalised by the number of frames used (600) and $(N_{mol1}N_{mol2})^{0.5}$ where $N_{mol1}$, $N_{mol2}$ are the number of copies of molecule 1 and molecule 2 in the simulation. The contact maps aggregated by particle type (centre column) are a matrix reduction of the contact map by particle index. Bead abundance in a molecule is shown by a blue dashed line. The contact maps (right column) are normalised by the square root of the relative bead abundance in the molecules.

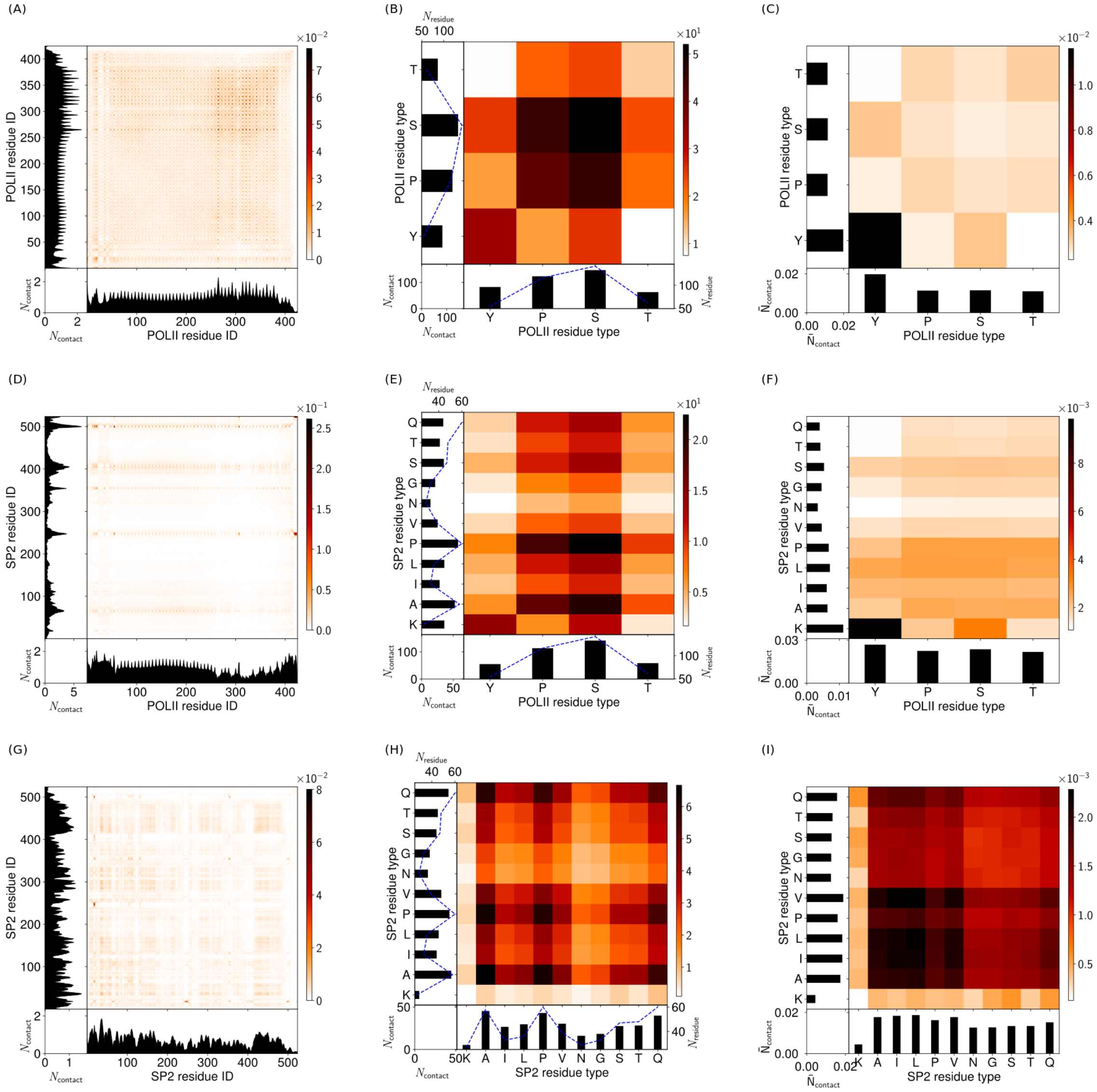

Figure S48: Molecule contact maps for a droplet simulation with 30 POLII molecules and 30 SP2 molecules at 150 mM and 300 K. (A) Intermolecular contact map by particle index for POLII with other POLII. (B) Intermolecular contact map by particle type for POLII with other POLII. (C) Intermolecular contact map by particle type, normalised by particle abundance for POLII with other POLII. (D) Intermolecular contact map by particle index for POLII with SP2. (E) Intermolecular contact map by particle type for POLII with SP2. (F) Intermolecular contact map by particle type, normalised by particle abundance for POLII with SP2. (G) Intermolecular contact map by particle index for SP2 with other SP2. (H) Intermolecular contact map by particle type for SP2 with other SP2. (I) Intermolecular contact map by particle type, normalised by particle abundance for SP2 with other SP2. The contacts in the contact maps by particle index (left column) are normalised by the number of frames used (600) and $(N_{mol1}N_{mol2})^{0.5}$ where $N_{mol1}$, $N_{mol2}$ are the number of copies of molecule 1 and molecule 2 in the simulation. The contact maps aggregated by particle type (centre column) are a matrix reduction of the contact map by particle index. Bead abundance in a molecule is shown by a blue dashed line. The contact maps (right column) are normalised by the square root of the relative bead abundance in the molecules.

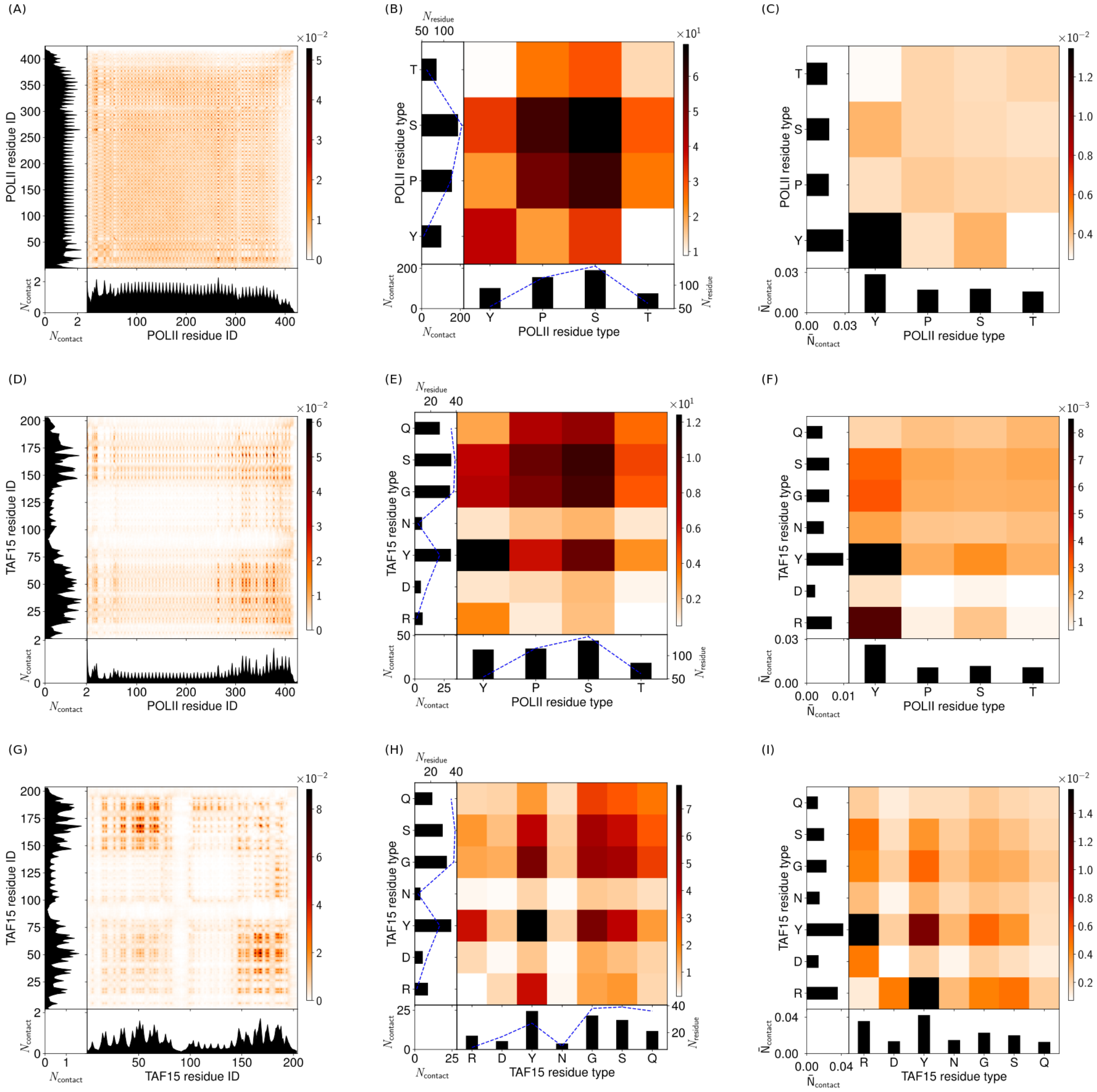

Figure S49: Molecule contact maps for a droplet simulation with 30 POLII molecules and 60 TAF15 molecules at 150 mM and 300 K. (A) Intermolecular contact map by particle index for POLII with other POLII. (B) Intermolecular contact map by particle type for POLII with other POLII. (C) Intermolecular contact map by particle type, normalised by particle abundance for POLII with other POLII. (D) Intermolecular contact map by particle index for POLII with TAF15. (E) Intermolecular contact map by particle type for POLII with TAF15. (F) Intermolecular contact map by particle type, normalised by particle abundance for POLII with TAF15. (G) Intermolecular contact map by particle index for TAF15 with other TAF15. (H) Intermolecular contact map by particle type for TAF15 with other TAF15. (I) Intermolecular contact map by particle type, normalised by particle abundance for TAF15 with other TAF15. The contacts in the contact maps by particle index (left column) are normalised by the number of frames used (600) and $(N_{mol1}N_{mol2})^{0.5}$ where $N_{mol1}$, $N_{mol2}$ are the number of copies of molecule 1 and molecule 2 in the simulation. The contact maps aggregated by particle type (centre column) are a matrix reduction of the contact map by particle index. Bead abundance in a molecule is shown by a blue dashed line. The contact maps (right column) are normalised by the square root of the relative bead abundance in the molecules.
